# Geometry Linked to Untangling Efficiency Reveals Structure and Computation in Neural Populations

**DOI:** 10.1101/2024.02.26.582157

**Authors:** Chi-Ning Chou, Royoung Kim, Luke A. Arend, Yao-Yuan Yang, Brett D. Mensh, Won Mok Shim, Matthew G. Perich, SueYeon Chung

**Author notes:** These authors contributed equally as second authors (Royoung Kim, Luke A. Arend, Yao-Yuan Yang).

## Abstract

From an eagle spotting a fish in shimmering water to a scientist extracting patterns from noisy data, many cognitive tasks require untangling overlapping signals. Neural circuits achieve this by transforming complex sensory inputs into distinct, separable representations that guide behavior. Data-visualization techniques convey the geometry of these transformations, and decoding approaches quantify performance efficiency. However, we lack a framework for linking these two key aspects. Here we address this gap by introducing a data-driven analysis framework, which we call Geometry Linked to Untangling Efficiency (GLUE) with manifold capacity theory, that links changes in the geometrical properties of neural activity patterns to representational untangling at the computational level. We applied GLUE to over seven neuroscience datasets—spanning multiple organisms, tasks, and recording techniques—and found that task-relevant representations untangle in many domains, including along the cortical hierarchy, through learning, and over the course of intrinsic neural dynamics. Furthermore, GLUE can characterize the underlying geometric mechanisms of representational untangling, and explain how it facilitates efficient and robust computation. Beyond neuroscience, GLUE provides a powerful framework for quantifying information organization in data-intensive fields such as structural genomics and interpretable AI, where analyzing high-dimensional representations remains a fundamental challenge.

---

An animal’s survival depends on its ability to swiftly identify objects in its environment and respond appropriately. Consider a cat detecting movement in nearby bushes: it must rapidly determine whether the source is a predator (like a snake) or prey (like a mouse) and respond accordingly (e.g., fleeing or stalking, Figure 1a, left). Encounters with snakes under varying lighting conditions and different levels of visual clutter lead to a wide variety of retinal activity patterns. Geometrically, these patterns form continuous, low-dimensional, curved surfaces known as *category manifolds* [1] within the space of neural responses (Figure 1a, middle). Distinct objects, such as snakes and mice, form manifolds within a population activity space (e.g., a brain area) at a given time. However, the manifolds of many different objects can be *tangled* within the activity space of sensory neurons—for example, under certain conditions, a partially obscured mouse with a long, curly tail may resemble a snake, in which case their category manifolds may be highly proximal or even overlap (Figure 1a, middle). Despite such complex and continuous variability in the stimulus space, the visual system has evolved the remarkable ability to distinguish and recognize objects through a cascade of neural transformations that progressively *untangle* [2] these category manifolds into distinct percepts (Figure 1a, right, dotted lines), thereby enabling the animal to take appropriate behavioral action, such as fleeing or stalking (Figure 1a, right).

**Figure 1:**
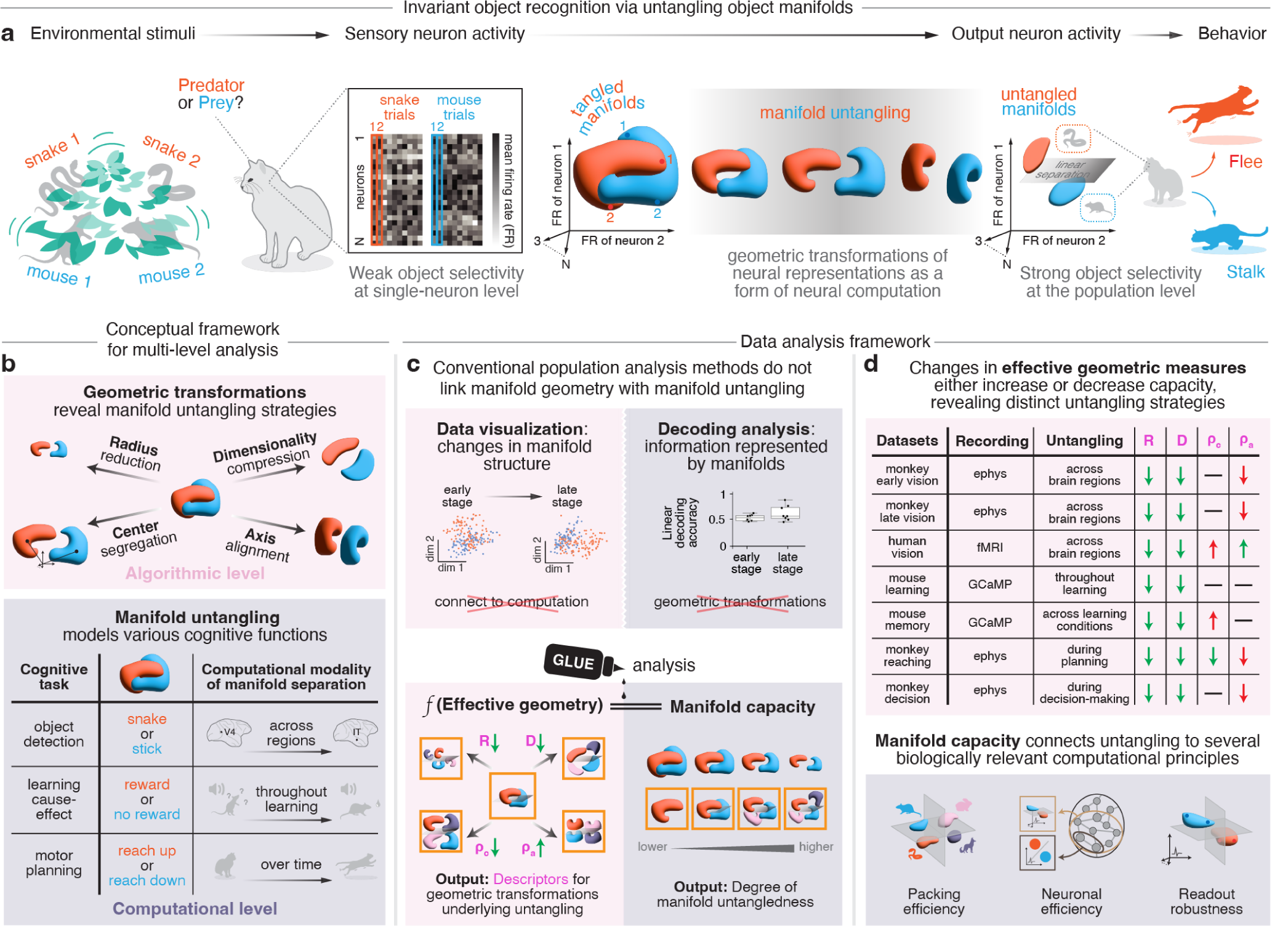
Neural manifold untangling and our proposed analysis framework, Geometry Linked to Untangling Efficiency via manifold capacity theory (GLUE). **a**. A schematic illustration of manifold untangling in the context of invariant object recognition. When a cat sees something moving behind a bush, it must quickly decide—based on the object’s category (e.g., predator or prey)—whether to flee or stalk, a decision crucial to its survival. Different instances of environmental stimuli from the same object category (e.g., snakes with different postures) induce different patterns of neural activity in the sensory area (e.g., retina, visual cortex), forming an object manifold in that area for the category. Because the cat can easily distinguish prey from predator and execute the corresponding behavior, this suggests that the neural manifolds of these categories are well separated from each other in the neural activity space just before the output neurons (e.g., motor neurons). This separation allows for fast and robust actions despite the high-dimensional variabilities at the sensory end. This process, known as invariant object recognition, is proposed to occur through a sequence of untangling these object manifolds. **b**-**d**. Our proposed conceptual and data analysis framework for studying cognitive untangling at both the structural level (pink) and the computational level (purple) in neural populations. **b**. A conceptual framework for studying cognitive brain functions by modeling the underlying geometric transformations that drive the untangling of task-relevant neural manifolds. Top: Changes in manifold geometry can reveal the differences in manifold untangling as there might be distinct geometric transformations for manifold untangling due to factors such as stimuli, tasks, and neural circuits. Bottom: Manifold untangling can model many cognitive brain functions, including visual recognition (Figure 3), decision-making (Figure 4), and motor control (Figure 5). **c**. Top: Conventional analyses such as visualization method (top-left) and decoding method (top-right) are able to study the organizational changes and task-relevant information of manifold untangling respectively. However, there lacks a framework to connect these two levels of abstraction. Bottom: Our proposed analysis framework, GLUE, enables data analyses for manifold untangling through an analytical link between the geometrical properties of manifolds (left, measured by effective manifold geometric measures) and manifold untangling (right, measured by manifold capacity). Specifically, in our theory, manifold capacity is a function of effective geometric measures (such as effective radius and dimension, see Supplementary Information S1). The orange boxes correspond to the ribbon metaphor, illustrating how the shape and organization of manifolds (in a maximally packed configuration) connects to the number of packable manifolds. **d**. GLUE enables the conceptual framework described in panel b by offering a data-driven analysis framework. Top: In GLUE, one can study the distinct geometric transformations of manifold untangling across different neural systems by analyzing the changes in effective geometric measures of task-relevant manifolds. The gray boxes are summaries of how each effective geometric measure changed from before to after untangling. See later sections and figures for quantitative versions of the results. Bottom: In GLUE, manifold capacity connects manifold untangling to three biologically relevant computational principles (Method M3.4), thereby allowing for a wide range of interpretations for manifold untangling in data analysis. See Figure 6 for more details.

Viewing cognitive brain functions—such as the abovementioned invariant object recognition—as processes of untangling task-relevant neural representations (category manifolds) offers a systematic way to investigate their neural underpinning. To think about how the organization of category manifolds are related to the process of untangling them, imagine arranging ribbons in a storage box: if the ribbons are jumbled without clear partitions, picking up a specific piece with a tweezer becomes hard and error-prone. Similarly, the population activity space defined by *N* recording units can be thought of as an *N*-dimensional storage box, and category manifolds as the ribbons inside. Tangled category manifolds occupy more space and impede the brain’s ability to swiftly distinguish between different representations to determine the appropriate percept, thereby disrupting behavior. To untangle the ribbons, one might compress them into small balls, align them in stacks, or use some intermediate method; the most efficient mechanism to untangle a set of ribbons will depend on their geometrical properties, such as their size or alignment. Likewise, the geometrical properties of category manifolds—shaped by factors such as stimulus characteristics, evolutionary pressures acting on the neural system, and/or the intrinsic properties of neural circuits—affect how they can be untangled. Depending on these geometrical properties and the neural system involved, different *geometric transformations*—methods for untangling manifolds based on their geometrical properties—may be at play (Figure 1b, top). This framework of untangling task-relevant category manifolds through geometric transformations provides a broadly applicable analytical language for modeling how the brain solves various cognitive untangling challenges (Figure 1b, bottom).

To determine whether cognitive untangling is in fact achieved in the brain via geometric transformation-based manifold untangling, we need to be able to discern the geometrical properties of category manifolds from within large sets of neural activity data. Current approaches, however, cannot accommodate this type of data analysis. For example, single-neuron tuning analyses are not optimized to appreciate broader, population-based sensory perceptions, which present as mixed selectivity in higher-order brain areas [3–5]. Moreover, while existing population-level analysis methods may be able to examine changes in neural activity patterns or detect the presence of manifold untangling processes, none can connect these two aspects (Figure 1c, top). For instance, dimensionality reduction techniques [6–8] provide valuable geometric intuitions by visualizing high-dimensional data and revealing the underlying spatial organization of neural activity patterns in latent spaces. However, in prioritizing some variables over others they may discard information that would be critical to detect how the category manifolds themselves are untangled to direct task-related behavior (Figure 1c, top left). In contrast, decoding methods [2, 9, 10] quantify task-relevant information but offer limited insight into underlying neural mechanisms. Similarly, approaches linking information to neural correlations [11, 12] rely primarily on low-order statistics, limiting their effectiveness for neural populations with complex tuning properties or mixed selectivity (Figure 1c, top right).

Here we address these limitations and gaps in existing analysis methods by presenting a novel analysis framework, called **Geometry Linked to Untangling Efficiency via manifold capacity theory (GLUE)**. GLUE builds on a rich body of theoretical work on *perceptron capacity* [13–19] and its recent extension to category manifolds, known as *manifold capacity* [20, 21]. We advanced the mathematical theory by refining previous assumptions, which often fail in biological data. Our theory revealed geometrical properties of category manifolds that can be used to determine the degree to which those manifolds are untangled (Figure 1c, bottom). GLUE makes it possible to identify the geometric transformations driving manifold untangling across a wide range of neuroscience datasets. The result is a first-of-its-kind framework that can be used to systematically probe the computational strategies underlying brain functions that involve the transformation of overlapping input signals into discrete outcomes, such as visual discrimination, decision-making, or motor control.

To understand how GLUE helps us study manifold untangling, consider the ribbon metaphor again: One can assess the packing efficiency of ribbons in a box by calculating the maximum number of ribbons per unit volume of storage that still allows for easy retrieval. Similarly, using GLUE we can quantify the degree of manifold untangledness by calculating **manifold capacity**—defined as the maximum load of category manifolds in the neural activity space that still allows for easy discrimination by downstream processing (Figure 1c, bottom right; Supplementary Information S1). Just as there are various ways to pack ribbons efficiently, there are multiple geometric arrangements of category manifolds that can achieve high manifold capacity (Figure 1b, top). Using GLUE, we are able to derive the **effective manifold geometric measures**^1^ of category manifolds that contribute to manifold capacity, which can in turn be used to quantify the degree of manifold untangling (Figure 1c, bottom). Just as packing more ribbons into a box is easier when they are compact, thin, and/or evenly distributed, higher manifold capacity can be achieved by reducing manifold **radius**, compressing manifold **dimensionality**, and/or optimizing organization, i.e., increasing manifold **center segregation** or enhancing manifold **axis alignment** (Figure 1b top).

Here, we present GLUE as a novel conceptual and data analysis framework for studying distinct mechanisms of manifold untangling across different neural systems through their underlying geometric transformations (Figure 1b-1d). In addition, we apply GLUE to a wide range of neural data involving multiple organisms (human, monkey, mouse), task modalities (vision, decision-making, motor), and recording techniques (electrophysiology, calcium imaging, fMRI), as well as artificial neural networks, to track changes in manifold capacity and geometric measures. From there, we are able to discern new computational principles underlying complex tasks (Figure 6). For example, how the idea of manifold packing efficiency may apply to object recognition, or how decoding efficiency (the ability to signal using relatively few neurons) may apply to stimulus discrimination, or how readout robustness (the alignment of neural activity across many neurons in support of a particular outcome) may apply to motor planning. The strength of our approach is in the ability to link structural properties within high-dimensional representations to their relevant computational/functional roles. Beyond neuroscience, GLUE is likely to advance other data-intensive fields like structural genomics or interpretable AI by addressing a fundamental challenge in complex systems—how to transform distributed representations into categorical patterns.

## Results

### Revealing structure in high-dimensional data by linking the geometry of neural activity patterns to perceptron capacity

A common approach to assessing the accuracy and efficiency of information decoding from neural representations is to model the readout operation of a downstream neuron as a linear classifier^2^, also known as a perceptron [22, 23]. The ratio between the number of linearly discriminable patterns and the size of a neural state space (often quantified by the number of neurons) is then known as perceptron capacity [13–15]. This approach has inspired decades of rich theoretical work [16–19]. However, the original concept of perceptron capacity has a significant limitation: it considers discrete points as the counting unit for patterns. This assumption does not apply to biological data. Indeed, in most real-life scenarios, robust object discrimination involves separating representations with smooth nuisance variations (e.g., changes in the input stimuli that are invariant to the animal’s perception/behavior), making this classical notion of capacity irrelevant.

To better address real-world scenarios, the classical notion of perceptron capacity for discrete points has recently been generalized to encompass the capacity for *category manifolds* [20, 21]. For example, consider a task with two conditions (e.g., the prey and predator object category as in Figure 1a) and neural recordings with *N* units in a certain brain area. The population neural response from a single trial corresponds to a vector in the *N*-dimensional neural activity space. Grouping the vectors associated with the same condition forms two point clouds, each representing a condition, or “object”, that may need to be discriminated by downstream processing. Numerically, the capacity of these two point cloud manifolds can be estimated by 2*/N* ^*^, where *N* ^*^ is the minimum number of neurons required to ensure that the two point clouds are linearly separable (see Method M3.2). For example, tangled data point clouds, such as the classical XOR problem [23] (Figure 1a middle), require a high-dimensional embedding space for linear separation (i.e., larger *N* ^*^), necessitating a larger number of neurons and thereby reducing capacity [24, 15].

Within the context of neural computation, the process of manifold untangling may involve distinct mechanisms depending on stimuli, tasks, neural circuits, or other factors (Figure 1a) which, as discussed above, define the geometrical properties of the manifold. Characterizing changes in the geometric properties of category manifolds is an intuitive and quantitative way to study the untangling process. Recent theoretical work [20, 21] has employed statistical physics models to derive data-driven formulas for approximating capacity values (see Supplementary Information S1). Crucially, these formulas reveal geometric properties of high-dimensional manifolds that affect capacity, thereby opening a new avenue to investigate the underlying untangling mechanism by tracking changes in manifold geometry. Applications of these early theories on manifold capacity have shown promising results in interpreting high-dimensional data in both neuroscience [25–27] and artificial neural networks [28–32]. However, previous theoretical work [20, 21] relies on two critical simplifying assumptions—Gaussian correlations between manifolds and random task labels—which often fail in biological datasets by ignoring higher-order correlations and task-specific structures, thereby limiting their applicability (see Figure 2 and Supplementary Figure S3).

**Figure 2:**
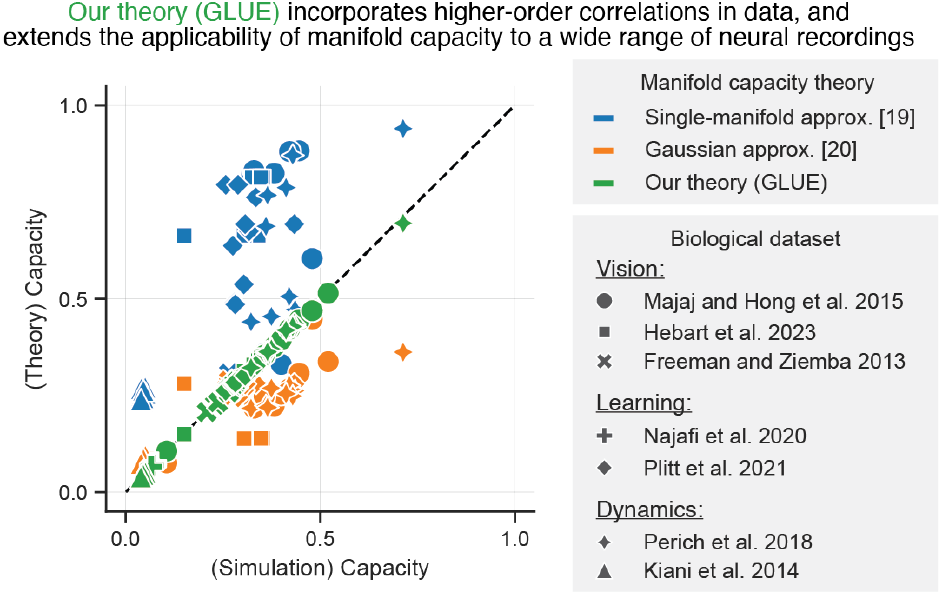
Theoretical advances enable capacity-based analyses. The x-axis is the (ground truth) capacity value estimated from numerical simulation (Method M3.4) and the y-axis is the capacity value estimated from a manifold capacity formula. Each point corresponds to the result of a random (e.g., a session, an animal) analysis run from that dataset. Our theory significantly improves the theory-simulation match in neural recordings from earlier theory [20, 21], hence enabling capacity-based analyses to a broad range of neural datasets for the first time. See Supplementary Figure S3 for a detailed comparison.

### Using the GLUE theory, we can mathematically predict numerical estimations of perceptron capacity based on high-dimensional biological data

We eliminated all the mathematical assumptions inherent in previous theoretical approaches. This enabled—for the first time—the application of capacity-based analysis to a broad range of neural data. We achieved this by incorporating interdisciplinary ideas from integral geometry and convex optimization theory to advance the theoretical foundation of capacity theory. Specifically, we derived a new manifold capacity formula, which (1) achieves perfect agreement with the empirical estimation (Figure 2 and Supplementary Figure S3) and (2) connects directly to the organization of neural activity patterns and task structure. Here, we offer a glimpse of our theoretical results using the two point-cloud manifold example described above. Suppose we have *M* trials per condition and 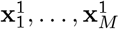 and 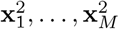 as the population neural activity vectors for the two conditions respectively. Then, we derive the following capacity formula:

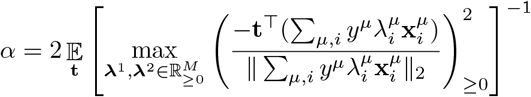

where 𝔼_**t**_ denotes taking the expectation over **t** being sampled from the *N*-dimension isotropic Gaussian distribution, and *y*^1^ = 1, *y*^2^ = − 1 are the labels for the two point cloud manifolds respectively. The critical advance of our theory is that the capacity (*α*) can now be expressed as a closed-form formula directly dependent on the data 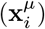 and a specific choice of task label (*y*^*µ*^), whereas the formulas from previous work rely on assumptions of random directions [20], pre-processing [28], or transformations [21] of the data and task labels. See Supplementary Information S1 for the full version of our theory and see Figure 2 and Supplementary Figure S3) for the theory-numerics match.

#### Using the GLUE theory, we can extract geometric features of neural activation patterns that reveal the mechanistic basis of manifold capacity

The mathematical formula for manifold capacity in our theory not only achieves perfect estimation accuracy of capacity value (Figure 2), but also enables us to extract geometric features of neural activation patterns that are analytically connected to the capacity value through the formula. We introduced a set of effective geometric measures that incorporate complex shapes and correlations of category manifolds (e.g., beyond second order statistics) and are analytically linked with manifold capacity through our new formula (Method M3 and Supplementary Information S1). In particular, several geometric transformations can lead to an increase in manifold capacity: reducing manifold radius *R*, compressing manifold dimensionality *D*, segregating manifold centers *ρ*_*c*_, and aligning manifold axis *ρ*_*a*_ (Figure 1b top, Supplementary Information S1.4). For example, manifold capacity can be approximated by a simple function of the geometric measures, e.g., *α* ≈ (1 + *R*^−2^)*/D* (Supplementary Information S1.4).

In addition, we established equivalences between manifold capacity and several normative principles of computation(Figure 1c, bottom). This in turn enables new biologically-relevant interpretations and broadens the applicability of our theoretical framework to more diverse datasets. Specifically, our theoretical result shows that an increase in manifold capacity corresponds to an increase in packing efficiency (being able to pack more manifolds, Figure 6a), an increase in neuronal efficiency (requiring fewer neurons for decoding, Figure 6b), and an increase in readout robustness (Figure 6c).

In summary, our theoretical framework quantifies manifold untangling via manifold capacity, linking it to the shapes and organization of neural representations and, for the first time, enabling capacity-based methods to be applied to a wide range of neuroscience problems.

#### Applying GLUE to a wider range of neural data

The test of GLUE as a theoretical framework applicable to biologically-relevant neural computations is whether it can be generalized to various cognitive tasks. For instance, while it is relatively intuitive to imagine object detection as a process of untangling two or more category manifolds (snake manifold versus stick manifold), it is less intuitive to think about the process of learning or motor planning in this way. Here, we present evidence of the potential to use GLUE to reveal the role of manifold untangling within three different cognitive modalities: across brain regions (as is the basis of visual object detection), over the course of learning (before/after learning that a tone signals reward), and over time (before initiating a movement, after initiation, during movement).

### Using GLUE to reveal the degree of untangling and the geometric transformations involved in computations occurring across brain regions

Having established the theoretical framework, we next asked whether GLUE could reveal how the brain transforms complex sensory inputs into categorical percepts. The ventral visual stream offers an ideal test case, as it must convert variable retinal inputs into stable object representations. Using GLUE analysis, we set out to determine how brain regions along the ventral visual stream encode object categories, i.e., manifolds comprising collections of neural activity patterns in a sensory area (e.g., IT) associated with a specific object category (e.g., cat). We tested the hypothesis that these object categories become increasingly untangled as information flows through the visual system hierarchy, supporting downstream tasks like invariant object recognition.

To test this hypothesis, we analyzed three datasets comprising recordings from various stages of the visual hierarchy: a monkey electrophysiology dataset [35] with texture pattern category manifolds in the early ventral stream (Supplementary Figure S8), a monkey electrophysiology dataset [33] with object category manifolds in the late ventral stream (Figure 3a–c), and a human fMRI dataset [34] with subordinate-level category manifolds in the ventral temporal (VT) cortex (Figure 3e–h). Our study revealed three new findings, which are described in detail in the following section. Briefly, we find that, (1) across all three datasets, object category manifolds progressively untangle along the visual hierarchy, as indicated by increased manifold capacity; (2) the untangling processes shared common geometric transformations—a reduction in manifold radius and a compression of manifold dimension; and (3) distinct geometric transformations emerged that varied with processing stage and input category type, as evidenced by differences in alignment measures.

**Figure 3:**
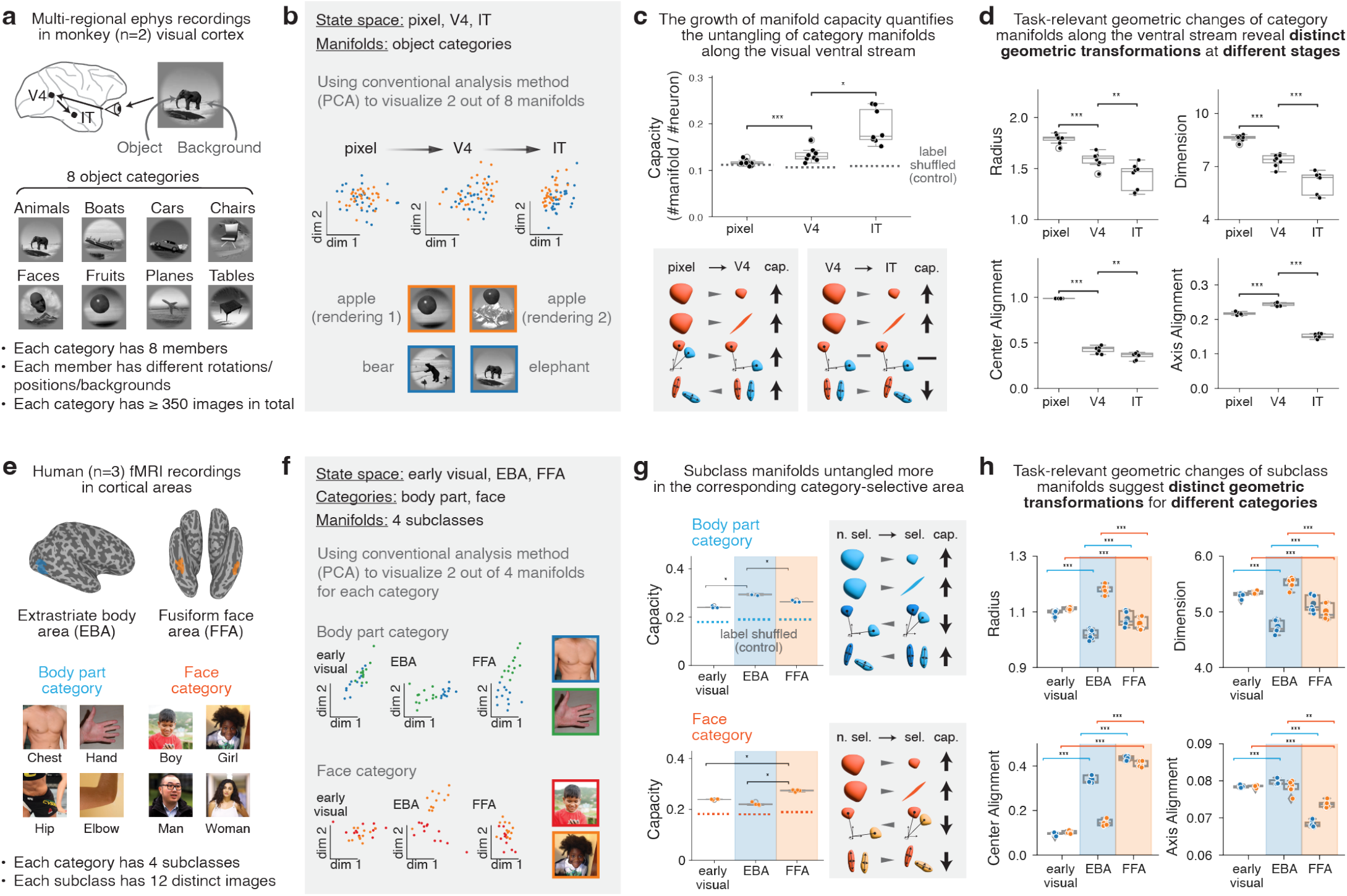
GLUE reveals the underlying geometric transformations of the untangling of object manifolds along the visual hierarchy. **a**. A dataset from [33] consisting of multi-electrode recordings in monkeys’ (n=2) V4 and IT. The monkeys were presented with a sequence of images, where each image contained a 2D rendering of a 3D object superimposed on a natural scene background. There were 8 basic-level object categories, and each category included 8 subordinate objects. **b**. For a brain area (V4 or IT) or the image pixel space, we define 8 basic-level object manifolds. An object manifold contains the (preprocessed) neural responses (or pixel vector for the image pixel space) to all stimuli from the same object class (at least 350 images per category). **c**. Manifold capacity analysis. Top: capacity of object manifolds and capacity of the shuffled manifolds. A data point corresponds to one of the possible 1-vs-rest dichotomies of classifying 8 object categories (Mann–Whitney U test; *: *p <* 0.05; **: *p <* 0.01; ***: *p <* 0.001). Bottom: a schematic summary of the geometric transformations of object manifolds and their relation to capacity. **d**. Geometric transformations of object manifolds from pixel level to V4 and IT (Mann–Whitney U test; *: *p <* 0.05; **: *p <* 0.01; ***: *p <* 0.001). **e**. A dataset from [34] consisting of fMRI data in human subjects’ (n=3) cortical areas, such as extrastriate body area (EBA) and fusiform face area (FFA). The brain surface is the native brain surface of the example subject (sub-02). Participants viewed object images while performing an oddball detection task, responding to occasional artificially generated images. We selected 4 body part subordinate categories and 4 face subclasses, where each contains 12 object images. **f**. For a region of interest (e.g., EBA or FFA) and an object category (body part or face), we define 4 subordinate category manifolds. A subordinate category manifold is the collection of single-trial beta coefficients estimated from BOLD responses (with the number of voxels randomly subsampled to 400) corresponding to the object images in that subordinate category. **g**. Manifold capacity analysis. Top-left: capacity of body part manifolds. Top-right: a schematic summary of the geometric transformations of body part manifolds and their relation to capacity. Bottom-left: capacity of face manifolds. Bottom-right: a schematic summary of the geometric transformations of face manifolds and their relation to capacity. A data point corresponds to one of the possible 1-vs-rest dichotomies of classifying 4 subclasses (Mann–Whitney U test; *: *p <* 0.05; **: *p <* 0.01; ***: *p <* 0.001). **h**. Geometric transformations of subordinate manifolds along the ventral visual stream (Mann–Whitney U test; *: *p <* 0.05; **: *p <* 0.01; ***: *p <* 0.001).

#### Object category manifolds within neural population activation data are untangled via compression and decorrelation along the primate ventral stream

We considered a dataset [33] with simultaneous electrophysiology recordings on macaque monkeys (n=2) from the V4 and inferior temporal (IT) cortices (126 sites in V4 and 189 sites in IT). For these recordings, head-fixed monkeys were presented with a sequence of images for 100ms each, with 100ms of blank screen in between. Each image contained a 2D rendering of a 3D object superimposed on a natural scene background. There were eight object categories (a.k.a., basic-level object categories in [33], e.g., animals, boats, cars, chairs, faces, fruits, planes, tables), and each category included eight members (a.k.a., subordinate-level objects, i.e., 64 objects in total) (Figure 3a). For each basic-level category, we defined the category manifold as the collection of neural responses (firing rate vectors) in a particular brain area to images from that category (at least 350 images per category). Thus, a category manifold encom-passed neural responses to different objects within the category, varying view parameters (e.g., position, scale, pose) of the objects, and different background scenes, all sharing the same category label (Figure 3b).

When we apply GLUE analysis, we find that category manifold capacity increases from the image pixel space to V4 to IT (Figure 3c, top). To confirm that these structural changes were specific to categorical representations rather than global changes, we performed the same analysis on label-shuffled manifolds and observed no increase in capacity (Figure 3c, top). Similarly, in a dataset featuring texture family categories [35], we observed an increase in manifold capacity from V1 to V2 (Supplementary Information S3.1). These findings support the hypothesis that category manifold untangling occurs along the ventral stream.

We next investigated the geometric transformations of this untangling process. Our analyses reveal that both manifold radius and dimension decrease from pixel space to V4 and from V4 to IT (Figure 3d, right). On the other hand, changes in axis alignment delineate the differences between V4 and IT. Thus, manifold centers become more segregated from V4 to IT—improving capacity—while manifold axes become less aligned—reducing capacity (Figure 3d). These findings reveal shared as well as distinct geometric transformations for category manifold untangling at different stages of processing in the visual ventral stream, raising intriguing questions for future investigations into the underlying neurophysiology of the computational task handled by this pathway (see Supplementary Information S3.2 for more discussion and interpretation).).

#### Specific object subclassification manifolds from within human fMRI data are untangled via distinct geometric transformations

The improved estimation accuracy of manifold capacity by GLUE (Figure 2) overcomes the challenge of analyzing population representations in fMRI data [36, 37], enabling capacity-based analysis for the first time. For this analysis, we focused cortical regions known to be selective for particular categories, such as the fusiform face area (FFA) for faces and the extrastriate body area (EBA) for body parts. We utilized the THINGS dataset [34] comprising fMRI recordings from human participants (n=3) who viewed object images while performing an oddball detection task, responding to occasional artificially generated images. The object images were richly labeled, allowing us to define four subordinate categories (boy, girl, man, woman) for the face category and four subordinate categories (chest, hand, hip, elbow) for the body part category (Figure 3e). Each subordinate category contained 12 unique object images. For each cortical area, we defined a subordinate category manifold as the collection of single-trial beta coefficients estimated from blood oxygenation level-dependent (BOLD) responses (voxels randomly subsampled to 400) corresponding to the object images in that subordinate category (Figure 3f).

We applied GLUE analysis to test the hypothesis that subordinate category manifolds become more untangled in their corresponding selective regions. Our results showed that manifold capacity increased along the visual hierarchy, peaking in the relevant selective area: body part manifolds reached their highest capacity in EBA (Figure 3g, top-left) and face manifolds in FFA (Figure 3g, bottom-left). Furthermore, after controlling for noise differences across regions using a non-selective control category (e.g., electronic devices; see Method M1.7), we observed a consistent increase in manifold capacity from earlier regions in the visual hierarchy (Supplementary Figure S10). Together, these findings support the hypothesis that subordinate-level category manifolds become more untangled in areas known to be selective for that category.

To further elucidate the geometric transformations underlying these untangling processes, we analyzed the extent to which different effective geometric measures drive the changes in capacity across regions. Consistent with our earlier findings from the monkey ventral stream, changes in manifold radius and dimension were apparent along the computational pathway; both reached their smallest values in the corresponding category-selective regions (Figure 3h). Interestingly, changes in alignment measures as manifolds untangled were distinct in EBA and FFA: body part manifolds exhibited lower center alignment and higher axis alignment in EBA—thereby enhancing capacity—while face manifolds showed the opposite trend in FFA, which reduced capacity. These findings highlight that category manifold untangling strategy varies with stimulus category and selective area, raising intriguing questions for future investigations (see Supplementary Information S3.3 for more discussion and interpretation).

### Using GLUE to reveal the degree of untangling and the geometric transformations involved in computations related to learning

Above, we demonstrate how category manifolds are organized in mature circuits (Figure 3). However, these findings raise a crucial question: How do neural populations acquire appropriate geometrical structure as a circuit is forming, i.e., during learning?

#### Challenges in analyzing neural population representations during learning

Analyzing how neural populations represent information during learning presents a fundamental challenge due to the high dimensionality and dynamic nature of neural activity [39–41]. Conventional methods often struggle to capture how distributed neural representations evolve as new associations are formed. To do this requires tracking changes in the geometrical structure of population activity over time [5, 6], which is difficult or impossible using conventional analysis methods such as visualization or decoding analysis. GLUE has the potential to overcome this challenge because it can focus on tracking the geometric properties that are relevant for a task.

In this section, a category manifold corresponds to the collection of neural representations associated with a specific stimulus category related to a task being learned (a “stimulus category manifold”; e.g., a cue that is associated with reward versus another cue that is not). We investigated the hypothesis that the degree of stimulus category manifold untangling is related to the performance and/or the procedure of learning. For example, better behavioral accuracy may be related to more untangled manifolds, and different learning procedures may result in varying levels of manifold untangling.

We analyzed two datasets: calcium imaging from mice learning an auditory decision-making task [38] (Figure 4) where we studied if the manifold capacity increases alongside improvements in behavioral accuracy ; and calcium imaging data from mice learning an associated context-discrimination task [42] (Supplementary Figure S14) where we studied the difference in manifold capacity and geometry between mice trained under different prior experience. In summary, our GLUE analysis showed that learning may restructure neural representations, and the degree of category manifold untangling can serve as a useful task-relevant measure to quantify such changes and link to behavior. Furthermore, the geometric measures in GLUE provide a dictionary of potential geometric transformations for how such untangling occurs.

**Figure 4:**
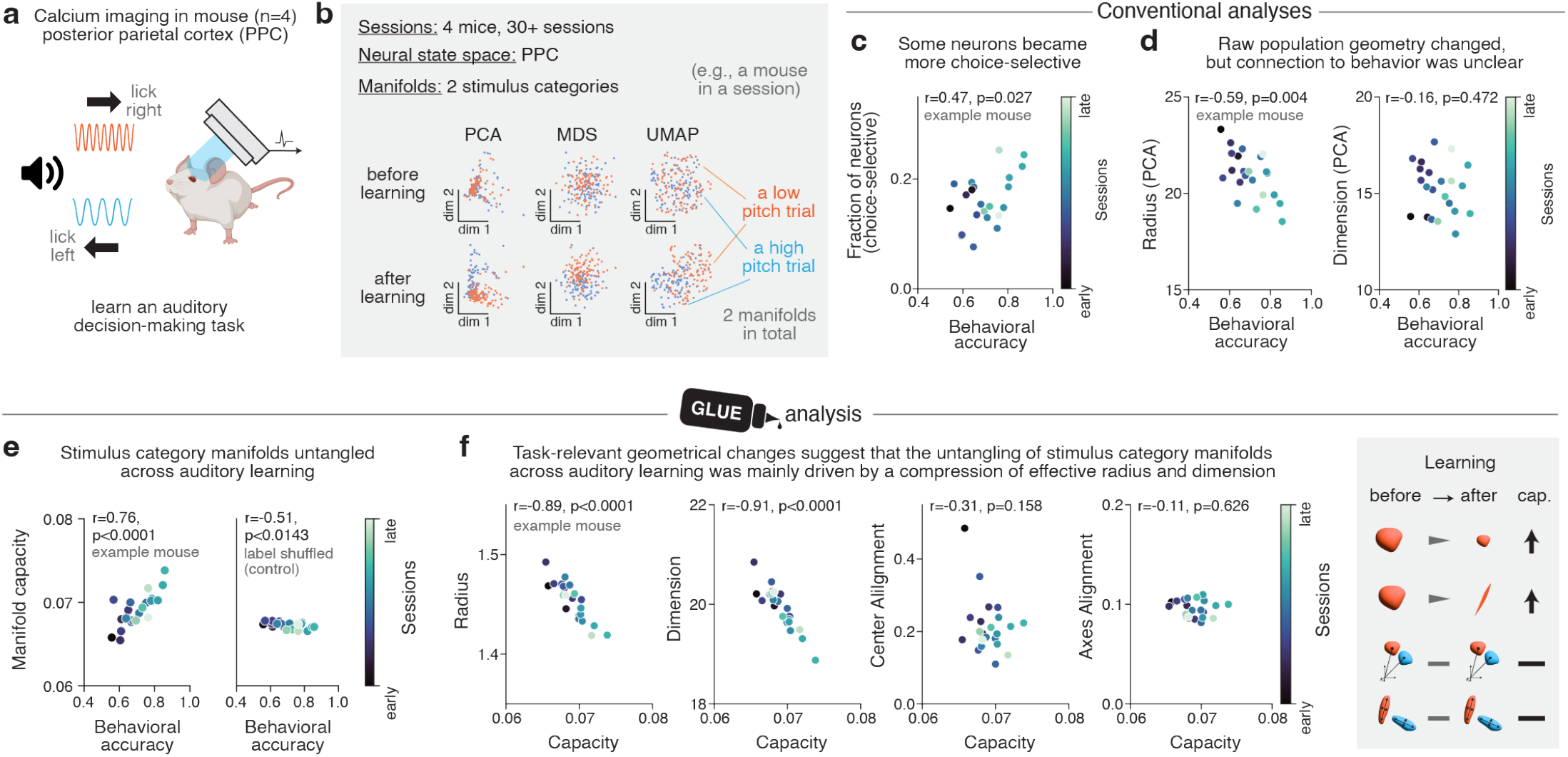
GLUE quantifies the task-relevant geometric transformations of stimulus category manifolds across auditory learning. **a**. A dataset from [38] consisting of calcium imaging recordings from posterior parietal cortex in mice (n=4). The mice were trained to report whether the rate of an auditory cue was high or low compared to a learned category boundary (16 Hz) by licking either a left or right waterspout. **b**. For a session, we defined 2 category manifolds. A category manifold contains the (inferred and preprocessed) neural responses to stimuli from the same cue category class (at least 50 trials per category). Visualization (using PCA, MDS, and UMAP) of the category manifolds. Top: an example session before learning. Bottom: an example session after learning. **c**. Single-neuron choice selective analysis. Fraction of neurons that are significantly choice selective were estimated for each session with a receiver operating characteristic (ROC) analysis (example mouse, 22 sessions, Pearson *r* = 0.47, *p* = 0.0270). **d**. Raw PCA-based population geometry analysis, where the radius and dimension are lower-order statistics of the eigenvalue spectrum of the manifold-wise covariance matrix. **e**. Manifold capacity analysis. Left: correlation between the capacity of category manifolds and behavioral accuracy (example mouse, 22 sessions, Pearson *r* = 0.76, *p <* 0.0001). Right: correlation between the capacity of label shuffled manifolds and behavioral accuracy (example mouse, 22 sessions, Pearson *r* = −0.51, *p* = 0.0143). **f**. Geometric analysis. (example mouse, 22 sessions; radius: Pearson *r* = −0.89, *p <* 0.0001; dimension: Pearson *r* = −0.91, *p <* 0.0001; center alignment: Pearson r=-0.31, p=0.158; axis alignment: Pearson *r* = −0.11, *p* = 0.626).

#### Auditory learning untangles and compresses stimulus category manifolds

We considered a dataset [38] involving mice (n=4) trained to report decisions about the repetition rate of multisensory event sequences (Figure 4a). The mice reported whether event rates were high or low compared to a learned class boundary (16 Hz) by licking either a left or right waterspout. After a few sessions, the mice learn to associate sensory stimuli with behavioral actions. During each session, neural activity in the posterior parietal cortex (PPC) was monitored through two-photon calcium imaging. Within the data from each session, we defined the high (or low) stimulus category manifold as the collection of population firing rates 0–97 ms before choice from the high (or low) rate trials (Figure 4b).

We started with conventional single-neuron analysis and analyzing the selectivity of individual neurons for decision outcome. We performed a receiver operating characteristic (ROC) analysis [43] on single-neuron responses (Method M2.2). A neuron was identified as choice selective when the area under the ROC curve (AUC) differed significantly (*p <* 0.05) from a shuffled distribution. We found that the fraction of choice-selective neurons was around 0.05 to 0.25 and this fraction was weakly correlated with the behavior accuracies (example mouse, 22 sessions, Pearson *r* = 0.47, *p* = 0.027; Figure 4d). This finding is consistent with the original analysis in [38] and indicates the existence of changes in PPC neurons’ tuning properties across learning. However, because only a small fraction of neurons (*<* 25%) are choice-selective, single-neuron analyses cannot explain the geometric transformations observed at the population level (see Figure 4b and Method M2.1). Moreover, while dimensionality reduction visualizations (Figure 4b) reveal changes in neural representations, they do not determine whether these changes are functionally beneficial. In particular, conventional geometric measures based on ellipsoidal approximations and PCA (Method M2.3)—lacking a direct connection to how the represented information are used—fail to extract and quantify the geometric transformations underlying representational changes across learning (example mouse, 22 sessions; radius: *r* = −0.59, *p* = 0.004; dimension: *r* = −0.16, *p* = 0.472; Figure 4d).

To overcome the limitations of conventional analyses, we applied GLUE to investigate the population-level mechanisms underlying mice learning an auditory decision-making task. We find that the capacity of stimulus category manifolds is highly correlated with behavioral accuracy (example mouse, 22 sessions, Pearson *r* = 0.76, *p <* 0.0001; Figure 4e, left), and these correlations were category-specific, as label-shuffled capacity values showed no significant correlation with behavioral accuracy (example mouse, 22 sessions, Pearson *r* = 0.51, *p* = 0.0143; Figure 4e, right). These findings suggest that the untangling of task-relevant stimulus category manifolds may be a key mechanism enabling mice to accurately associate stimulus categories with their decisions.

Next, we used GLUE to investigate the changes in effective geometry measures to uncover the transformations driving stimulus category manifold untangling during learning. We find that reductions in manifold radius and compressions of manifold dimensionality are the primary factors of increased manifold capacity (example mouse, 22 sessions; radius: *r* = − 0.89, *p <* 0.0001; dimension: *r* = − 0.91, *p <* 0.0001; Figure 4f), while alignment measures did not change significantly (example mouse, 22 sessions; center alignment: *r* = − 0.31, *p* = 0.158; axis alignment: *r* = − 0.11, *p* = 0.626; Figure 4f). These findings, revealed by GLUE, directly link computational-level measures (capacity) with structural changes in neural representations (effective geometric measures), revealing insights that previous methods could not capture.

### Using GLUE to reveal the degree of untangling and the geometric transformations involved in computations occurring within a neural population over time

#### Challenges in analyzing neural population representations in neural dynamics

While the learning-related changes we observed unfold over days to weeks, the brain must also rapidly reorganize neural representations to support moment-by-moment behavior. An example of this is motor planning. Recent studies indicate that the temporal dynamics of neural population activity are crucial for encoding and executing computations that drive goal-directed behavior [45]. While many analytical approaches rely on linear methods to characterize category manifolds, these techniques may be insufficient for capturing the full complexity of neural trajectories, particularly in behaviors that engage high-dimensional, non-linear dynamics across a broader region of neural state space [46]. To address this, we applied GLUE analysis to track how category manifold geometries evolve during individual trials of sensorimotor decisions (reaching tasks). We consider the hypothesis that behavior-relevant manifolds (e.g., the collection of neural responses to a reaching target, Figure 5b) become untangled throughout the temporal dynamics of a trial, allowing downstream areas to distinguish between different task conditions (e.g., distinct reaching targets).

**Figure 5:**
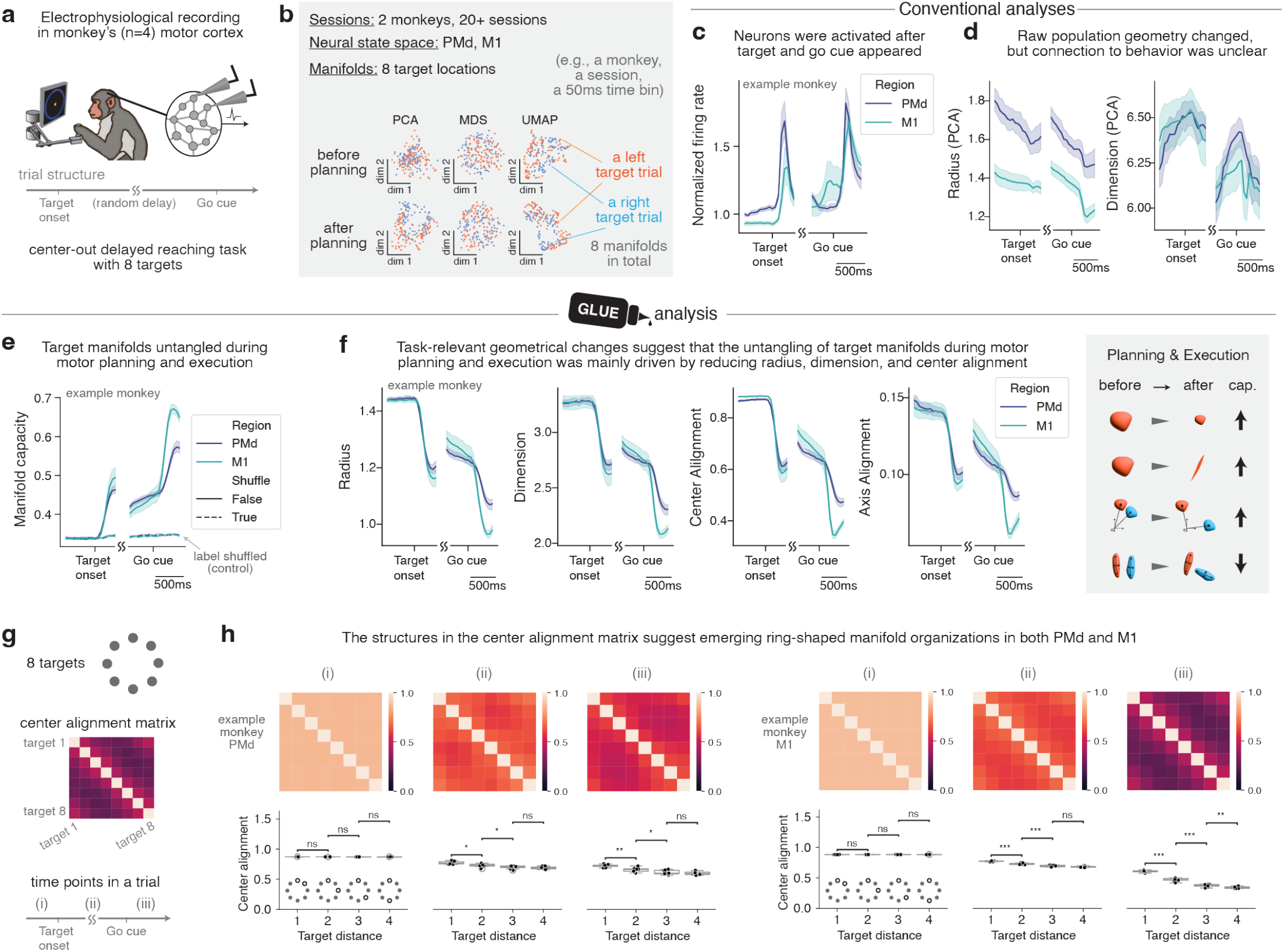
Motor planning and execution untangle target-specific manifolds. **a**. A dataset from [44] consisting of electrophysiological recordings from primary motor cortex (M1) and premotor cortex (PMd) of rhesus macaque monkeys (n=4). The monkeys were trained to perform a center-out reaching task with eight targets. **b**. For a trial, we time binned neural responses into 50ms frames and either center with respect to the target onset or from the go cue onset. For a session, we defined 8 target manifolds per time bin, where a target manifold contains the (averaged) neural responses to that target in the time bin (*>* 30 trials per target). Visualization (using PCA, MDS, and UMAP) of the stimulus category manifolds. Top: an example time bin before planning. Bottom: an example time bin after planning. **c**. Normalized firing rate analysis. A trial is divided into 50ms time bins and centered either with respect to target onset or go cue onset. The baseline firing rate was estimated for each neuron in a session as the average firing rate of the 300ms period before target onset. The normalized firing rate was the firing rate divided by the baseline firing rate. The error bands denote 95% confidence interval over the sessions (example monkey, 62 sessions; *>* 90 neurons for each session and brain area). **d**. Raw PCA-based population geometry analysis, where the radius and dimension are lower-order statistics of the eigenvalue spectrum of the manifold-wise covariance matrix. **e**. Manifold capacity analysis. The manifold capacity of target manifolds were measured per time bin and plotted. The error bands denote 95% confidence interval over the sessions (example monkey, 62 sessions; *>* 30 trials per manifold and session). **f**. Geometric analysis. (example monkey, 62 sessions; *>* 30 trials per manifold and session). **g**. Top: The targets lied evenly around a circle. Middle: We visualized the correlation between manifold centers by plotting an 8 by 8 matrix, with the *i, j* entry being the cosine between the center of the *i*-th and *j*-th manifold. Bottom: We sampled three time bins across different time point of a trial. **h**. The structures in the center alignment matrix suggest emerging ring-shaped manifold organization. Top: Visualization of the center alignment matrix at different time bins. Bottom: Averaged center alignment values of all pairs of manifolds with a certain distance (*k*) on the circle. After motor planning and movement, there were clear decreases of center alignment as *k* becomes larger, suggesting the centers of the manifolds form a ring-shaped organization.

To test this hypothesis, we analyzed two datasets: electrophysiology data [44] from monkeys completing a center-out reaching task (Figure 5) and electrophysiology data from monkeys completing a random-dot decision-making task set [47, 48] (Supplementary Figure S12). We observed an increase in the capacity of behavior-relevant manifolds during the within-trial temporal dynamics in both datasets. Moreover, for the first time, we were able to identify the key geometric transformations that drive the within-trial untangling process. These findings demonstrate that GLUE excels at capturing task-relevant representational changes in neural dynamics, and illustrate how adaptive behavior is supported by the geometrical reorganization of neural activity.

#### Motor planning and execution untangle target-specific manifolds

In the center-out reaching task, monkeys (n=4) trained on eight targets evenly distributed around a circle (Figure 5a). Neural activity was recorded from electrode arrays in the primary motor cortex (M1) and dorsal premotor cortex (PMd). We binned the neural spikes into 50ms intervals. For each time bin and target, we defined the target manifold as the collection of population firing rates on trials where the given target is cued (Figure 5b).

We first analyzed the temporal dynamics of average single-neuron firing rates. Neurons in both M1 and PMd fired significantly immediately after target onset and the “go” cue (Figure 5c). This finding is consistent with previous work [49–51] indicating abrupt changes in motor area activity during motor planning and execution, though its relation to task performance remains unclear. Next, we conducted population-level analyses by visualizing target manifolds using three dimensionality reduction methods (Figure 5b). While PCA, MDS, and UMAP hinted at underlying geometric transformations, definitive conclusions could not be drawn from visualization alone. Moreover, a straightforward quantitative analysis—estimating manifold radius and dimension via ellipsoidal approximation and PCA—did not reveal a clear trend along the trial dynamics (Figure 5d).

To overcome the limitations of the above conventional analyses, we used GLUE analysis to probe the task-relevant population-level changes in the temporal dynamics. We used manifold capacity to test the hypothesis that target manifolds become untangled during motor planning and execution. We find that the capacity of target manifolds increases sharply in both PMd and M1 immediately after the target was presented to the monkeys (Figure 5d). Similarly, target manifold capacity also increased immediately after the “go” cue onset. To make sure these representational changes were category-specific, we measured the capacity of label-shuffled manifolds and found no temporal changes (Figure 5d). These results indicate a two-stage target manifold untangling process occurring in a reaching task—one for planning and another for execution.

Next, we employed effective geometric measures from GLUE to investigate the structural factors that facilitate the untangling of target manifolds. We observed that manifold radius, dimension, center alignment, and axis alignment all sharply decreased following target onset and the go cue (Figure 5f). Furthermore, the correlation structure in the center alignment matrices suggests an emerging ring-shaped organization in both PMd and M1 (Figure 5g, Figure 5h), hinting at a correspondence between the circular task structure (with eight evenly distributed targets) and neural representations. These results not only exhibit much clearer trends than conventional geometric analyses (Figure 5d), but also closely linked with capacity changes through GLUE theory.

### Interpreting the computational advantages of manifold untangling within the biological context

While our analyses across different neural systems demonstrate the ubiquity of manifold untangling, a key question remains: What computational advantages does geometric reorganization provide? Our theoretical framework reveals three distinct but complementary interpretations of manifold capacity, each highlighting how untangling enhances neural computation. Higher capacity may indicate: (1) more efficient packing of categorical information, allowing more categories to be stored in the same neural state space (Figure 6a), (2) more efficient use of neuronal resources, enabling accurate decoding with fewer neurons (Figure 6b), or (3) more robust readout, reducing classification errors by downstream neurons (Figure 6c). The existence of three interpretations helps explain why manifold untangling is a common motif across diverse neural systems—it simultaneously optimizes multiple aspects of neural computation through coordinated changes in population geometry. In this case, the power of GLUE is that we can evaluate how well mathematical predictions based on a given computational principle match the neural data (Figure 6d-6f), thus revealing a possible biological interpretation.

**Figure 6:**
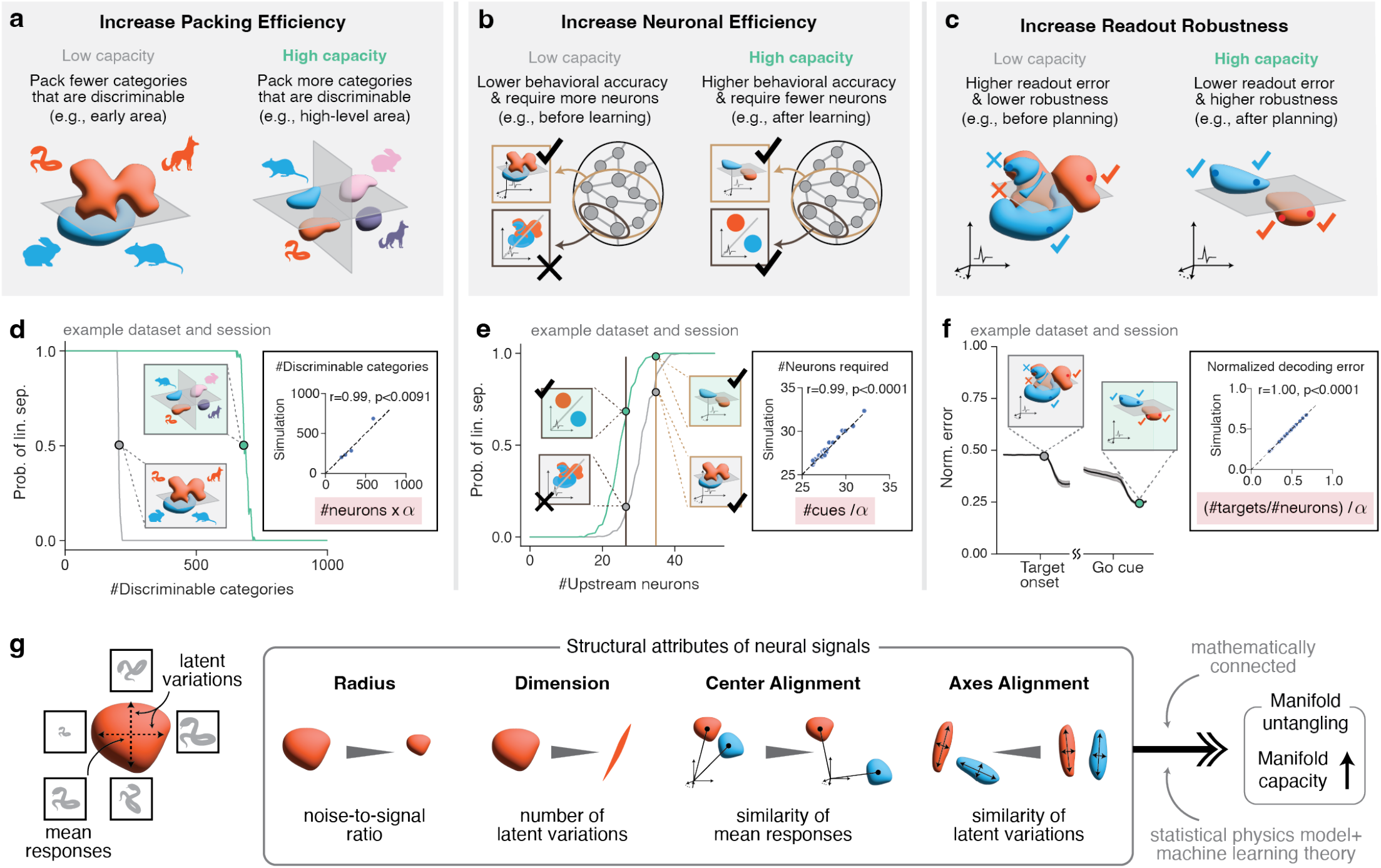
Biologically-relevant interpretations for manifold capacity and its effective geometric measures. **a**. Higher manifold capacity corresponds to higher packing efficiency, i.e., a neural activity space (e.g., a visual area) can accommodate more distinct categories while allowing downstream neurons to easily discriminate them. **b**. Higher manifold capacity corresponds to higher neuronal efficiency, i.e., a downstream neuron (modeled as a linear plus threshold readout) being able to discriminate between manifolds by reading out from fewer upstream neurons. **c**. Higher manifold capacity corresponds to higher readout robustness, i.e., the responses of a readout neuron (e.g., a motor neuron) to the same task condition (e.g., reaching toward a specific target direction) are more consistent across different trials. **d**. We quantify packing efficiency by the maximum number of linearly separable manifolds (Method M3.4). We considered deep neural networks (ResNet-50) pretrained on the Imagenet dataset. For each *P*, we randomly selected *P* categories and estimated the probability of linear separability of the corresponding stimulus category manifolds in that layer. We presented the results on an early layer (grey, conv1 layer) and a late layer (green, avgpool layer). Intercept: In our theory (Method M3.4 and Supplementary Information S1), the packing efficiency is connected to manifold capacity by the formula *P* ^*^ = *αN*, where *α* denotes manifold capacity. We numerically checked the theory in the intercept (Method M3.4). **e**. We quantify neuronal efficiency by the minimum number of upstream neurons required for a linear classifier to separate the manifolds (Method). We considered example sessions from ([38], see also Figure 4). For each *N*, we estimated the probability of the two stimulus manifolds being linearly separable after being projected to *N* random neurons. We presented the results on an early session (grey) and a late session (green). Intercept: In our theory (Method M3.4 and Supplementary Information S1), the neuronal efficiency is connected to manifold capacity by the formula *N* ^*^ = *P/α*. We numerically checked the theory in the intercept (Method M3.4). **f**. We quantify readout robustness by a normalized error (err^*^) that measures the fraction of responses that are not consistent with the majority response (Method M3.4). We presented the results of an example session from ([44], see also Figure 5) and estimated the temporal dynamics of the normalized error. Intercept: In our theory (Method M3.4 and Supplementary Information S1), the readout robustness is connected to manifold capacity by the formula err^*^ = (*P/N*)*/α*. We numerically checked the theory in the intercept (Method M3.4). **g**. Left: A neural manifold represents the collection of neural activity patterns associated with a task-relevant condition, such as an obkect category. The manifold center (mean activity responses) represents the prototypical condition, while the manifold axes correspond to latent variations. Right: The manifold radius reflects the noise-to-signal ratio, manifold dimension represents the number of latent variations, center alignment measures the similarity of mean responses, and axis alignment quantifies the similarity of latent variations across manifolds. These geometric measures are analytically connected to manifold capacity (Supplementary Information S1.4) via our theoretical framework using statistical physics model and machine learning theory.

#### Capacity quantifies the efficiency of packing categorical information

In the first interpretation, manifold capacity is linked to how many distinct categories a neural population can reliably encode (Figure 6a). Using the ribbon analogy, higher manifold capacity corresponds to being able to place more ribbons into the storage box while still ensuring easy retrieval. GLUE theory makes it possible to quantify this: providing a precise mathematical relationship between manifold capacity and the maximum number of separable manifolds that can be stored in a given neural state space while remaining discriminable by a linear classifier (Figure 6d). This relationship reveals that higher manifold capacity directly corresponds to greater storage efficiency—as manifolds become more untangled, more categorical information can be packed into the same neural space (Figure 6d, Method M3.4). This interpretation extends classical theories of neural perceptron capacity from point-like patterns to the more realistic case of continuous manifolds of activity.

#### Capacity predicts the minimal neuronal resources required for reliable decoding

Manifold capacity can also be used to determine how many neurons a down-stream decoder needs to reliably classify different categories (Figure 6b). Again using the ribbon analogy, higher manifold capacity corresponds to being able to store the same number of ribbons in a smaller storage box. Our theoretical framework establishes a quantitative relationship between manifold capacity and the minimum number of upstream neurons required for accurate classification (Figure 6e, Method M3.4). We validated this relationship in the mouse auditory learning dataset, where we found a striking correlation (*r* = 0.99, *p <* 0.0001) between theoretically-predicted and empirically-measured decoding efficiency (Figure 6e). This finding allowed us to interpret the increase of higher manifold capacity in PPC during learning as enabling downstream areas to correctly classify stimulus categories using progressively fewer neurons, demonstrating how geometric reorganization can enhance computational efficiency.

#### Capacity determines the robustness of down-stream readout

The third interpretation connects manifold capacity to the reliability, or robustness, of information transmission to downstream neurons (Figure 6c). In the ribbon analogy, higher manifold capacity corresponds to being able to easily grab the ribbon of interest. GLUE theory shows that higher capacity reduces the normalized decoding error—a measure of how consistently a downstream neuron responds to inputs from the same category (Figure 6f, Method M3.4). We validated this relationship in the monkey reaching task, finding that normalized decoding error in both PMd and M1 decreased sharply following target presentation and movement initiation, precisely when manifold capacity increased. The strong correlation (*r* = 0.98, *p <* 0.0001) between theoretical predictions and empirical measurements supports the interpretation that manifold untangling enhances the robustness of motor command transmission to downstream neurons (Figure 6f).

#### Interpretations for the effective geometric measures

Intuitively, the center of a manifold—defined by the mean activity responses (Figure 6g, left; see Methods)—can be viewed as the prototypical representation of that condition. Thus, internal coordinates within the manifold (i.e., manifold axes) correspond to possible latent variations of the condition, such as changes in the size of the input stimulus (Figure 6g, left). With this interpretation, the manifold radius can be understood as reflecting the noise-to-signal ratio, where the manifold center represents the signal, and the manifold axes capture the directions of noise (or nuisance variability). The manifold dimension corresponds to the number of directions spanned by latent variations. Lastly, center alignment quantifies the similarity of mean responses across manifolds, while axis alignment measures the similarity of latent variations (Figure 6g). Importantly, the effective manifold geometries are determined by the combination of each manifold’s underlying intrinsic geometry, its interaction with other manifolds, as well as the task structure, as opposed to a naive characterization of the geometry without taking the computation into account.

## Discussion

Our results demonstrate that analyzing neural manifold geometry provides fundamental insights into how the brain transforms complex sensory inputs into categorical representations that guide behavior. Through systematic application of GLUE analysis across diverse neural systems, we revealed both universal principles of neural coding and specific computational strategies adapted to different tasks. In the visual system, we found that object manifolds become progressively untangled along the ventral stream through systematic changes in their geometric properties—specifically through reductions in manifold radius and dimensionality. This untangling was particularly pronounced in category-selective regions like FFA and EBA for their preferred stimulus categories. In learning contexts, we showed that manifold capacity increases track behavioral improvement, revealing how neural circuits restructure their representational geometry to support newly acquired categorical associations. Finally, our analysis of motor planning and execution demonstrated rapid, dynamic reorganization of manifold geometry aligned with distinct behavioral phases, illustrating how neural populations can flexibly reconfigure their coding properties on millisecond timescales.

### The geometry of task-efficient coding

These findings significantly advance our understanding of neural computation in several ways. First, they provide a rigorous framework for quantifying task-relevant information processing that extends beyond classical efficient coding principles [52]. While efficient coding theory successfully explained how early sensory circuits compress information while preserving relevant statistics, it provided limited insight into how neural populations in downstream cortical areas selectively maintain task-relevant features while discarding irrelevant variations [53, 54]. GLUE bridges this gap by demonstrating how the geometric properties of neural representations directly influence a downstream decoder’s capability and efficiency in performing invariant tasks. This allows us to identify specific mechanisms—like reduced manifold radius or increased axis alignment—that enable more efficient encoding of categorical structure.

### Emergent structure in mixed selectivity neurons

Our results also help reconcile observations of complex mixed selectivity at the single-neuron level with orderly computation at the population level. Previous work has shown that individual neurons, particularly in higher brain areas, often respond to multiple task variables in apparently complex ways [3, 4]. GLUE reveals how these seemingly *messy* response properties can give rise to highly organized population geometries optimized for specific computations. The framework provides a principled way to analyze such distributed representations by quantifying how their geometrical structure supports task performance.

### Connection to other geometrical frameworks in neuroscience

GLUE connects to and extends existing geometrical frameworks in neuroscience in two important ways. First, it provides a robust alternative to visualization-based approaches for analyzing neural population structure. While dimensionality reduction techniques like t-SNE and UMAP have provided valuable insights [6, 1], recent work has shown that they can suggest categorical structure that reflects parameter choices rather than genuine organization in the data [7]. GLUE addresses this limitation by measuring geometric properties in the full high-dimensional neural space, without relying on potentially distorted low-dimensional projections. This becomes particularly crucial as neuroscience moves toward studying increasingly naturalistic behaviors, where maintaining rigorous links between neural activity and computation is essential.

Second, GLUE provides a new perspective on the ongoing debate about dimensionality in neural representations [55]. A fundamental puzzle in systems neuroscience is the apparent mismatch between the high dimensionality of neural populations (millions of neurons) and the lower dimensionality of behavioral variables they control (e.g., a few muscles or decision variables) [56, 57]. Our framework helps bridge this gap by explicitly connecting neural population size, manifold dimensionality, and computational capacity. Importantly, we show that dimensionality alone is insufficient to characterize neural representations—other geometric properties like manifold radius and organization play crucial roles in determining how effectively neural populations encode task-relevant information. This comprehensive geometric view helps reconcile seemingly contradictory observations about neural coding: high-dimensional mixed selectivity can support flexible computation, while low-dimensional structure can reflect key task variables.

### Population geometry of complex shapes beyond low-order statistics

The theory also provides new insights into information-limiting correlations in neural populations [12, 11]. While traditional approaches focus on pairwise statistics, GLUE can capture how complex patterns of stimulus-driven variability affect coding capacity. This is particularly important for understanding how neural circuits handle stimuli-driven variations in inputs [2], like changes in object pose or motion trajectories, which create structured correlations that cannot be characterized by simple statistical measures.

### Future directions

Several important questions remain for future investigation. First, while we focused on classification tasks, the framework could be extended to continuous-valued outputs or multiple simultaneous tasks [1]. This could help illuminate how neural circuits multiplex different computations or generate complex motor patterns. Second, our current analysis treats each time point independently—developing extensions for analyzing manifold dynamics could reveal additional principles of neural computation. Finally, establishing causal links between manifold geometry and behavior will require perturbation experiments guided by GLUE predictions.

The geometric principles revealed by GLUE may have broader implications beyond neuroscience. Similar challenges in transforming distributed representations into categorical patterns arise in fields ranging from artificial intelligence to structural biology. For instance, GLUE could provide insights into how deep neural networks organize information across layers [58, 59] or how cell-type-specific gene expression patterns emerge from complex molecular measurements [60]. More broadly, we propose that geometric measures like those developed here can serve as computational order parameters—quantities that capture how well a representation is organized to support specific computations. Just as order parameters in physics reveal universal principles governing phase transitions [61, 62], geometric measures may reveal general principles for how both biological and artificial systems optimize their representations.

In conclusion, GLUE provides a powerful new framework for understanding how neural circuits transform complex patterns of activity into functionally relevant representations. By linking geometry to computation, it reveals both universal principles and area-specific specialization in neural coding. The theoretical foundations and analytical tools developed here open up exciting directions for future work in neuroscience and beyond.

## Acknowledgements

We thank Billy Broderick, Jenelle Feather, Bo Hu, Jose Hurtado, Zeyu Jing, Hang Le, Artem Kirsanov, Sebastian Lee, Isabella Rischall, Sonica Saraf, Will Slatton, Brabeeba Mien Wang, and the members of Chung lab, for the discussion regarding many preliminary results and early versions of the manuscript. We thank Gabrielle Edgerton and Lucy Reading-Ikkanda for editorial support. We thank Corey Ziemba, and Tony Movshon for sharing their datasets. This work was supported by the Center for Computational Neuroscience at the Flatiron Institute, Simons Foundation, and by NIH grant R01DA059220. S.C. was partially supported by a Sloan Research Fellowship, a Klingenstein-Simons Award, and the Samsung Advanced Institute of Technology project, “Next Generation Deep Learning: From Pattern Recognition to AI.” All experiments were performed using the Flatiron Institute’s high-performance computing cluster.

## Data Availability

Monkey early vision dataset: Freeman et al. [35]. Monkey late vision dataset: Majaj et al. [33]. Monkey motor dataset: Perich et al. [44]. Monkey perceptual decision dataset: Kiani et al. [47, 48]. Human fMRI dataset: Hebart et al. [34]. Mouse spatial memory dataset: Plitt et al. [42]. Mouse decision-making dataset: Najafi et al. [38].

## Code Availability

Codes will be available for public usage in the final version. Requests on accessing to the current version of the code should be made to the first and corresponding author.

## Methods

### M1 Datasets

In general an analyst needs to choose *N* (number of units subsampled from those available) and *M* (number of examples/points subsampled from those available). For example, when quantifying manifolds in two recordings with 100 and 200 cells, one might subsample both to *N* = 50 to make a fair comparison. *N* and *M* should be chosen to lie within a range such that the resulting measurements are qualitatively insensitive to change of parameters.

#### M1.1 Monkey early vision dataset

We use the dataset from [35]. 13 anesthetized macaques were presented with a set of 450 images of 15 naturalistic textures while neural responses were recorded from visual cortex. Each stimulus is an intact or scrambled version of one of 225 grayscale images. Each intact image is one of 15 randomly synthesized examples of a texture from one of 15 texture families. The images were scrambled by randomizing phase in the 2D Fourier domain, preserving a naturalistic power spectrum while destroying the high-order statistics distinguishing texture. The stimuli were presented multiple times in pseudorandom order. Presentation lasted 100 ms with a 100 ms gap between presentations. Responses were recorded from several neurons at a time using quartz-platinum tungsten microelectrodes (Thomas Recording) in cortical areas V1 and V2. Responses to 20 repeats of all stimuli were collected for 102 neurons in V1 and 103 neurons in V2.

The neural responses are a collection of population vectors corresponding to stimulus presentations. A population vector contains spike count in the 100 ms following response onset for each recorded neuron. In the pixel data, a population vector contains the intensity of each pixel in one image in the stimulus set. We produce a set of manifolds for each population pixels, V1 and V2. For each population, we create 15 manifolds by grouping together neural responses to 15 different textures. Each manifold is a collection of presentation-averaged neural responses to a random subset of 10 images sampled from one texture. We randomly sample responses from a subset of 90 neurons in the ensemble. For pixel data, we use random projections to sample 90 dimensions from the 102400 pixel dimensions. We assemble separate manifold sets from responses to scrambled and intact images. (We also compare against a random baseline where population responses are shuffled with respect to the stimuli before grouping into manifolds.)

We use GLUE to compute manifold capacity, radius, dimension, axis alignment, center alignment and center-axis alignment for each neural population. We show error bars across 10 random runs, and use a Mann-Whitney test to determine whether geometric measures differ significantly between neural populations.

#### M1.2 Monkey late vision dataset

We use the dataset from [33]. Two macaques were presented with images of objects rendered against naturalistic backgrounds while neural responses were recorded from visual cortex. Each stimulus is one of 5760 grayscale images of a 3D object rendered against a randomly selected naturalistic background. The object is one of 64 3D models, where 8 example models belong to each of 8 object categories (animals, boats, cars, chairs, faces, fruits, planes, and tables). 90 stimuli are rendered from each 3D model by varying object pose, scale, location. Renderings are made at three levels of variation: none (10 images), medium (40 images), and high (40 images). The stimuli were presented in series during passive fixation. Presentation lasted 100 ms with a 100 ms gap between presentations. Population responses were recorded using three 96-channel multi-electrode arrays in cortical areas V4 and IT. Responses to all stimuli were isolated for 189 sites in IT and 126 sites in V4 (a typical site captured spiking activity from 1-4 neurons).

The neural responses are population vectors corresponding to stimulus presentations. A population vector contains the z-scored number of spikes from 50 to 150 ms after stimulus onset at one electrode site. We produce a manifold set for each population pixels, V4 and IT by grouping neural responses to images of the same 3D object. Each manifold is composed of responses from a random subset of 100 units to 30 randomly selected images of the object. For pixel data, we use random projections to sample 100 dimensions from the 196608 pixel dimensions. We assemble separate manifold sets from images at all 3 levels of variation. For a random baseline, we shuffle neural responses with respect to the stimuli before grouping into manifolds.

We use GLUE to compute manifold capacity, radius, dimension, axis alignment, center alignment and center-axis alignment for each neural population. We aggregate results across 10 random runs and apply a Mann-Whitney test for statistically significant differences between populations.

#### M1.3 Monkey reaching dataset

We use the dataset from [44]. Two macaques performed an eight-target center-out reaching task while neural responses were recorded from motor cortex.

A recording session consisted of about a hundred trials across three consecutive epochs: baseline, adaptation and washout. During baseline trials, the reaching task was not perturbed in any way. During adaptation trials, a motor perturbation was applied where either external forces were applied to the hand (“curl field”) or a fixed rotation was applied to the visual feedback (“visual rotation”). Population responses were recorded using 96-channel multi-electrode arrays in cortical areas PMd and M1. Spike sorting was applied at the session level (9 curl-field and 7 visual-rotation sessions). Across all sessions, 137–256 PMd and 55–93 M1 neurons were isolated for Monkey C, and 66–121 PMd and 26–51 M1 neurons were isolated for Monkey M. The neural responses are spike rates from each neuron at varying time throughout a reaching task. Spike rates are derived by binning spike trains in 0.1 s intervals and convolving with a Gaussian kernel of bandwidth 0.2 s.

We produce manifold sets for each motor cortex population by grouping together neural responses which occur during the same reach direction, under the same trial block, and at the same time with respect to the planning period of the trial.

We run two analyses on the manifold geometry, one for studying the time course of neural responses across a trial and one for studying the effects of adaptation to the motor perturbation throughout a session. In the first analysis, we produce manifolds by grouping together neural responses which occur during the same reach direction and at the same time bin throughout the course of a trial. We use a time bin of width 0.2 s with step size 0.1 s to sample manifold sets at varying times throughout the trial. At each time step, we assemble direction manifolds by sampling 10 neural responses within the time bin for each of the 8 reach directions. We use responses from a random subsample of 50 units for each manifold set. In the first analysis, we compute the six manifold capacity metrics for PMd and M1 separately, and show each measured value as a function of time averaged across 5 random runs.

In the second analysis, we create manifolds at the session level by sampling from neural responses occurring under the same reach direction and adaptation condition with respect to the motor perturbation. Correlations among these manifolds encode information about how the population representation changes structurally to adapt to perturbation of the task.

We delineate adaptation conditions by establishing “early” and “late” trial blocks within the “adaptation” and “washout” epoch, as well as a “baseline” block of trials before the perturbation is induced. We define early and late blocks respectively as trials before and after the median trial index throughout the epoch, restricting analysis to those trials ending in reward. We then compute GLUE effective alignment matrices among the 40 reach direction-trial block manifolds. We run the analysis separately on manifolds sampled from before and after the onset of the reaching motion. We plot alignment matrices two ways by different ordering of the columns: first by assigning to adjacent columns manifolds from the same adaptation condition, and then by assigning to adjacent columns manifolds associated with the same reaching direction.

The main text shows analysis of the curl-field perturbation session on date 2016-09-15 (which is the first curl-field session).

#### M1.4 Monkey perceptual decision dataset

We use the dataset from [47, 48]. Two macaques performed a random-dot direction discrimination task while neural responses were recorded from prearcuate cortex.

In each task trial, two targets appeared on the screen after the monkey directed its gaze towards a central point. After a 250 ms delay, a random dot motion stimulus was presented for 800 ms, followed by a variable-duration delay period. Upon presentation of a Go cue, the monkeys were instructed to indicate their perceived motion direction by executing a saccade toward the corresponding target. Motion strengths were tuned for each monkey to provide a broad range of behavioral accuracies, ranging from chance to flawless performance.

Population responses were recorded using 96-channel multi-electrode arrays in prearcuate gyrus. Spike sorting identified 100-250 units in each recording session (a unit could be an isolated single neuron or a multi-unit), for 15 sessions in total. The neural responses are spike rates from neural units at varying time throughout a reaching task. Spike rates are derived by binning spike trains in 0.1 s intervals and convolving with a Gaussian kernel of bandwidth 0.2 s.

We produce manifold sets by grouping together neural responses which occur under the same dot-motion conditions and at the same time bin throughout the course of a trial. We use a time bin of width 0.2 s with step size 0.2 s to sample manifold sets at varying times throughout the trial. At each time step, we assemble dot-motion manifolds by sampling neural responses within the time bin from trials with the same motion coherence and direction. Each of the 12 dot-motion manifolds (6 for coherence levels and two opposing directions) consists of 20 randomly sampled population vectors from the same subset of 50 neural units. We restrict analysis to trials where the perceptual decision was correct.

We compute manifold capacity metrics for each time bin separately, and show measured values as a function of time averaged across 10 random runs. Time ranges from -1.0 s to 3.0 s where 0 is dot stimulus onset. We also show the correlation matrices for manifolds from four select time bins throughout the evidence integration period of the trial (0.0, 1.0, 2.0 and 2.2 s).

#### M1.5 Mouse auditory decision-making dataset

We use the dataset from [38]. Four mice performed an auditory decision task while neural responses were imaged in posterior parietal cortex. Trials consisted of simultaneous clicks and flashes, generated randomly (via a Poisson process) at rates of 5–27 Hz over 1,000 ms. Mice reported by licking a left or right waterspout whether event rates were high or low compared with a category boundary (16 Hz) learned from experience. A two-channel, two-photon microscope was used to record the activity of neurons in layer 2/3 of posterior parietal cortex in transgenic mice. The images were processed by an algorithm that simultaneously identified neurons and estimated *inferred spiking activity*, a number related to a neuron’s spiking activity during one frame. Recordings were made for 135 sessions in total, with a typical session isolating activity from 400-600 cells.

The neural responses are the inferred spiking activities during each frame of ROIs extracted from calcium images. Each ROI correspond to individual neurons identified during post-processing. The population responses are sampled at 30.9 Hz. We produce manifold sets by grouping together neural responses during trials with the same stimulus condition: high- or low-frequency multisensory stimulation. Each manifold is 50 randomly sampled population vectors from all recorded neurons. Manifolds are constructed from responses in the 100 ms following the go tone of each trial.

Recordings span multiple days of each individual learning the task. We run GLUE analysis on manifolds for each recording session and compare across days. For each session, we aggregate manifold metrics across 10 random runs. Then, for one individual, we plot each measured manifold metric as a function of task performance, where the points are colored by the day on which that measurement was taken throughout the course of learning.

#### M1.6 Mouse spatial memory dataset

We use the dataset presented in [42]. Ten mice performed spatial navigation along a linear track in a virtual-reality environment which took on 5 different contexts along a sensory continuum. The mice were separated into two cohorts: 4 “frequent”, and 6 “rare” animals. The ‘frequent’ cohort was habituated to all 5 contexts during training preceding experimental recording sessions. The ‘rare’ cohort individuals were only exposed to 2 of the 5 environments regularly during training (the other 3 were presented only very occasionally). Calcium images were captured at 15.46 Hz from a 1 mm^2^ field of view in hippocampal area CA1 during recording sessions lasted 60-120 trials, or approximately 40 minutes. Images were processed by a segmentation algorithm, identifying between 161 and 3657 ROIs per session as pyramidal neurons. Instantaneous activity (not interpreted as a spike rate) was computed for each ROI by deconvolving the indicator signal with a canonical calcium kernel.

The neural responses are activity rates estimated framewise for each ROI from calcium images. Each ROI corresponds to a pyramidal neuron identified during post-processing. For each session, we identify place cells with respect to each context using the spatial information criterion described in [42] We restrict analysis to the union of cells identified as place cells across all of the 5 contexts presented in the session. This gave a median of 986 (minimum 56, maximum 2015) cells per session. We compute capacity without subsampling from this population. For each session, we create a set of 5 manifolds by grouping together hippocampal responses from each of the 5 contexts. Each manifold is constructed from 10 population vectors sampled randomly from among all responses during time points following trial reward. We assemble a manifold set for each session recording from each individual. We use GLUE to compute manifold capacity, radius, dimension, axis alignment, center alignment and center-axis aligment of the recorded neural population. We show error bars across 50 random runs and use a Mann-Whitney test to determine whether geometric measures differ significantly between neural populations from each cohort.

#### M1.7 Human fMRI dataset

We use the dataset presented in [34]. Three human participants were presented with images of objects while fMRI data were collected. The images were taken from the THINGS object concept and image database [63]. Each stimulus is one of 8640 colored images of an object embedded in a natural background. The object belongs to one of 720 distinct concepts, with each concept consisting of 12 exemplars. The stimuli were presented in series while maintaining central fixation. The presentation lasted 500 ms with a 4000 ms gap between presentations. To maintain engagement, participants were instructed to perform an oddball detection task, detecting occasional artificially generated images amidst the stimulus presentation series.

Whole-brain fMRI data were collected using a 3 Tesla Siemens Magnetom Prisma scanner and a 32-channel head coil with 2mm isotropic resolution. Functional data were acquired for the main experiment for 12 scanning sessions, and functional localizers were acquired for additional 1-2 scanning sessions. The neural responses are single-trial response estimates of the BOLD response amplitude to each object image from each voxel. Single-trial estimates are derived by fitting a single-trial general linear model on the fMRI time series. Responses from 19 visual ROIs are used. (V1, V2, V3, hV4, VO1, VO2, TO1, TO2, V3b, V3a, EBA, FFA, OFA, STS, PPA, RSC, TOS, LOC, and IT). We select three higher-level categories (face, body part, and electronic device) and produce a set of manifolds for each higher-level category and for each ROI. For each ROI, we create 4-5 manifolds by grouping together neural responses to 4-5 different concepts belonging to the same higher-level category. Each manifold is a collection of neural responses to a random subset of 10 images sampled from one concept. We randomly sample responses from a subset of 400 voxels. We assemble subordinate manifolds from three higher-level categories—face, body part, and electronic device (as a control).

We use GLUE to compute manifold capacity, radius, dimension, axis alignment, center alignment, and center-axis alignment for each ROI and each higher-level category. We show error bars across 50 replicates and do a two-sample t-test for statistically significant differences between higher-level categories within each ROI.

#### M1.8 Artificial neural networks

We use the standard pretrained ResNet-50 [64] and Vision Transformer (ViT) [65] architecture trained on ImageNet from PyTorch. The neural responses are extracted from a pretrained ResNet-50 architecture trained on ImageNet. We focus on 19 layers in ResNet-50: conv1, relu, layer1.0.relu, layer1.1.relu, layer1.2.relu, layer2.0.relu, layer2.1.relu, layer2.2.relu, layer2.3.relu, layer3.0.relu, layer3.1.relu, layer3.2.relu, layer3.3.relu, layer3.4.relu, layer3.5.relu, layer4.0.relu, layer4.1.relu, layer4.2.relu, and avgpool; and focus on 14 layers in ViT: conv_proj, encoder.layers.encoder_layer_*i* for *i* = 0, 1, …, 11, and encoder.ln. For each random repetition, we fix a random projection matrix for each layer and project the neural activations to a 2000 dimensional subspace. In each repetition, we randomly select 50 categories from the 1000 ImageNet categories and randomly select 30 images from the top 10% accurate images of each category to for the manifolds (we follow the same protocol as in [28]). We run GLUE analysis (as well as previous analysis methods for a comparison) on the manifolds of each layer and in total we conduct 5 random repetitions.

### M2 Conventional analyses

#### M2.1 Visualization

For data visualization in Figure 3b, Figure 4b, Figure 5b, we used Python package sklearn to conduct PCA and MDS analysis, and used Python package umap to conduct UMAP analysis. In all plots, we used the default setting and plotted the top 2 components.

#### M2.2 ROC analysis

In the ROC analysis presented in Figure 4c, we measured the choice preference of individual neurons using the area under the ROC curve (AUC). Choice selectivity was quantified as twice the absolute difference between the AUC and the chance level (0.5), defined by the formula: choice selectivity = 2|AUC−0.5|. To identify neurons with significant choice selectivity, we performed ROC analyses on each neuron using shuffled trial labels, where left and right choices were randomly reassigned to trials. We repeated this shuffling process 50 times to estimate a distribution of AUC values under the null hypothesis for each neuron. A neuron was considered choice-selective if the probability of its actual AUC occurring within the shuffled AUC distribution was less than 0.05. See also Supplementary Figure S5 for an illustration of the (intuitive) connection between ROC analysis and manifold capacity.

#### M2.3 Manifold geometry analysis based on ellipsoidal approximation

In the raw population geometry analysis presented in Figure 4d and Figure 5d, we first measured the manifold-wise centered covariance matrix. For each manifold, we then computed the eigenvalues of its covariance matrix. Let *λ*_1_, …, *λ*_*N*_ be the eigenvalues of a covariance matrix and let *ℓ* be the center norm of the manifold (i.e., the *ℓ*_2_ norm of the barycenter of the manifold), the PCA radius and PCA dimension are defined as 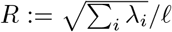 and 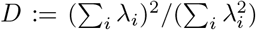 respectively. We then averaged the radius and dimension over all manifolds and plotted the results.

### M3 GLUE Framework and Theory

Here we define manifold capacity and effective geometric measures in a data-driven manner. For more details on the derivation, intuitions, and interpretations, please refer to the Supplementary Information S1.

#### M3.1 Notations

In the data-driven scenario, a manifold is modeled as a point cloud. For the simplicity of the presentation, here we assume every manifolds have the same number of points. Let *N* be the number of units (e.g., neurons, voxels), let *P* be the number of manifolds, and let *M* be the number of points per manifold. The *µ*-th manifold ℳ^*µ*^ consists of a collection of points 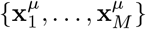 where 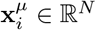 for each *i* = 1, …, *M*. In the analysis, **x** can be a population firing rate vector or a single-trial beta coefficient vector estimated from blood oxygenation level dependent (BOLD) responses. Without loss of generality, we assume points in the same manifolds are linearly independent (which can be done by adding tiny Gaussian noises if *N* ≥ *P* × *M*). In a data analysis, a manifold corresponds to a condition (e.g., the behavior outcome) and, its points are the (preprocessed) neural activity vectors from the trials in the condition.

#### M3.2 Manifold capacity

Given a collection of *P* data manifolds {ℳ^*µ*^} each containing *M* points in an *N*-dimensional ambient space. Let **y** ∈ {−1, 1} ^*P*^ be an indicator vector of a dichotomy. Namely, *y*^*µ*^ is the label associated to the *µ*-th manifold. The *simulated capacity* of these manifolds with respect to the label is defined as follows.

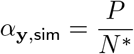

where *N* ^*^ is the smallest 1 ≤ *N* ^′^ ≤ *N* such that the probability of randomly projecting {ℳ^*µ*^} to an *N* ^′^-dimensional subspace while maintaining linear separability (w.r.t. **y**) is at least 0.5. While *α*_**y**,sim_ can be numerically estimated via binary search plus support vector machine, it is not analytical. In this work, we derive via an analytical formula *α*_**y**,glue_ that’s both provably and empirically well approximate *α*_**y**,sim_:

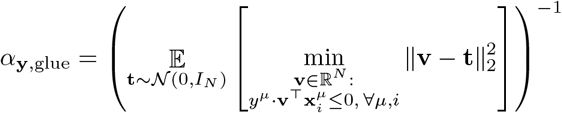

where 𝒩 (0, *I*_*N*_) stands for the multivariate isotropic Gaussian distribution in ℝ^*N*^. In many settings, we are interested in more than one dichotomy of the *P* manifolds. For a collection of dichotomies 𝒴 ⊆{−1, 1} ^*P*^, the corresponding manifold capacity is

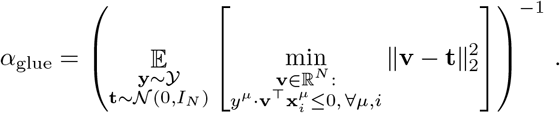

To estimate manifold capacity, we randomly sample *n*_*t*_ = 200 (**y, t**) pairs (corresponding to the 𝔼 in the above formula) and estimate the minimum value of 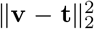 by solving a quadratic program. See Algorithm 1 for a pseudocode.

We consider the *one-versus-rest* dichotomy scheme in all the analyses in this paper. Namely, 𝒴= {(1, −1, −1, …, 1), (−1, 1, −1, …, −1), …, (− 1− 1, …, −1, 1)}. See Supplementary Information S1 for more details on other choices of dichotomies that could be useful in other data analysis.

#### M3.3 Effective geometric measures from manifold capacity formula

For each (**y, t**) pair, the optimization problem inside the expectation of the above capacity formula can be rewritten as follows, using the strong duality property.

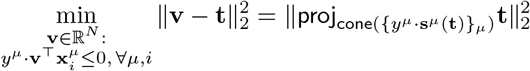

where **s**^*µ*^(**t**) = *y*^*µ*^ · **x**^*µ*^(**y, t**) and **x**^*µ*^(**y, t**) ∈ ℳ^*µ*^ is the support vector for the left-hand side of the above equation (see Supplementary Information for a derivation), cone denotes the linear cone span of a collection of vectors, and proj denotes the projection operator. In previous work [20], **s**^*µ*^ is also called *anchor point*. As (**y, t**) is randomly sampled, the induced anchor points form a distribution over the data manifolds. Importantly, this *anchor distribution* is analytically linked to the manifold capacity via the above equations, and we call the induced geometry as *anchor geometry*.

Finally, the effective geometric measures are defined on top of the anchor distribution. Following ideas from previous work [20], we decompose an anchor point **s**^*µ*^(**y, t**) into a center part 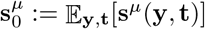,which is independent to **t**, and an axis part 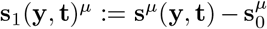.Let 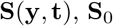,and **S**_1_(**y, t**) be the *P*× *N* matrices with their *µ*-th row being 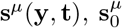,and 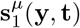 respectively. In the following, we directly define the effective geometric measures and leave the intuitions and discussions to Supplementary Information. Let

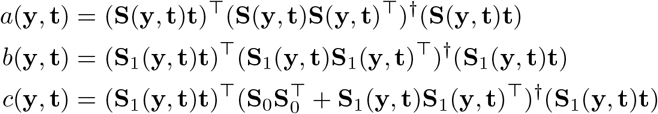

where † stands for pseudo-inverse. Note that 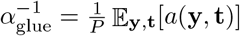.

- (Effective manifold dimension)

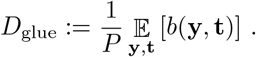
- (Effective manifold radius)

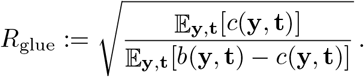
- (Effective utility rate)

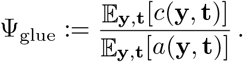

Note that the above three effective geometric measures give rise to the following *geometric formula* for manifold capacity.

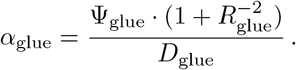

We provide derivations, intuitions, and examples of the above three measures in the Supplementary Information. Next, we also define the following effective manifold alignment measures as an average of some second-order statistics of anchor points.

- (Effective center alignment)

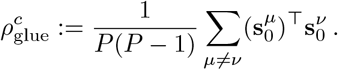
- (Effective axis alignment)

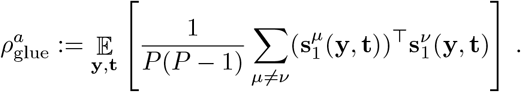
- (Effective center-axis alignment)

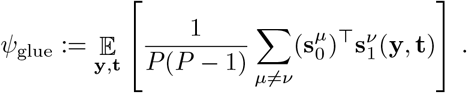

See Algorithm 1 for a pseudocode to estimate the above geometric measures.

#### M3.4 Equivalence of capacity to three normative objectives

Manifold capacity *α* is analytically linked to three distinct normative principles. This allows for broader interpretations for manifold untangling in different tasks and systems in neuroscience. Recall that *P* is the number of manifolds and *N* is the number of neurons in the data.

##### Packing efficiency

Higher the manifold capacity corresponds to the neural state space being able to store more distinct task conditions. We define the largest number of packable manifolds in the state space as *P* ^*^. Numerically, we estimate *P* ^*^ by first fixing the number of neurons to be *N*. Then, for each *P*, we estimate the probability of linear separability of *P* random manifolds and a random dichotomy by subsampling and repeating for 50 times. Finally, we let *P* ^*^ be the largest *P* such that the (estimated) probability of linear separability is at least 50%. In Figure 6d, we picked *N* = 2048 and subsampled from 1000 Imagenet categories.

Mathematically, we establish the following connection between manifold capacity *α* and packing efficiency: *P* ^*^ ≈ *αN*.

##### Neuronal efficiency

Higher manifold capacity also corresponds to a downstream neuron being able to correctly classify different task conditions by reading out from less number of upstream neurons. We define the smallest number of upstream neurons required as *N* ^*^. Numerically, we estimate *N* ^*^ by first fixing the number of manifolds to be *P*. Then, for each *N*, we estimate the probability of linear separability of the random projection of these manifolds and a random dichotomy by projecting them to a random *N*-dimensional subspace and repeating for 50 times. Finally, we let *N* ^*^ to be the smallest *N* such that the (estimated) probability of linear separability is at least 50%. In Figure 6e, we picked *P* = 2 and subsampled from *N* = 400 neurons.

Mathematically, we establish the following connection between manifold capacity *α* and neuronal efficiency: 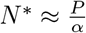.

##### Readout robustness

Higher manifold capacity also corresponds to less average error made by downstream neurons to discriminate different task conditions. We define a normalized error as

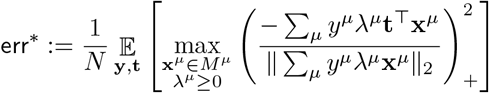

where **t** can be interpreted as a random linear classifier and ∑*λ*^*µ*^**x**^*µ*^ is a point from the convex hull of the data manifolds. Namely, the right-hand side of the above equation is analogous to an average case notion of the negative margin in support vector machine theory. Numerically, we estimate err^*^ by sampling 200 random **t** and solve the inner optimization problem by a quadratic programming solver.

Mathematically, we establish the following connection between manifold capacity *α* and decoding robustness: 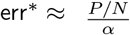.

#### M3.5 Algorithms for estimating capacity and geometric measures

The algorithm that computes capacity and geometric measures from data (e.g., {ℳ^*µ*^}) consists of two steps: (i) sample anchor points and (ii) calculate capacity and effective geometric measures from the statistics of these anchor points. The following are the pseudocodes for each step. In the following algorithms, **A** ← {**x**_*i*_} denotes putting the collection of vectors {**x**_*i*_} on the rows of **A**.

##### Algorithm 1 Pseudocode for estimating manifold capacity and geometric measures

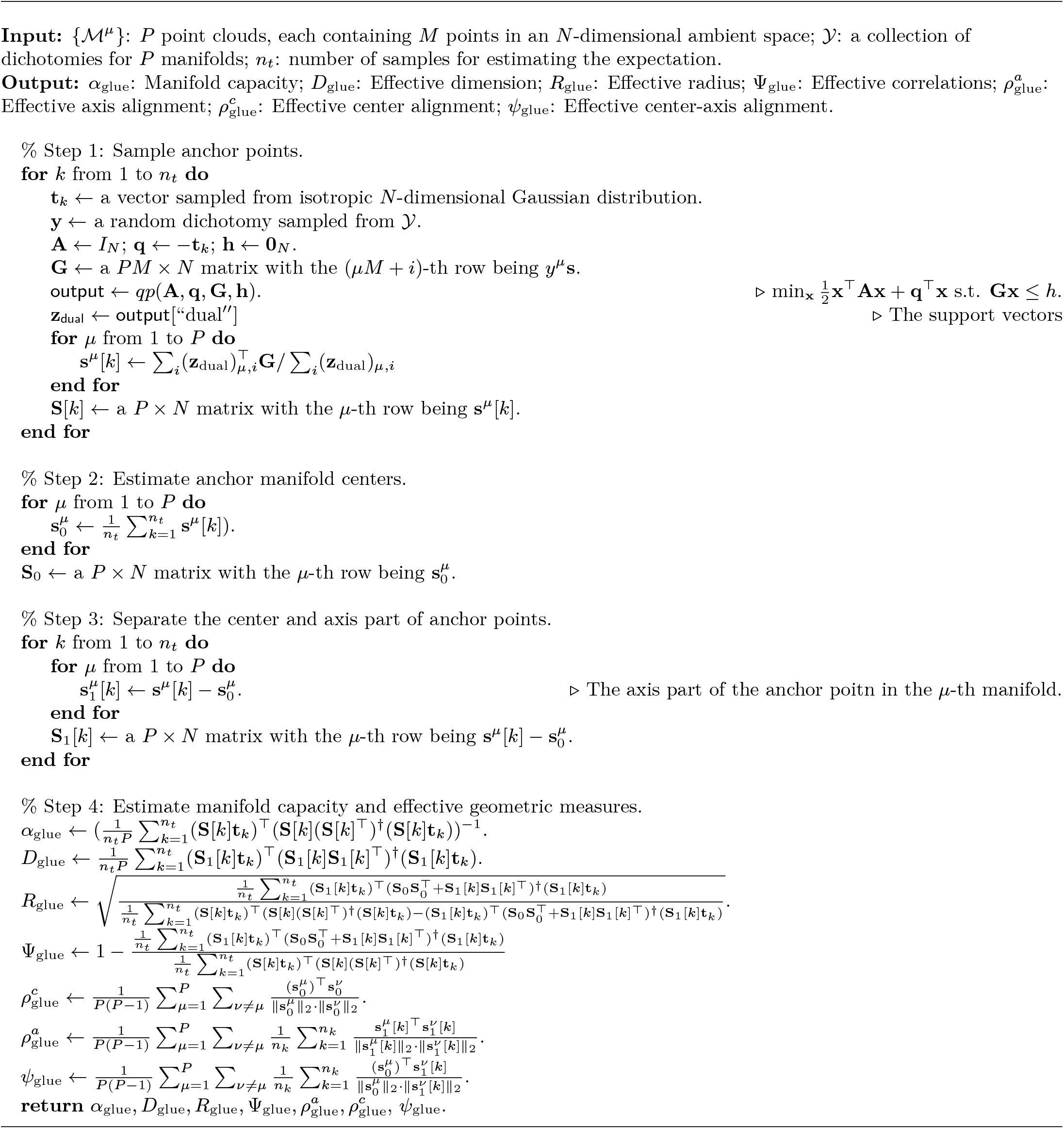

## Supplementary Information

This supplementary material contains four parts. In Supplementary Information S1, we provide a complete derivation of the Geometric measures from capacity of manifold categorization (GLUE) as well as relevant interpretations. In Supplementary Information S2, we use two toy models to illustrate the intuitions of GLUE. Finally, in Supplementary Information S3, we delineate the missing details and interpretations of the data analysis in the main text.

**Figure S1:**
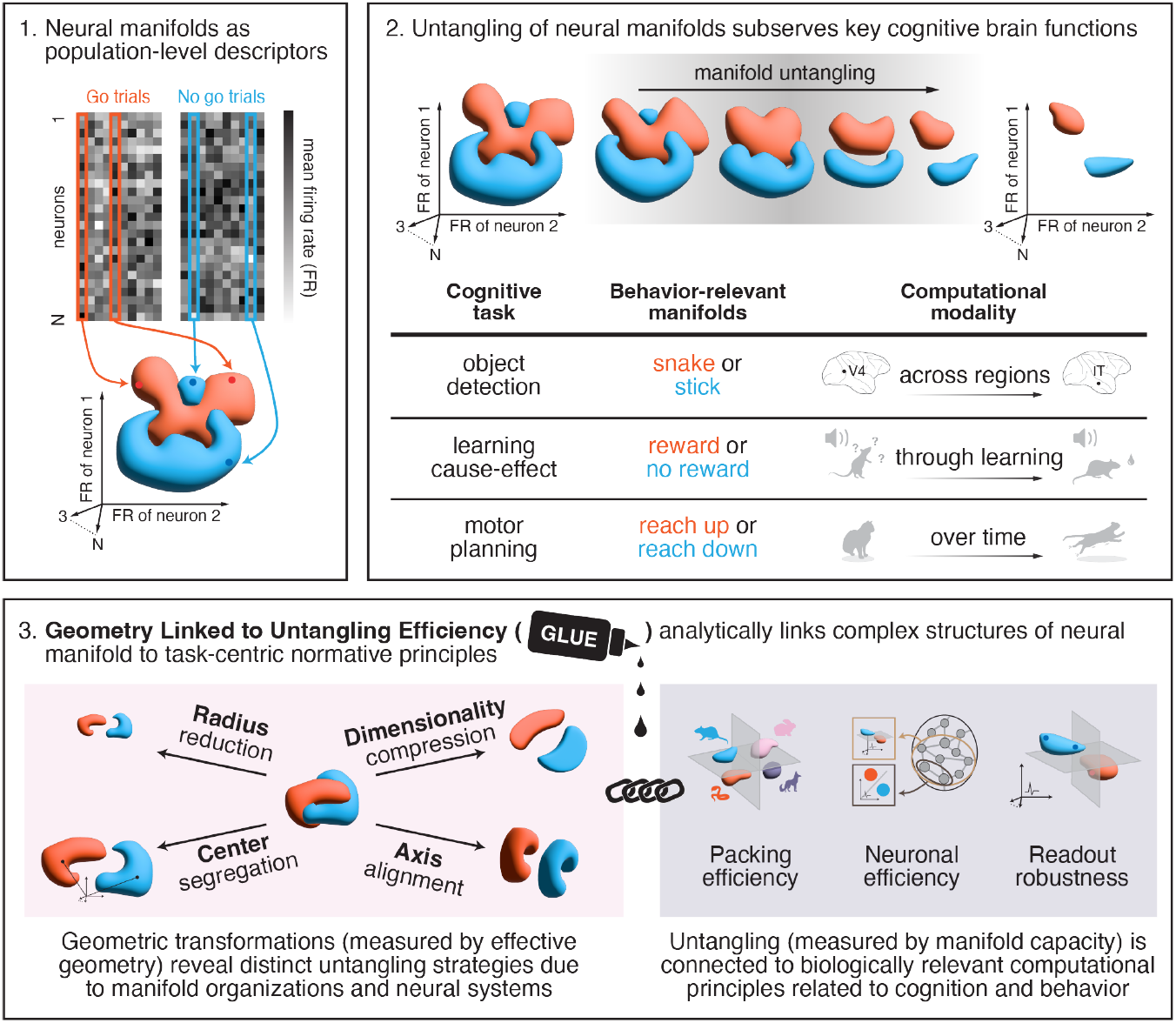
Graphical abstract.

**Figure S2:**
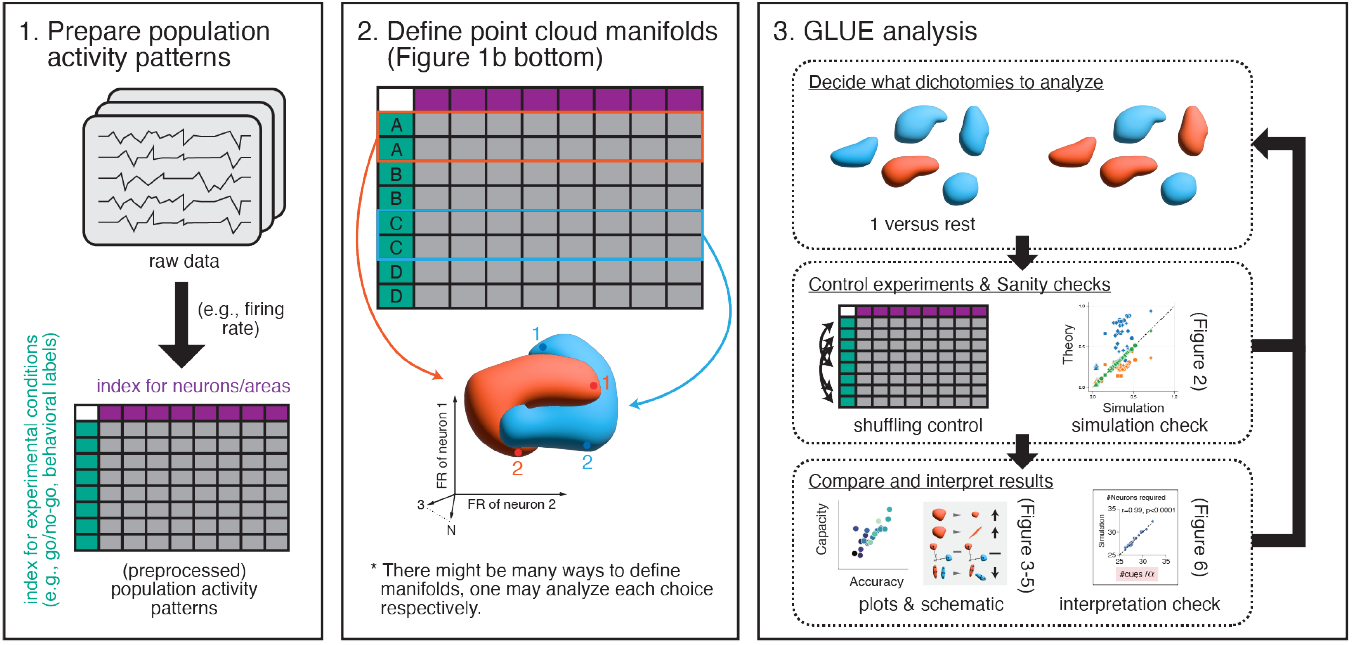
Analysis flow of using GLUE.

### S1 Manifold Capacity Theory in GLUE

Manifold capacity, *α*, aims to quantify the largest^3^ ratio *P/N* where *P* is the number of manifolds *P* that can be packed in an *N*-dimensional neural state space while being easily discriminable (e.g., by a linear classifier). In data analysis, we only have access to a finite number of manifold, *P*_data_, and a finite number of neurons, *N*_data_. To access the manifold capacity in such a data-driven setting, the most straightforward way is to numerically downsample the number of neurons and estimate the critical number of neurons required for manifold separation.

#### Simulation capacity

Given some finite data manifolds, (numerically) find the smallest number of neurons *N* ^*^ in which the manifolds are still separable^4^. The manifold capacity is then defined as *P*_data_*/N* ^*^.

In a theoretical setting, *P* and *N* are free parameters that can be arbitrarily chosen and sent to infinity. This allows previous works to characterize manifold capacity analytically in a mathematical setting, a.k.a., a mean-field model.

#### Mean-field capacity

Given some finite data manifolds, use the shapes and correlations among them to generate *P* “copies” of mean-field manifolds in an *N*-dimensional space. The manifold capacity is then defined as the largest ratio *P/N* whereas these mean-field manifolds are separable.

These two ideas lead to two (families of) definitions of manifold capacity: simulated capacity *α*_sim_ and mean-field capacity *α*_mf_. While *α*_sim_ is intuitive and easier to be understood, it is relatively mathematically ad hoc. In contrast, *α*_mf_ is mathematically well-defined and is analytically tractable, however, the choice of mathematical assumptions determine how well *α*_mf_ can approximate *α*_sim_. Indeed, previous works [20, 21] proposed mean-field models that incorporate the properties of data manifolds into mean-field models and use mean-field capacity (and its effective geometric measures) as a surrogate for the data manifolds. Nevertheless, these prior mean-field models [20, 21] rely on two critical simplifying mathematical assumptions: (i) Gaussian correlations between manifolds and (ii) random task labels. These assumptions often fail with biological datasets, as they ignore higher-order correlations and task-specific structures, limiting their applicability (see Supplementary Figure S3).

In this work, we directly analyze the simulation capacity and derive an analytical formula for *α*_sim_. We mathematically prove that our formula well-estimate *α*_sim_ and empirically demonstrate on 7 biological datasets, synthetic models, and artificial neural networks. In the rest of this section, we first introduce the more intuitive definition of manifold capacity via a *simulation model* in Supplementary Information S1.1.

**Table 1:**
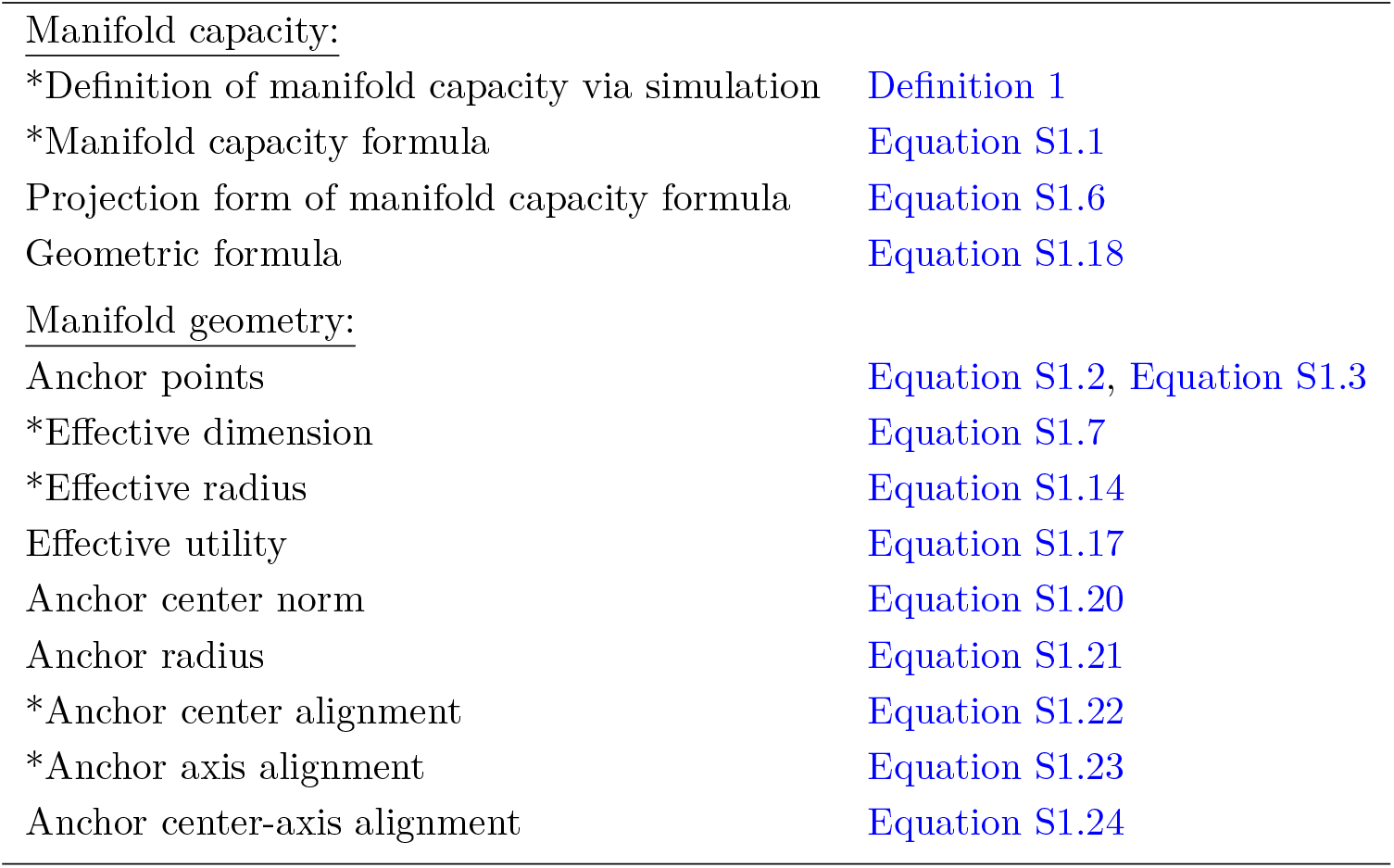
Look-up table for key concepts and definitions in manifold capacity theory. Those marked with an asterisk are covered in the main text while the rest are discussed in the supplementary materials.

Manifold capacity:
|  |  |
| --- | --- |
| *Definition of manifold capacity via simulation | Definition 1 |
| *Manifold capacity formula | Equation S1.1 |
| Projection form of manifold capacity formula | Equation S1.6 |
| Geometric formula | Equation S1.18 |
| Manifold geometry: |  |
| Anchor points | Equation S1.2, Equation S1.3 |
| *Effective dimension | Equation S1.7 |
| *Effective radius | Equation S1.14 |
| Effective utility | Equation S1.17 |
| Anchor center norm | Equation S1.20 |
| Anchor radius | Equation S1.21 |
| *Anchor center alignment | Equation S1.22 |
| *Anchor axis alignment | Equation S1.23 |
| Anchor center-axis alignment | Equation S1.24 |

**Table 2:**
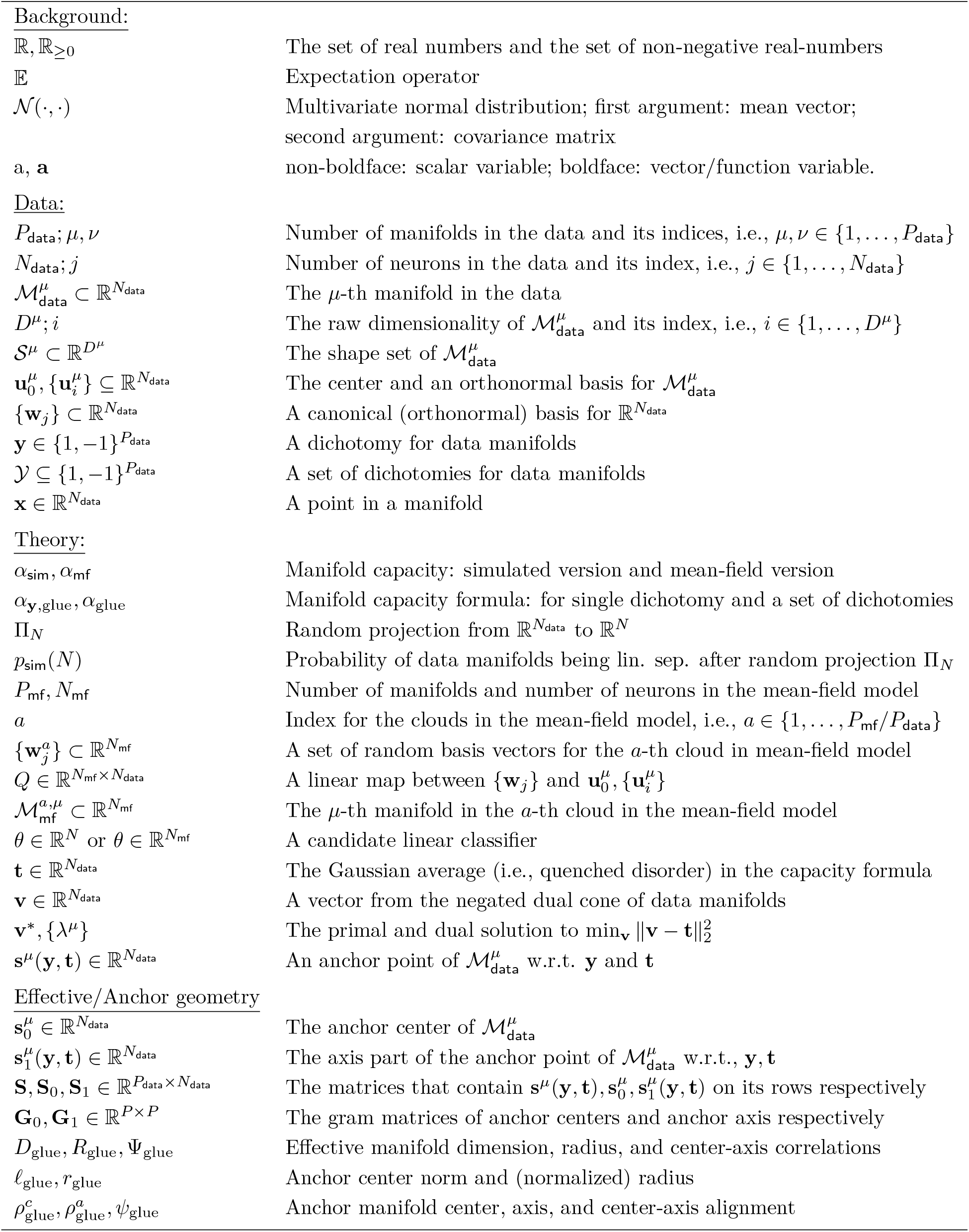
Notations.

#### S1.1 Definition of manifold capacity in a simulation model

Given a collection of (finite) data manifolds 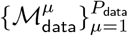. Let 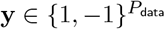 be a dichotomy for the data manifolds. For each *N* ∈ {1, 2, …, *N*_data_}, which should be thought of as the size of a subpopulation of neurons, define

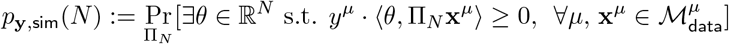

where Π_*N*_ is a random projection from 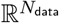 to ℝ^*N*^. Namely, *p*_sim_(*N*) is the probability of the data manifolds being linearly separable with respect to dichotomy **y** after a random projection to an *N*-dimensional subspace.

##### Definition 1

(Manifold capacity, simulated version). *Given a collection of (finite) data manifolds* 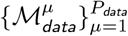 *and the above simulation model. Let* 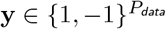 *be a dichotomy. The simulated manifold capacity is defined as*

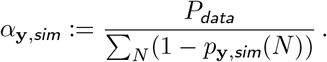

Intuitively^5^, the denominator 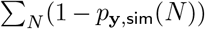 can be thought of as the *expected number of neurons* required for being able to linearly separate the manifolds according to **y**.

##### A remark on dichotomy

For the simplicity of the presentation, we focus on defining manifold capacity for a single dichotomy vector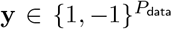. Every discussion can be directly generalized to a set of dichotomies 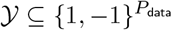 by taking the geometric average of manifold capacity over 𝒴.

#### S1.2 Manifold capacity formula

We derive a formula for *α*_sim_ as follows.

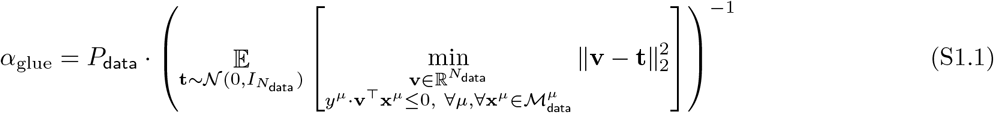

where 𝒩 (0, *I*_data_) denotes the multivariate isotropic normal distribution. Also see Supplementary Figure S3 for an empirical check on all the datasets we analyzed (and a comparison with the formulas proposed by previous works [20, 21]).

The above capacity formula (Equation S1.1) has two advantages: (i) it is computationally much cheaper to estimate than the simulation model (Definition 1), see Algorithm 1 for a pseudocode; (ii) the formula hints a route to define manifold geometric measures as illustrated in the subsequent subsections.

#### S1.3 Anchor points

To extract manifold geometry from the capacity formula (Equation S1.1), we follow the idea from [20] to rewrite the term^6^ with “min” in Equation S1.1 by the KKT condition [66, Chapter 5]. Concretely, let 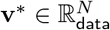 denote the optimizer of 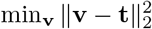, there exists ***λ***^*^ = {***λ***^*,*µ*^} (where ***λ***^*,*µ*^ : ℳ^*µ*^ → ℝ_≥0_) such that the following equations hold:

- (Primal feasibility) *y*^*µ*^ · ⟨**v**^*^, **x**^*µ*^⟩ ≥ 0 for all *µ* ∈ [*P*] and **x**^*µ*^ ∈ ℳ^*µ*^.
- (Dual feasibility) *λ*^*,*µ*^(**x**^*µ*^) ≥ 0 for all *µ* ∈ [*P*] and **x**^*µ*^ ∈ ℳ^*µ*^.
- (Complementary slackness) *λ*^*,*µ*^(**x**^*µ*^) · **y**^*µ*^ · ⟨**v**^*^, **x**^*µ*^⟩ = 0 for all *µ* ∈ [*P*] and **x**^*µ*^ ∈ ℳ^*µ*^.

**Figure S3:**
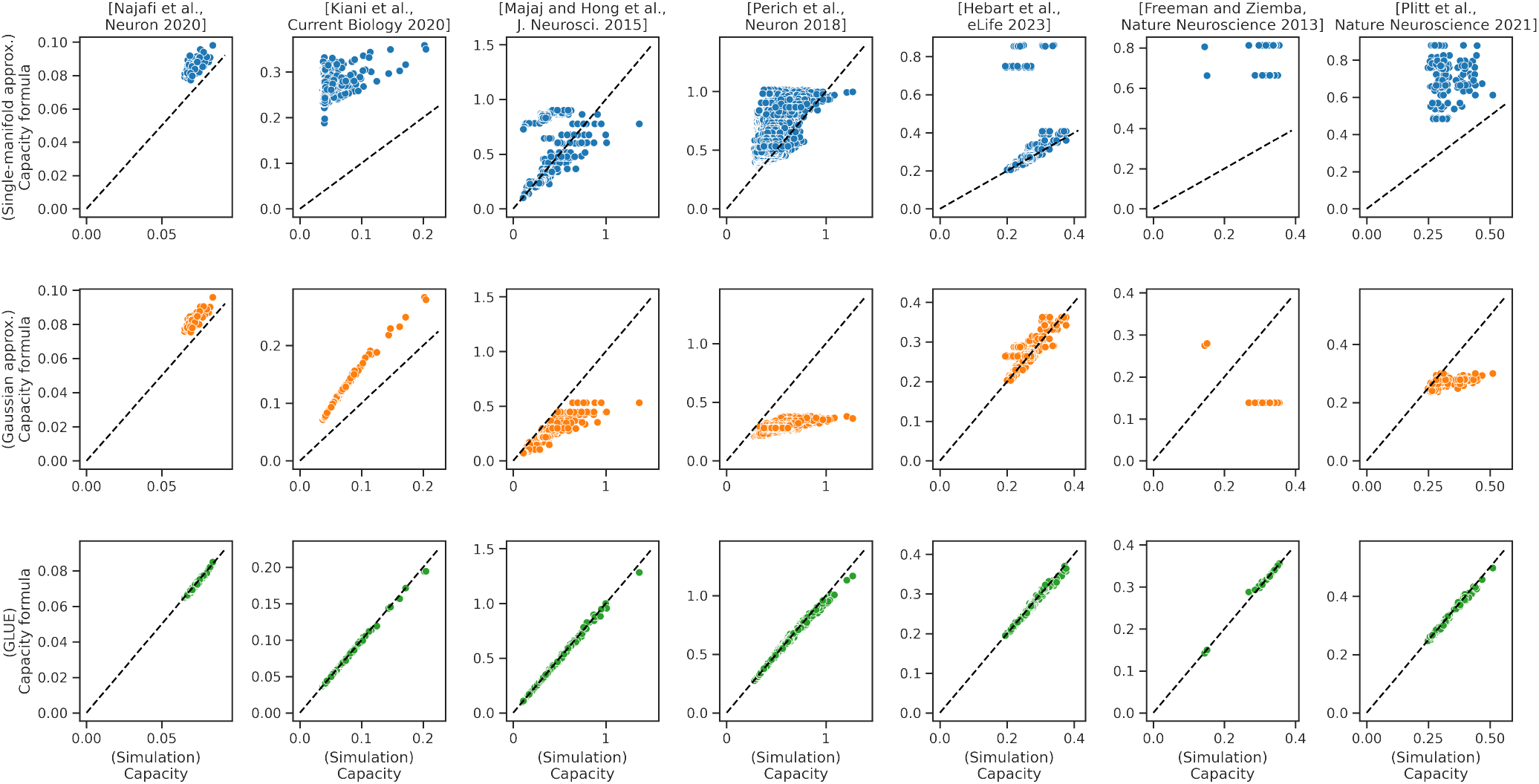
Comparison between simulation capacity and the estimate via capacity formulas. Each column corresponds to a dataset analyzed in this paper. The x-axis is the capacity value estimated from numerical simulation as described in Definition 1 and the y-axis is the capacity value estimated from a manifold capacity formula. The first and the second row correspond to the manifold capacity formula from the previous works [20, 21] respectively. The last row corresponds to the capacity formula derived in this paper, i.e., Equation S1.1.

We define the anchor points for each data manifolds as

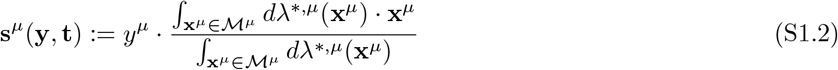

where we treat the dual function of the *µ*-th manifold, ***λ***^*,*µ*^ as a measure over ℳ ^*µ*^. For data analysis where the manifolds are point clouds 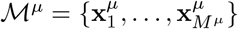,Equation S1.2 becomes

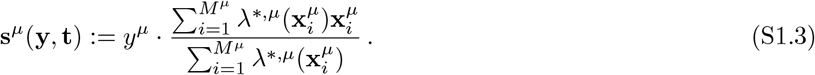

Anchor points connect to manifold capacity via a simple analytical formula. In particular, by putting Equation S1.1 and the KKT conditions ([66], Chapter 5) together, we have

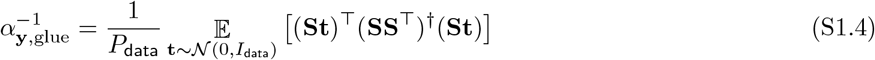

where the matrix 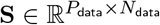 contains the anchor points **s**^*µ*^(**y, t**) on its rows and † denotes pseudo-inverse. Note that as **S**^⊤^(**SS**^⊤^)^†^**S** is performing the orthogonal projection to the subspace spanned by the rows of **S**. Namely,

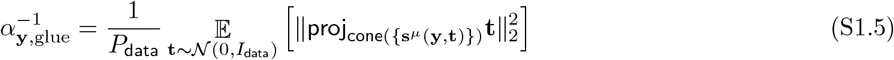

where proj is the projection operator and cone({**s**^*µ*^}) = {∑ _*µ*_ *λ*^*µ*^**s**^*µ*^ : ∀*µ, λ*^*µ*^ ≥ 0} is the convex cone spanned by {**s**^*µ*^}.

When considering a set of dichotomies of interest, 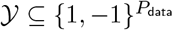,we have

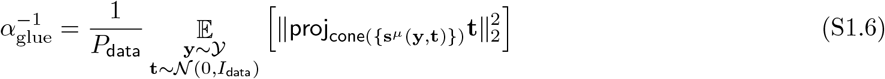

where **y** ∼ **𝒴** denotes (uniformly at random) sampling **y** from 𝒴. In the rest of this section, we use 𝔼_**y**,**t**_ to abbreviate the expectation term in the above formula when the context is clear.

As a remark, Equation S1.4 and Equation S1.5 are mathematically the same. The reason we present both here is that the former is more convenient for implementation and the latter is easier for intuitive reasoning.

#### S1.4 Effective manifold geometric measures from the capacity formula

##### The center and axis part of anchor points

Next, we again follow the intuition from [20] and decompose the anchor points **s**^*µ*^(**y, t**) _*µ*_ into the longitudinal component (i.e., manifold center) s^μ^_0_ and the intrinsic component 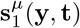, i.e.,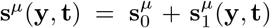 where *µ* is the manifold this anchor point corresponds to. Concretely, 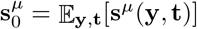 is the mean of the anchor points coming from the *µ*-th manifold and 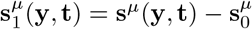.

To compute effective geometric measures in practice, we can adopt an equivalent form of capacity formula, Equation S1.4, which explicitly expresses the projection operator as some linear operators consisting low-order moments of the anchor points. Concretely, we can also decompose the matrix **S** = **S**_0_ + **S**_1_ where **S**_0_ contains 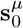 on its rows and **S**_1_ contains 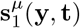 on its rows.

##### Effective manifold dimension

To gain some intuition before formally introduce our definition of effective manifold dimension, think about the case of projecting a random (Gaussian) vector in ℝ^*N*^ to a *D*-dimensional subspace. Let {**e**_1_, …, **e**_*D*_} be an orthonormal basis for this subspace. It is not hard to see that the expected value of the square of the projection is 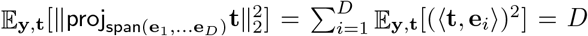.This inspires us to use the expected value of the square of the projection of a random (Gaussian) vector **t** to measure the (average) dimension of manifolds.

Moreover, only the axis part of the anchor points are reacting the in-category variation. Hence, we measure the (average) degree of freedom (i.e., number of directions) within manifolds by considering the projection of **t** to the cone spanned by the axis part of the anchor points, i.e.,

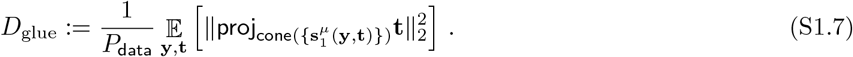

To empirically estimate the effective dimension, we can use the fact that 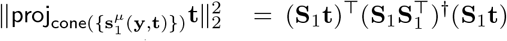 and the following formulas (which are implemented in the pseudocode in method):

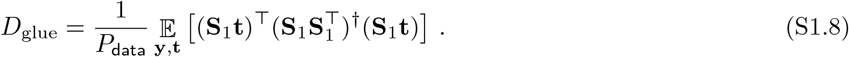

See Figure S6c for an example of how *D*_glue_ faithfully tracks the raw dimensionality of uncorrelated Gaussian point clouds.

##### Effective manifold radius

To motivate the definition of effective manifold radius, let us consider the case where there is only one manifold (i.e., the setting in [20]). In this case, the projection of **t** onto the cone spanned by the anchor point **s**(**t**) can be expanded as follows.^7^

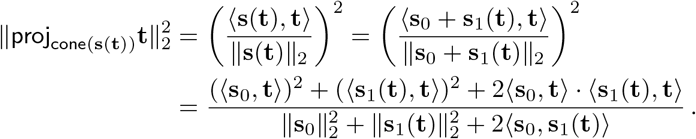

The key idea in [20] is to approximate the above by ⟨**s**_0_, **t**⟩ ≈ 0, ⟨**s**_0_, **t**⟩ · ⟨**s**_1_(**t**), **t**⟩ ≈ 0, ⟨**s**_0_, **s**_1_(**t**)⟩ ≈ 0, and consider

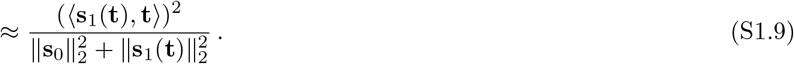

Note that in this setting, the effective dimension term is 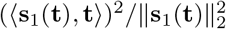 and the above can be further rear-ranged as

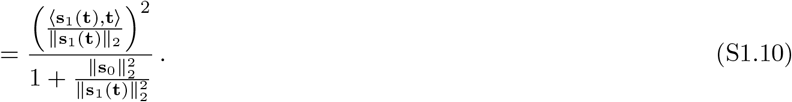

The approximation in Equation S1.9 (for each **t**) inspired [20] to define their effective radius as

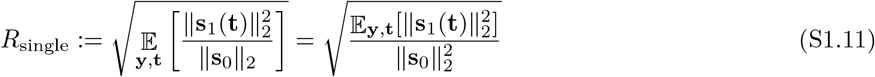

with the following geometric approximation to manifold capacity:

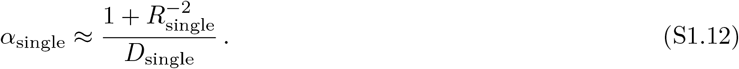

In this work, we start with extending the above single-manifold setting from [20] to multi-manifold case as follows. First, we consider the following multi-manifold version of Equation S1.9

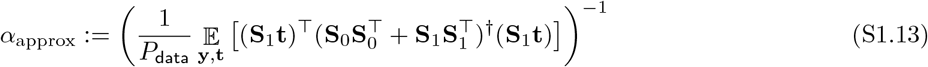

where **S**_0_, **S**_1_ are the matrices with the center and axis part of the anchor points on their rows respectively.

Next, consider the identity 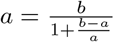 used in Equation S1.10 and set 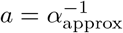 and *b* = *D*_glue_ to figure out what the radius term should be. Specifically, we define

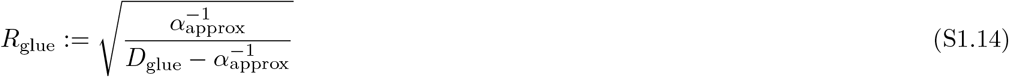

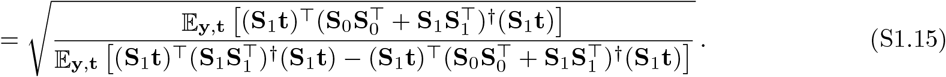

See Figure S6b for an example of how *R*_glue_ faithfully tracks the raw radius (normalized by the center norm) of uncorrelated Gaussian point clouds.

Note that from the above definition, we have that 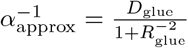 and similar to Equation S1.12, we have that

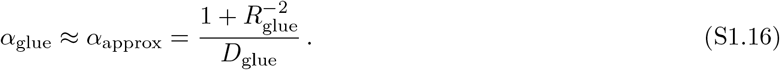

##### Effective utility rate

We do not stop at the geometric approximation for capacity formula via the center-axis decoupling idea from [20]. Specifically, we observe that the geometric approximation always *overestimate* the capacity (Supplementary Figure S4), i.e., 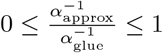.This motivates us to define an *effective utility rate* as

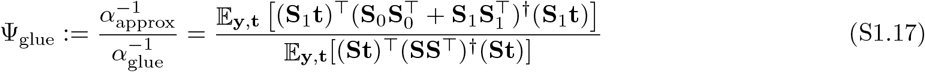

which is a quantity between 0 and 1 (when the inequality 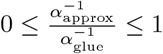 holds). Moreover, the larger the Ψ _glue_ is, the less effect of correlations has on the capacity. Together, we have an **exact** geometric formula for the manifold capacity as follows.

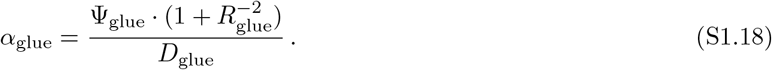

**Figure S4:**
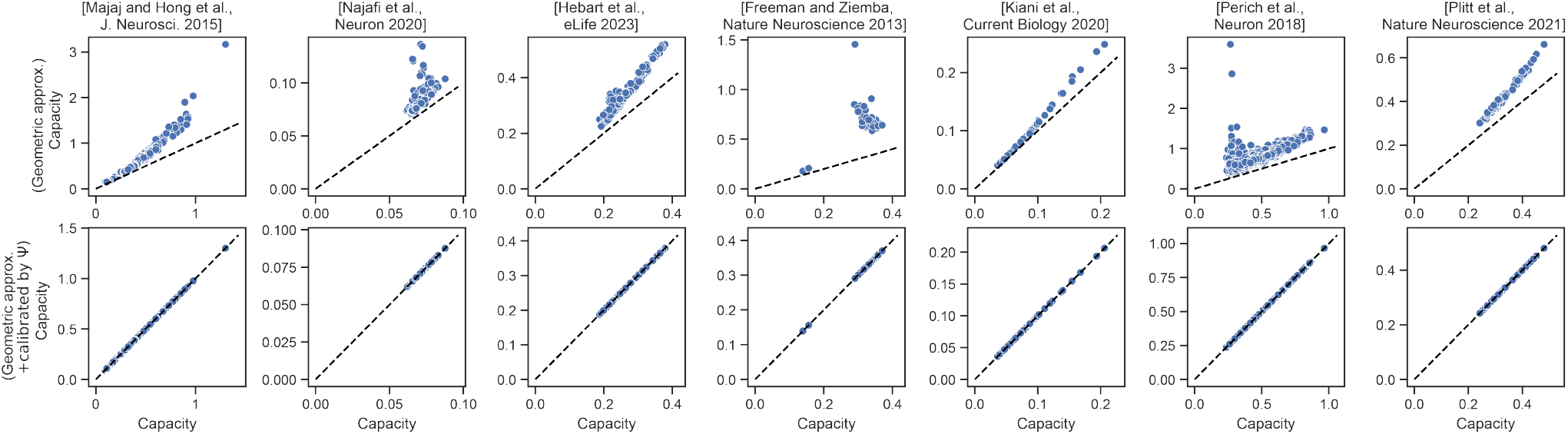
Comparison of simulation capacity, our capacity formula, and our geometric approximation. Each column corresponds to a dataset analyzed in this paper. The x-axis is the capacity value estimated from numerical simulation, as described in Definition 1. The y-axis of the first row corresponds to the geometric approximation, i.e., *α*_approx_ (Equation S1.13). The y-axis of the second row corresponds to the capacity formula, i.e., *α*_glue_ (Equation S1.6). Notice that, as shown in the first row, the geometric approximation always overestimates the capacity value.

We remark that we didn’t present the result on effective utility rate in the main text for simplicity purpose. In the single manifold case, the effective utility rate becomes,

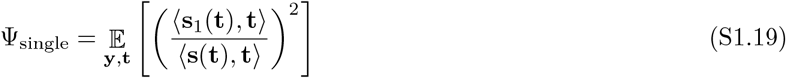

which intuitively is the fraction of “error” coming from the axis-part of the anchor points. Also, unlike effective radius and dimension, effective utility rate behaves less like a geometric measure, but more like an efficiency measure for capacity itself—intuitively *α*_approx_ can be thought of as the best possible capacity to achieve with a certain geometric constraint and hence the ratio *α*_glue_*/α*_approx_ can be thought of as how close the manifold capacity is to the upper bound. We post it as an interesting future direction to explore the meaning and consequence of effective utility rate.

To summarize, so far we have derived three effective manifold geometric measures, effective dimension *D*_glue_, effective radius *R*_glue_, and effective utility rate Ψ_glue_, in which a simple function of them gives rise to the manifold capacity as described in Equation S1.18. See Supplementary Figure S4 for an empirical check on Equation S1.17.

#### S1.5 Anchor manifold alignment measures

In the rest of this section, we are going to see some more manifold alignment measures defined via low-order statistics of the anchor points. Unlike the effective geometric measures, they do not analytically connect to the manifold capacity via a formula. However, they can be regarded as a replacement to the analogous correlation measures conventionally defined via lower-order statistics of the data points, and can be used as summary statistics for data analysis.

- Anchor center norm:

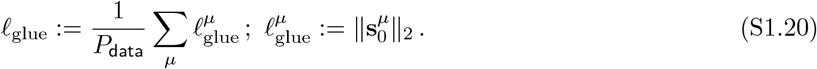 For simplicity of the presentation, we did not put the results on anchor center norm in the main figures. Nevertheless, for future applications of GLUE, anchor center norm can potentially be an important factor that explains the changes in manifold organization and capacity.
- Anchor (normalized) radius:

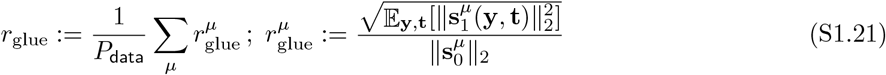 Note that the anchor (normalized) radius defined here is different from the effective radius *R*_glue_ as defined in Equation S1.14. Here *r*_glue_ and 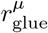 do not have a simple analytical connection to manifold capacity, while *R*_glue_ is analytically linked to *α*_glue_ through Equation S1.18. For simplicity of the presentation, we did not put the results on anchor radius in the main figures. Nevertheless, for future applications of GLUE, anchor radius can potentially augment the intuition from other effective geometric measures for understanding the changes in manifold organization and capacity.
- Anchor center alignment:

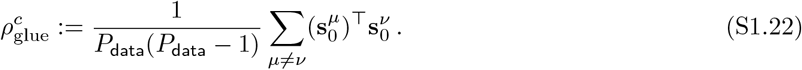 See Figure S6d for an example of how 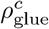 faithfully tracks the raw center correlations of Gaussian point clouds.
- Anchor axis alignment:

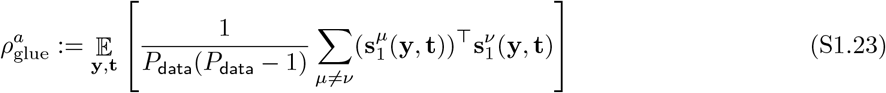

where the superscript for 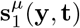 indicates the index of the manifold that anchor point corresponds to. See Figure S6e for an example of how 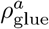 faithfully tracks the raw axis correlations of Gaussian point clouds.
- Anchor center-axis alignment:

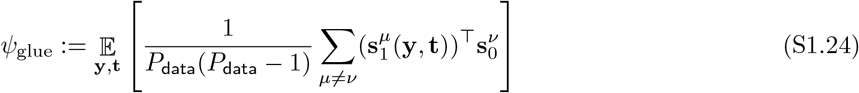 For simplicity of the presentation, we did not put the results on anchor center-axis alignment in the main figures. Nevertheless, for future applications of GLUE, anchor center-axis alignment can potentially be an important factor that explains the changes in manifold organization and capacity.

We remark that we didn’t present the result on anchor center norm, radius, and center-axis alignment in the main text for simplicity purpose. Also, we remark that the effective/anchor geometry of manifold capacity theory can go beyond the metrics introduced here, and we leave it as an interesting future direction to explore other possible effective/anchor geometric measures.

### S2 Toy Examples for Intuition Building

To build intuitions on effective manifold geometric measures as well as interpret the results on real datasets, it is instructive to examine toy examples where the underlying ground truth is known. In this section, we provide three toy examples for the readers to build up intuitions on interpreting and understanding manifold capacity and its effective geometric measures. First, in Supplementary Information S2.1, we compare manifold capacity with conventional Receiver Operating Characteristic (ROC) for single-neuron selectivity. Next, in Supplementary Information S2.2, we consider synthetic manifolds generated from Gaussian mixture models, which should be regarded as idealized models for real neural manifolds. By systematically sweeping through various latent variables, we can learn how the organization of Gaussian manifolds affect the effective geometric measures. Hence, this serves as a dictionary for mapping the effective geometric values from real data back to an idealized latent space. Finally, in Supplementary Information S2.3, we illustrate the equivalence between manifold capacity and the geometry of the cone of linear classifiers.

#### S2.1 Comparison to single-neuron selectivity analysis

In Supplementary Figure S5, we compare the ROC analysis for single-neuron selectivity to manifold capacity analysis. The two methods bear surprising similarity, and hence the readers can easily draw intuition on manifold capacity from analogous concepts in the ROC analysis.

**Figure S5:**
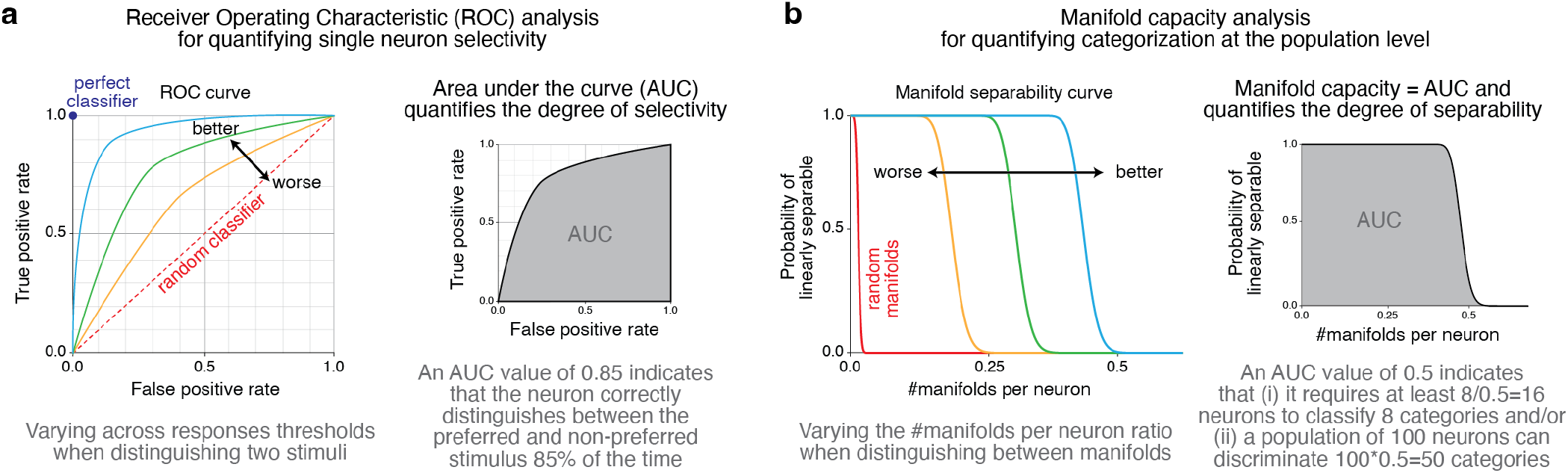
Comparison between single-neuron selectivity analysis and manifold capacity. **a**, ROC analysis for single-neuron selectivity, where the degree of selectivity is quantified by the AUC. **b**, Manifold capacity analysis for categorization at the population level, where the degree of separability/decodability is quantified by the AUC as well, which coincides with manifold capacity.

##### Single-neuron selectivity

The Receiver Operating Characteristic (ROC) curve for a single neuron depicts the relationship between the True Positive Rate and the False Positive Rate across varying response thresholds when distinguishing between two stimuli (Figure S5a). The Area Under the Curve (AUC) of the ROC curve serves as a quantitative measure of the neuron’s selectivity, reflecting its ability to discriminate between the two stimuli. Specifically, the AUC represents the probability that a neuron, using a threshold-based decision rule, will correctly identify a preferred stimulus over a non-preferred one. For instance, an AUC of 0.85 indicates an 85% probability that the neuron’s response to a randomly selected preferred stimulus exceeds its response to a randomly selected non-preferred stimulus. This metric provides a robust assessment of single-neuron discriminative performance, independent of the chosen threshold.

##### Manifold capacity

The manifold load curve for a population of neurons illustrates the relationship between manifold load—defined as the number of categories normalized by the number of neurons—and the probability of successfully classifying population responses from different categories (Figure S5b). Manifold capacity is quantified as the Area Under the Curve (AUC) of the manifold load curve, thereby encapsulating the population’s ability to discriminate between categories. Specifically, manifold capacity indicates the maximum normalized number of categories that the neural population can reliably distinguish using a linear readout. For example, a manifold capacity of 0.25 implies that to distinguish between 8 categories, approximately 32 neurons are required (since 8 categories / 0.25 = 32 neurons). Conversely, if a population comprises 100 neurons, it can differentiate among 25 categories (100 neurons × 0.25 = 25 categories). In comparison, a system with an manifold capacity of 0.1 would necessitate 2.5 times more neurons to achieve the same level of classification performance. This metric provides a scalable measure of neural population efficiency in encoding and discriminating complex categorical information.

#### S2.2 A correlated Gaussian point cloud manifold model

In item S6 we consider point cloud manifolds generated from a Gaussian model with eight latent parameters: number of neurons (*N*), number of manifolds (*P*), number of points per manifold (*M*), raw radius (*R*), raw dimension (*D*), center correlations (*ρ*_*c*_), axis correlations (*ρ*_*a*_), and center-axis correlations (*ψ*). See a pseudo-code for the Gaussian model below as well as Figure S6a for a schematic illustration.

- Parameters: *N* ∈ ℕ (ambient dimension); *P* ∈ ℕ (number of manifolds); *D* ∈ ℕ (manifold dimension); *M* ∈ ℕ (number of points per manifold); *ρ*_*a*_ ∈ [0, 1) (axes correlations); *ρ*_*c*_ ∈ [0, 1) (center correlations); *ψ* ∈ ℝ_≥0_ (center-axes correlations).
- Procedures:
  1. Randomly sample *P* manifold centers 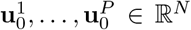 independently from the Gaussian distribution with mean 0 and variance 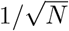 for each coordinate.
  2. Let 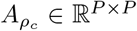 be the square root of the matrix with diagonal being 1s and off-diagonal being 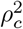.Set the vectors 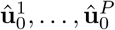 to be the columns of 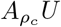 where *U* contains 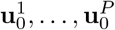 in its columns.
  3. For each *µ* = 1, …, *P*, randomly sample *D* vectors 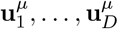 from the Gaussian distribution with mean 0 and variance 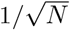 for each coordinate.
  4. Let 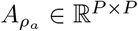 be the square root of the matrix with diagonal being 1s and off-diagonal being 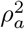.For each *i* = 1, …, *D*, set the vectors 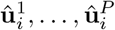 to be the columns of 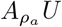 where *U* contains 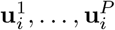 in its columns.
  5. Define the *µ*-th manifold to be the point cloud containing the following vectors: 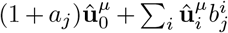 for each *j* = 1, …, *M* where *a*_*j*_ is independently sampled from the Gaussian distribution with mean 0 and variance *ψ*^2^ and 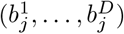 is uniformly sampled from the unit sphere in ℝ^*D*^.

**Figure S6:**
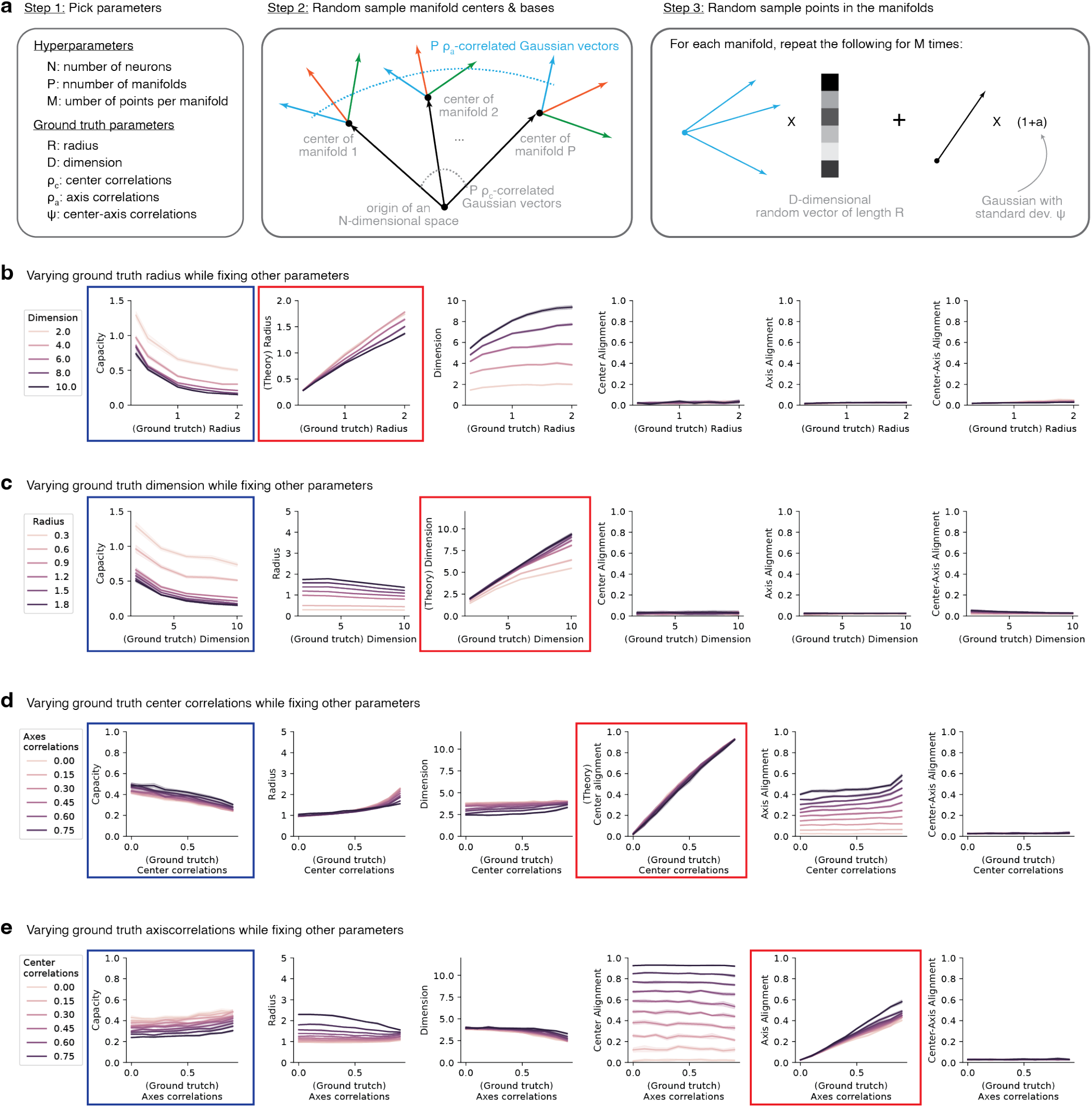
GLUE analysis on synthetic manifolds via a Gaussian point cloud manifold model. **a**, Schematic illustration of the model. **b**-**e**, Results of GLUE analysis. See Supplementary Information S2.2 for experimental details.

Now, we consider two settings of the above-mentioned synthetic model. First, we consider uncorrelated Gaussian point clouds (i.e., *ρ*_*c*_ = *ρ*_*a*_ = *ψ* = 0) and investigate the effect of *ground truth* radius and dimension on manifold capacity and effective geometric measures. In this case, we found that both effective radius and effective dimension faithfully tracks the ground truth value. Next, we fix the raw radius and dimension, and vary the center and axis correlations in the model. We found that both the effective center alignment and effective axis alignment track the corresponding ground truth value as well.

##### Uncorrelated Gaussian point clouds and manifold radius and dimension

We pick *N* = 1000, *P* = 2, *M* = 200, *ρ*_*c*_ = *ρ*_*a*_ = *ψ* = 0, vary *R* between 0.3 and 2, and vary *D* between 2 and 10 in Figure S6b and Figure S6c with 50 repetitions for each parameter setting.

In Figure S6b, one can see that the increase of raw radius *R* indeed hurts the manifold capacity, and the effective radius *R*_glue_ quantitatively tracks the value of *R*. In particular, *R*_glue_ approximates *R* well when *D* is small. We remark that in general *R*_glue_ would not be exactly the same as *R* due to finite size effect—when the raw dimension is large and the number of points per manifold (*M*) is small, the Gaussian point clouds from our synthetic model would be far away from being a sphere and hence the effective radius *R*_glue_ in general would be smaller than the raw radius *R*.

In Figure S6c, one can see that the increase of raw dimension *D* indeed hurts the manifold capacity, and the effective dimension *D*_glue_ quantitatively tracks the value of *D*. In particular, *D*_glue_ approximates *D* well when *R* is large. We remark that in general *D*_glue_ would not be exactly the same as *D* due to finite size effect—when the raw radius is small and the number of points per manifold (*M*) is small, the Gaussian point clouds from our synthetic model would be far away from being a sphere nor a subspace and hence the effective dimension *D*_glue_ in general would be smaller than the raw dimension *D*.

Lastly, we remark that all the three effective alignment measures correctly stay close to 0 (they would not be exactly 0 due to finite-size effect) in this uncorrelated Gaussian point clouds setting.

##### Correlated Gaussian point clouds and alignment measures

We pick *N* = 1000, *P* = 2, *M* = 200, *R* = 1, *D* = 4, and vary *ρ*_*c*_, *ρ*_*a*_, *ψ* between 0 and 0.8 in Figure S6d and Figure S6e with 50 repetitions for each parameter setting.

In Figure S6d, one can see that the increase of raw center correlations *ρ*_*c*_ indeed hurts the manifold capacity, and the effective center alignment 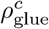 quantitatively tracks the value of *ρ*_*c*_. Noticeably, we found that the effective radius *R*_glue_ increases along with center correlations. Intuitively, this is because the anchor point distribution becomes biased toward the global center (i.e., the center of the center of manifolds). See also Figure S7g. Similarly, the effective axis alignment 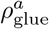 increases along with the raw center correlations when the raw axis correlations are high.

In Figure S6e, one can see that the increase of raw axis correlations *ρ*_*a*_ indeed helps the manifold capacity, and the effective axis alignment 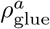 quantitatively tracks the value of *ρ*_*a*_. Noticeably, we found that the effective dimension *D*_glue_ decreases along with axis correlations. Intuitively, this is because the anchor point distribution becomes biased toward the shared variation subspace across manifolds. See also Figure S7h. Similarly, the effective radius *R*_glue_ decreases along with the raw axis correlations when the raw radius is high.

#### S2.3 The cone of linear classifiers

It turns out that the manifold capacity is related to the *statistical dimension* [67] of the convex cone that contains all the linear classifiers. This connection provides an alternative angle to intuitively understand the interaction between manifold capacity and its effective geometric measures. In the following, we will provide a simplified illustration of this connection. For interested readers, see [68] for a more comprehensive coverage on convex cone geometry.

Given a collection of (finite) data manifolds 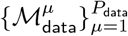 and a dichotomy 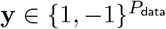 for the data manifolds. The cone of linear classifiers for 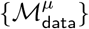 with respect to **y** is

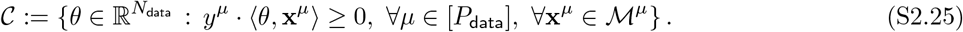

See Figure S7a for an example of classifying two manifolds.

It is not difficult to see that *𝒞* is a convex cone, i.e., (i) ∀*θ* ∈ 𝒞, *aθ* ∈ *𝒞* for all *a*≥ 0 and (ii) ∀*θ, θ*^′^, ∈ *𝒞aθ* + *a*^′^*θ*^′^*𝒞* for all *a, a*^′^ ≥ 0 with *a* + *a*^′^ = 1. While there is a huge and rich theory for the geometry of convex cones [67], here we focus on a concept called *statistical dimension* (see Definition 2.2 and Proposition 5.12 of [68]). Intuitively, the statistical dimension of a convex cone is analogous to the dimension of linear subspaces. In fact, the statistical dimension of a *D*-dimensional subspace is *D*. So, we encourage the readers to think about the statistical dimension of C as a measure of how large C is^8^.

Now, we state the connection between manifold capacity of 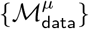 with respect to **y** and the statistical dimension 𝒞 of C as follows.

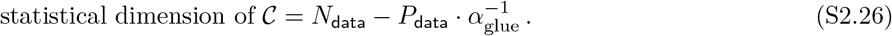

We omit the proof of Equation S2.26 for two reasons: (i) it is an immediate corollary of [68, Proposition 5.12] and (ii) hence the bulk of the proof is about defining statistical dimension. We encourage interested readers to read [68].

With Equation S2.26, we immediately know that manifold capacity *α*_glue_ increase (or decreases) along with the statistical dimension of the cone of linear classifiers 𝒞. See Figure S7b for an example. As a consequence, when thinking about how manifold organization affects manifold capacity, one can freely switch to thinking about the size of the cone of linear classifiers. In the rest of this subsection, we will use such connection to present several useful intuitions for manifold capacity and its effective geometric measures.

##### The distribution of anchor points connects manifold geometry to manifold capacity

While manifold capacity quantifies the degree of untangledness of manifolds, distinct manifold organizations might exhibit the same capacity value (see Figure S7c for an example). As mentioned in the main text, a high capacity value can either be achieved by having small manifold radius or having small manifold dimension. To see the underlying geometric composition of the manifold capacity value, we can look at the difference in the distribution of anchor points (see Equation S1.2 and Equation S1.3). While in the previous section we introduce anchor points in a relatively abstract manner, here using the intuitive picture of cone of linear classifiers we can easily understand anchor points as the worst-points of a random non-classifier (see Figure S7d). With this intuition in mind, we can then easily understand how different manifold organizations achieve the same capacity value in different ways. For example, in Figure S7e, the anchor points distribution will be quite different for the two cases—for the example on the left, the anchor points will be more concentrated to the center (i.e., small radius) and the centers of the anchor points will be closer with each other (i.e., higher center correlations); for the example on the right, the anchor points will be less concentrated to the center (i.e., larger radius) while the centers of the anchor points will be more separated from each other (i.e., small center correlations).

##### How manifold geometry affects manifold capacity

As mentioned in Figure S7b, the manifold capacity value increases when the size/volume (or more precisely, the statistical dimension) of the cone of linear classifiers increases. Using this intuitive picture, we can easily understand how a decrease in radius, a decrease in dimension, a decrease in center alignment, or an increase in axis alignment makes the size/volume of the cone larger, hence making the capacity higher (Figure S7f).

##### Interaction/Correlation between effective geometric measures

Once we are familiar with how different manifold geometric properties affect manifold capacity, as shown in Figure S7f, we can further understand how the effective geometric measures interact and correlate with one another. We note that the effective geometric measures defined and used in GLUE are statistics (of anchor points) that summarize the average properties across all data manifolds. That being said, the cone picture discussed in this section is not perfectly identical to the effective geometric measures. Nevertheless, at a high level, the intuition can be shared.

**Figure S7:**
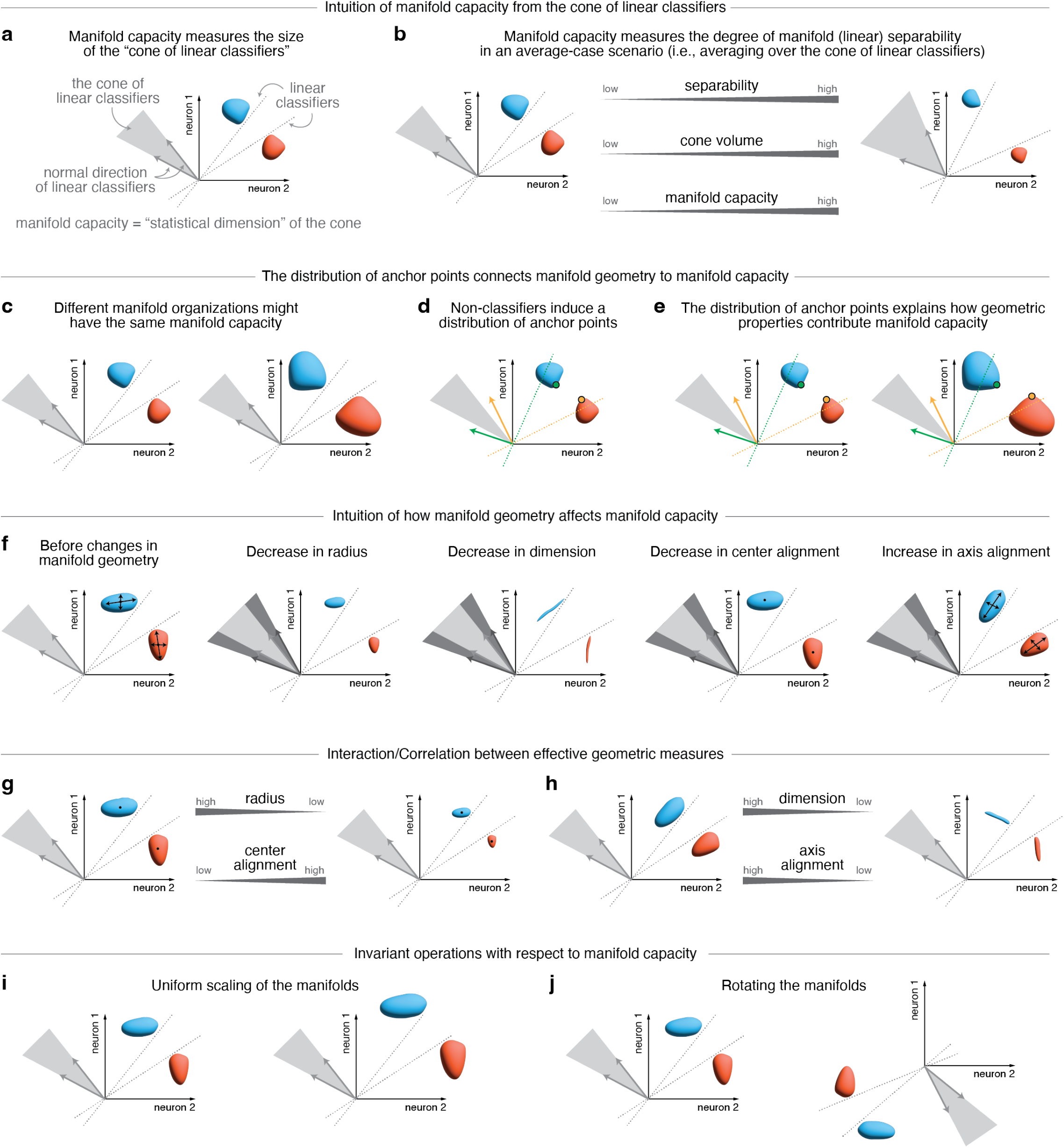
Intuitive pictures for manifold capacity and its effective geometric measures.

First, there is an interaction between radius and center alignment. As shown in Figure S7g), the cone of linear classifiers could stay the same while simultaneously shrinking the manifold radius and increasing the manifold center alignment. Indeed, our effective radius *R*_glue_ captures this interaction and reflect in the geometric formula for manifold capacity (Equation S1.18). Recall that there is no explicit term on center alignment in Equation S1.18 so its effect has been reflected in the effective radius term. For example, in the correlated Gaussian point cloud model, we found that large center correlations in the generative model would be reflected by an increase in effective radius (Figure S6d).

Next, there is an interaction between dimension and axis alignment. As show in Figure S7h, the cone of linear classifiers could stay the same while simultaneously shrinking the manifold dimension and decreasing the manifold axis alignment. Indeed, our effective dimension *D*_glue_ captures this interaction and reflect in the geometric formula for manifold capacity (Equation S1.18). Recall that there is no explicit term on axis alignment in Equation S1.18 so its effect has been reflected in the effective dimension term. For example, in the correlated Gaussian point cloud model, we found that large axis correlations in the generative model would be reflected by an decrease in effective dimension (Figure S6e).

##### Invariant operations with respect to manifold capacity

Finally, the cone of linear classifiers and the manifold capacity stay invariant under some operations. For example, in Figure S7i, the cone and capacity stay invariant when uniformly scaling the manifolds by some non-zero multiplicative factor. This is the reason why our effective radius *R*_glue_ is normalized by the center norm. Namely, *R*_glue_ is the corresponding invariant quantity with respect to uniform scaling. Similarly, in Figure S7j, the cone and capacity stay invariant under some (subspace) rotations. Intuitively, the effective dimension *D*_glue_ is related to the size of the invariant rotation group, and we leave it as a mathematical open problem for future theoretical investigations to make this connection rigorous.

### S3 Data analysis

In this section, we provide our interpretations on the results we got from the GLUE analysis on each dataset.

**Table 3:**
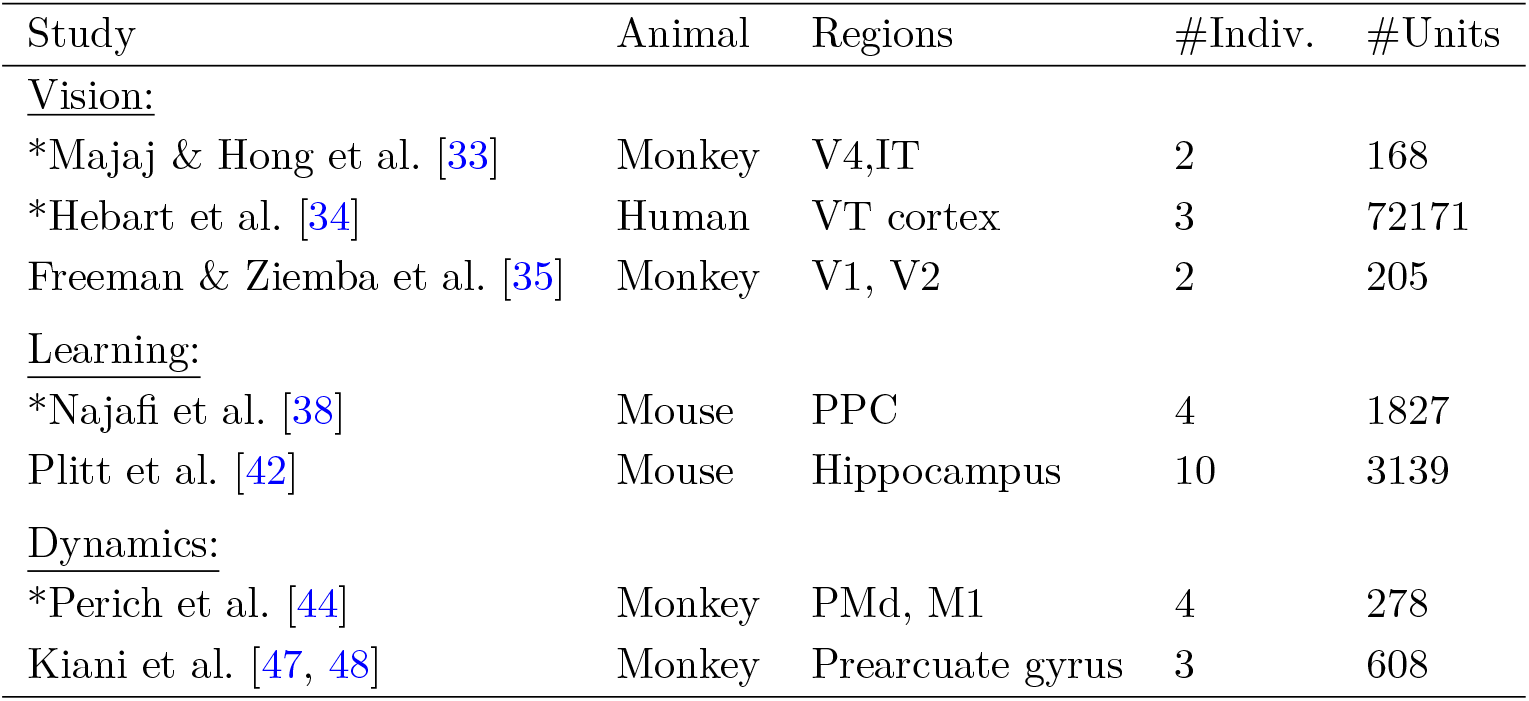
Datasets analyzed by GLUE in this paper. Those marked with an asterisk are covered in the main text while the rest are discussed in the supplementary materials.

#### S3.1 Monkey early vision dataset

In Supplementary Figure S8 we analyze the dataset from [35] on a monkey vision task with electrophysiological recordings in V1 and V2. See Methods for more analysis details.

In the GLUE analysis, we see an increment of manifold capacity from pixel level to V1 and to V2 in both texture manifolds and noise manifolds (Supplementary Figure S8b). The interpretation of capacity here corresponds to the encoding efficiency of a representation for distinguishing among the different texture patterns (or their noisy version). In particular, the capacity growth from V1 to V2 is mainly driven by effective radius, effective dimension, and effective center alignment. This suggests that early visual areas make texture encoding more efficient by compressing neural manifolds (i.e. reducing radius and dimension) while decorrelating the neural responses to different textures (i.e. reducing center alignment). Interestingly, there is less compression of effective dimension for texture than for noise manifolds. We suspect that higher-order correlations in the texture manifolds may make them less compressible for early visual areas. Also, we find that the capacity and effective geometry of shuffled manifolds (i.e. shuffling neural responses with respect to stimuli) are quantitatively close to those of the unshuffled manifolds (Supplementary Figure S8a). This suggests that much of the structure in early visual responses is invariant to stimulus shuffling. Specifically, while the growth of capacity hints the improvement of coding efficiency from V1 to V2, the shuffling results reveal that the improvement is stimulus-independent (i.e., for different possible grouping of input stimuli) and not categorical (i.e., for the specific grouping of input stimuli according to the texture families). We speculate that such stimulus-independent structure may be relevant for flexible readout in downstream areas. Hence, we pose it as an interesting future research direction for further exploration.

**Figure S8:**
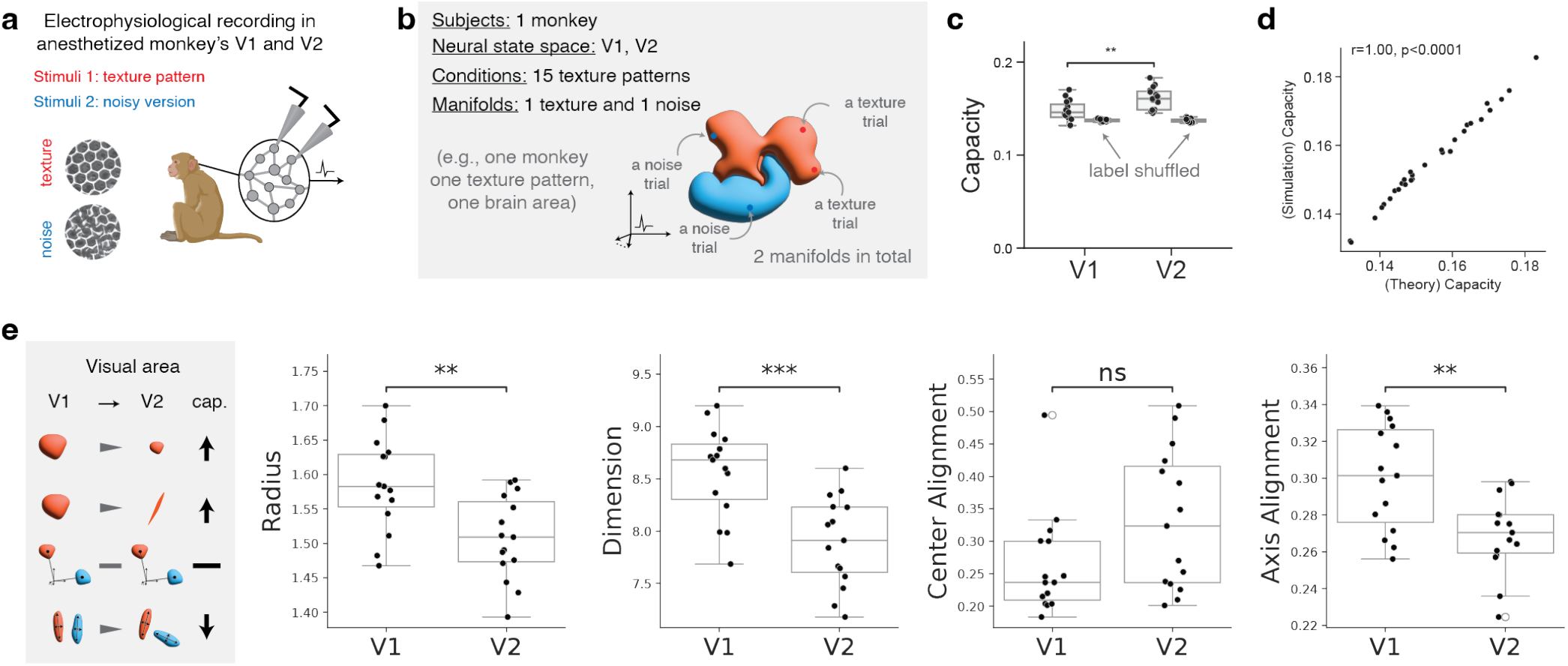
Supplementary figure for the GLUE analysis on a monkey early vision dataset [35].

Note that our analysis in the previous paragraph is different from the original analysis in [35] where the authors studied the difference in neural activation between a texture pattern and its noisy version. For completeness, in Supplementary Figure S8b we consider the analogous setting of analyzing the manifold capacity between a texture manifold and its noisy version. Concretely, for each texture family, we conduct GLUE analysis on two manifolds: a texture manifold formed by the collection of neural responses to the input stimuli from this texture family and the noise manifold formed by the collection of neural responses to the noisy version of input stimuli. We again see a growth of capacity along the visual hierarchy for most texture-noise pairs. Surprisingly, this time the center alignment increases from V1 to V2 (it decreased in the previous analysis); both effective radius and dimension decrease as expected whereas the axis and center-axis alignment increase. We interpret the result as hinting there are two opposing forces from V1 to V2 to the segregation of the texture and noisy manifold pairs: compression and categorization (i.e., aligning the two manifolds with the same lower order statistics). Further investigations into the changes of neural manifold organization in early visual areas require additional experimental design (e.g., subordinate texture families) and we pose it as an interesting future research direction.

#### S3.2 Monkey late vision dataset

In Figure 4a-c and Supplementary Figure S9 we analyze the dataset from [33] on a monkey vision task with electrophysiological recordings in V4 and IT. See Methods for more analysis details.

In the GLUE analysis, we see an increment of manifold capacity from pixel level to V4 and to IT in all variation levels (Figure 3a). The interpretation of capacity here corresponds to the encoding efficiency among the 8 basic categories (e.g., Animals, Cars, etc.), i.e., the level of categorization. From low variation to high variation level, there are also decreases of capacity for all brain areas as expected. The two experimental parameters (i.e., brain area and variation level) allow us to probe how manifold organization improves from V4 to IT. First, the increase of capacity from V4 to IT is a joint effect of decrease in effective radius and dimension as well as a reorganization of manifold alignments. In particular, from the comparison with synthetic examples (item S6c-d), we suspect that the center-axis correlations mainly drives the changes of the alignment measures and global scaling. Second, the decrease of capacity from low variation to high variation level is due to increase of effective radius and dimension as well as increase of effective center-axis alignment. Note that the geometric changes from low to high variation levels is exactly opposite the geometric changes from V4 to IT. Thus, we speculate that manifold compression and reorganization of center-axis alignments play critical roles in V4 and IT processing. Finally, we remark that manifold geometry at the subordinate category level (Supplementary Figure S9) closely resembles that at the basic level. We suggest delving into hierarchical manifold organization in V4 (e.g., [69]) and IT as an interesting future research direction.

#### S3.3 Human fMRI dataset

In Figure 4d-f and Supplementary Figure S10 we analyze the dataset from [34] on a human vision task with fMRI recordings in multiple ROIs. See Methods for more analysis details.

In the GLUE analysis, we see an increment of manifold capacity gain, i.e., the difference between the subordinate capacity of the category of interest (body part or face) and the control category (electronic devices), from early cortical areas (e.g., V1, V2) to late cortical areas (e.g., LOC, EBA, FFA) (Figure 3d). We interpret the capacity gain as an indication of functional selectivity to the category of interest. Specifically, this notion of selectivity captures the coding efficiency for distinguishing subordinate categories (e.g., boy face and girl face subordinate categories in the face category) in a brain region. In Figure 3e and Supplementary Figure S10, we see for every human subjects, EBA has the highest capacity gain for body part category and FFA has the highest capacity gain for face category. Meanwhile, we see no consistent gain in decoding accuracy in EBA and FFA for body part and face (Supplementary Figure S10). This finding significantly augments the conventional definition of brain area selectivity via activity level or decoding accuracy. Furthermore, the geometric measures also reveal a potential difference in the manifold organization of body part category and face category: while both categories have significant effective radius and dimension reduction in higher cortical areas, only the body part category exhibit strong axis alignment gain there. We pose it as an interesting future research direction to explore the definition of selectivity via manifold capacity in fMRI data.

#### S3.4 Monkey motor dataset

In Figure 6 and Supplementary Figure S11, we analyze the dataset from [44] on a monkey reaching task with electrophysiological recordings in PMd and M1. See Methods for more analysis details. In the following, we discuss four key observations and interpretations in the GLUE analysis: (i) task-relevant in-trial manifold dynamics, (ii) quantitative differences between premotor and primary motor area, (iii) manifold geometric signature for rapid adaptation, and (iv) changes in manifold geometry across sessions.

Recall that a trial consists of target presentation followed by a delayed period, then a go cue followed by movement onset (Figure 5a). We see a two-stage increase of manifold capacity (Figure 5b) throughout the trial, with capacity increasing faster during movement than during the delay period. The dynamics of capacity can be primarily explained by the compression of manifolds (i.e. a decrease of effective radius and dimension). Interestingly, there is a spike in both axis alignment and center-axis alignment between go cue and movement onset. This suggests rapid movement of manifolds along their center direction, as captured by the global scaling. Altogether, GLUE analysis provides a quantitative characterization of the preparatory and movement computations during monkey reaching: preparation focuses on manifold compression, while movement pushes the manifolds away along their center direction(Figure 5c).

Next, we see quantitative and qualitative differences in the in-trial manifold geometric dynamics between the two brain regions (Figure 5b). While the capacity of PMd and M1 are at the same level during preparatory period, the effective dimension of PMd is higher (this is consistent with previous findings using PCA [44]). In contrast, the effective radius exhibits an opposite trend where M1 has higher values. There are also significant differences in the effective alignment measures. During reach movement, the M1 capacity shoots higher than PMd capacity and is mainly driven by rapid compression of manifold radius. We speculate that the detailed geometric differences between PMd and M1 may lead to richer understanding in how motor signals are being transformed.

**Figure S9:**
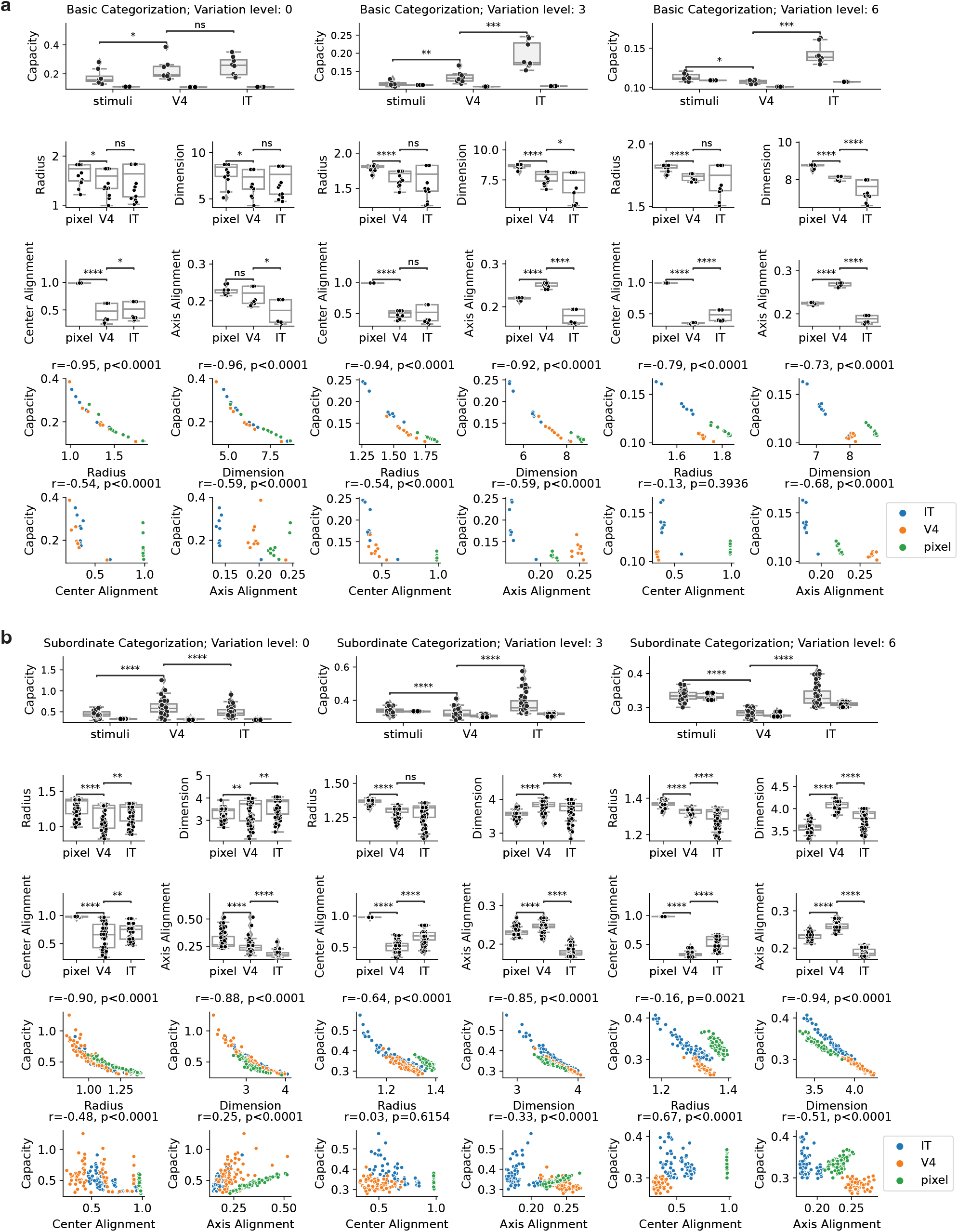
Supplementary figure for the GLUE analysis on a monkey late vision dataset [33].

**Figure S10:**
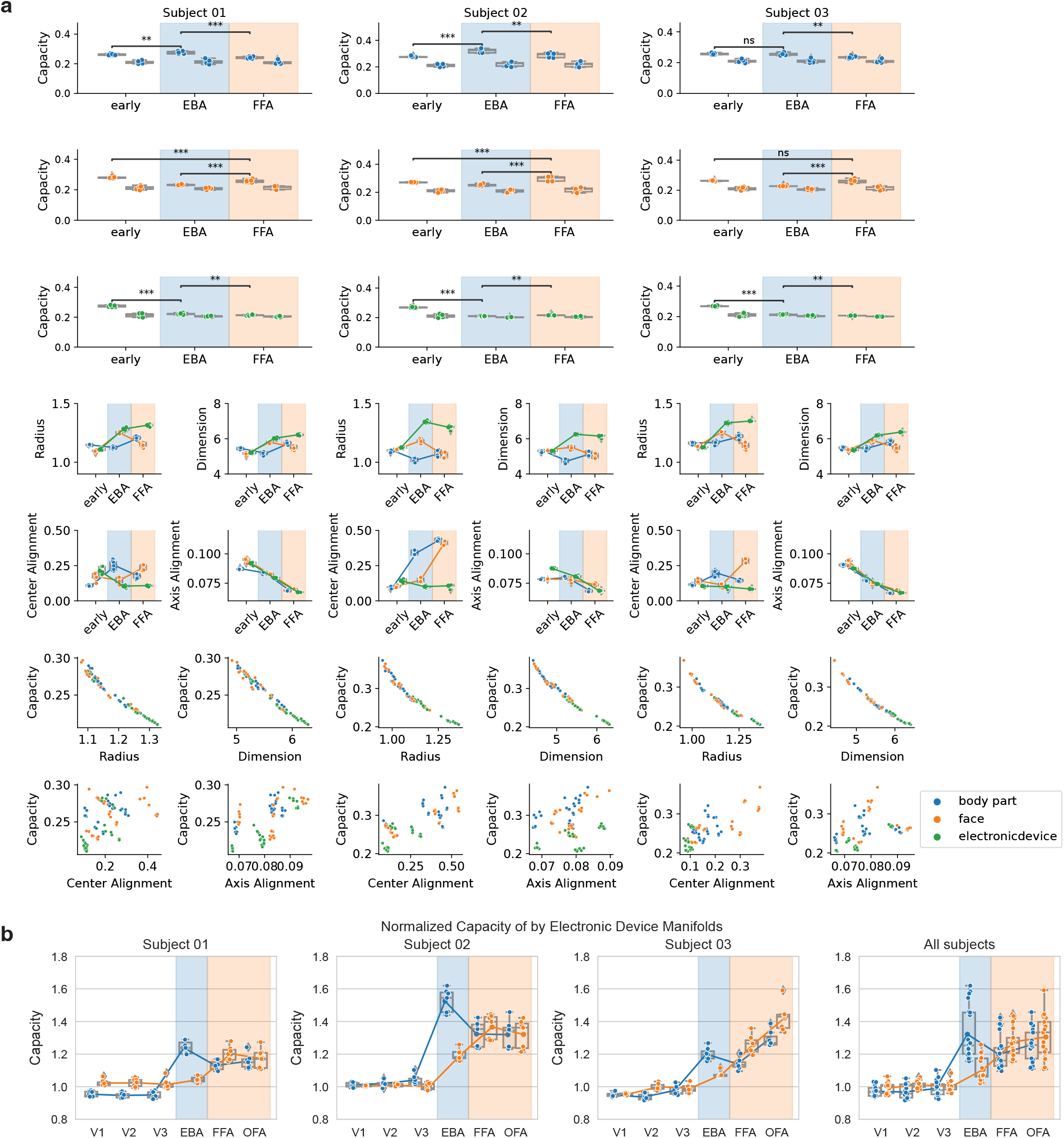
Supplementary figure for the GLUE analysis on a human fMRI dataset [34].

By analyzing manifolds from all session epochs (Supplementary Figure S11e), we see surprising geometric structures among the target-specific epoch manifolds. First, the capacity in the adaptation and washout epochs are generally higher than capacity in the baseline epoch, suggesting a global change in representational geometry during rapid learning in favor of encoding efficiency. Interestingly, the only place where the capacity of all these epochs being the same happens in M1 at the movement onset, hinting that the critical information for motor execution is well preserved across all epochs. Second, through analyzing the manifold alignment measures across all 40 = 5 × 8 (#epoch×#targets) manifolds, we see that the shift of neural activities in M1 (as partially suggested by previous work such as [44]) happens in the form of axis and center-axis alignment after movement onset. We suspect this is a consequence of an affine shift of the target-specific manifolds.

Finally, in Supplementary Figure S11 we investigate the difference of manifold geometry across sessions. Interestingly, we find that the strong axis and center-axis alignment within the adaptation epochs in M1 disappears in later sessions (Supplementary Figure S11a-c). Meanwhile, there is strong axis and center-axis alignment in PMd in later sessions (Supplementary Figure S11c). Note that this manifold signature does not appear in the visuomotor task. We do not have a functional or mechanical explanation for these cross-session manifold geometric dynamics and pose it as an interesting future research direction.

#### S3.5 Monkey perceptual decision dataset

In Supplementary Figure S12 we analyze the dataset from [47, 48] on a monkey perceptual decision task with electrophysiological recordings in prearcuate gyrus. See Methods for more analysis details.

Recall that a trial consists of the target presentation followed by a delayed period, then go cue followed by saccade (Supplementary Figure S12a). In Supplementary Figure S12b, we see a strong correlation between coherence level in the random dots experiment and the capacity increment throughout a trial: the capacity between two target-specific manifolds (i.e., left or right) grows higher when dot motion coherence is higher. The effective geometry further suggests that evidence integration during the task corresponds to a compression of the decision manifold (i.e., decrease of effective radius and dimension) and an increase of manifold center norm (as reflected by the growth of global scaling).

Next, we examine effective manifold alignment measures among the target-specific manifolds from all coherence levels (Supplementary Figure S12). First, the dynamics of the center alignment suggests a formation of curve-shaped organization, which is consistent with the observation reported in previous work [70]. Moreover, through the dynamics of the other alignment measures we find out that there are strong axis and center-axis alignment between manifolds with the same outcome label (i.e., left or right) and higher coherence level. Putting all the observations together, we propose the following potential mechanical description for that the evidence integration of prearcuate gyrus in a random dots experiment. There are two main forces for the dynamics of decision manifolds: (i) evidence condensation (as shown by a decrease in variability of each response manifold due to gradual evidence accumulation) and (ii) decision segregation (as shown by the rearrangement of the manifold alignment measures and curve-shaped organization, which is metabolically optimal as suggested by [70]). While the two forces both facilitate the separation of decision manifolds with different outcome label, they might come from different neural mechanisms and provide distinct functional consequences in more complicated tasks, e.g., multi-context decision-making. We leave this as an interesting future research direction.

#### S3.6 Mouse auditory learning and decision-making dataset

In Figure 5 and Supplementary Figure S13 we analyze the dataset from [38] on a mouse decision-making task with calcium imaging recordings in posterior parietal cortex. See Methods for more analysis details.

Recall that this is a mouse auditory discrimination task where a manifold corresponds to either high or low pitch stimuli in a session (Figure 5a). In the GLUE analysis, we first find that the capacity is positively correlated with the behavioral accuracy, which is consistent with previous work using decoding analysis [38]. Notice that the behavioral accuracy (as well as capacity and decoding accuracy) does not monotonously increase with the session days, i.e., generally the accuracy in both early and late session are lower. Surprisingly, we can explain the non-monotone phenomenon via the manifold geometry because, despite all individual geometric measures changing monotonically over time, different measures dominate the overall capacity during the early and late sessions and cause it to change non-monotonically. For example, effective radius, axis and center-axis alignment increase over time (except some small decrease in the first few sessions) while effective dimension and center alignment decrease over time. This explains the capacity and accuracy improvement throughout earlier sessions via compression of manifold dimension and decision segregation by a reduction in center alignment. In later sessions, once mice have learned the task (hence the dimension and center alignment stay low), the manifold radius increases and hurts capacity (perhaps mice are less attentive after learning). We speculate that there might be multiple neural mechanisms involved in this learning process (e.g., through different sub-populations such as excitatory and inhibitory neurons) and leave it as an interesting future research direction.

**Figure S11:**
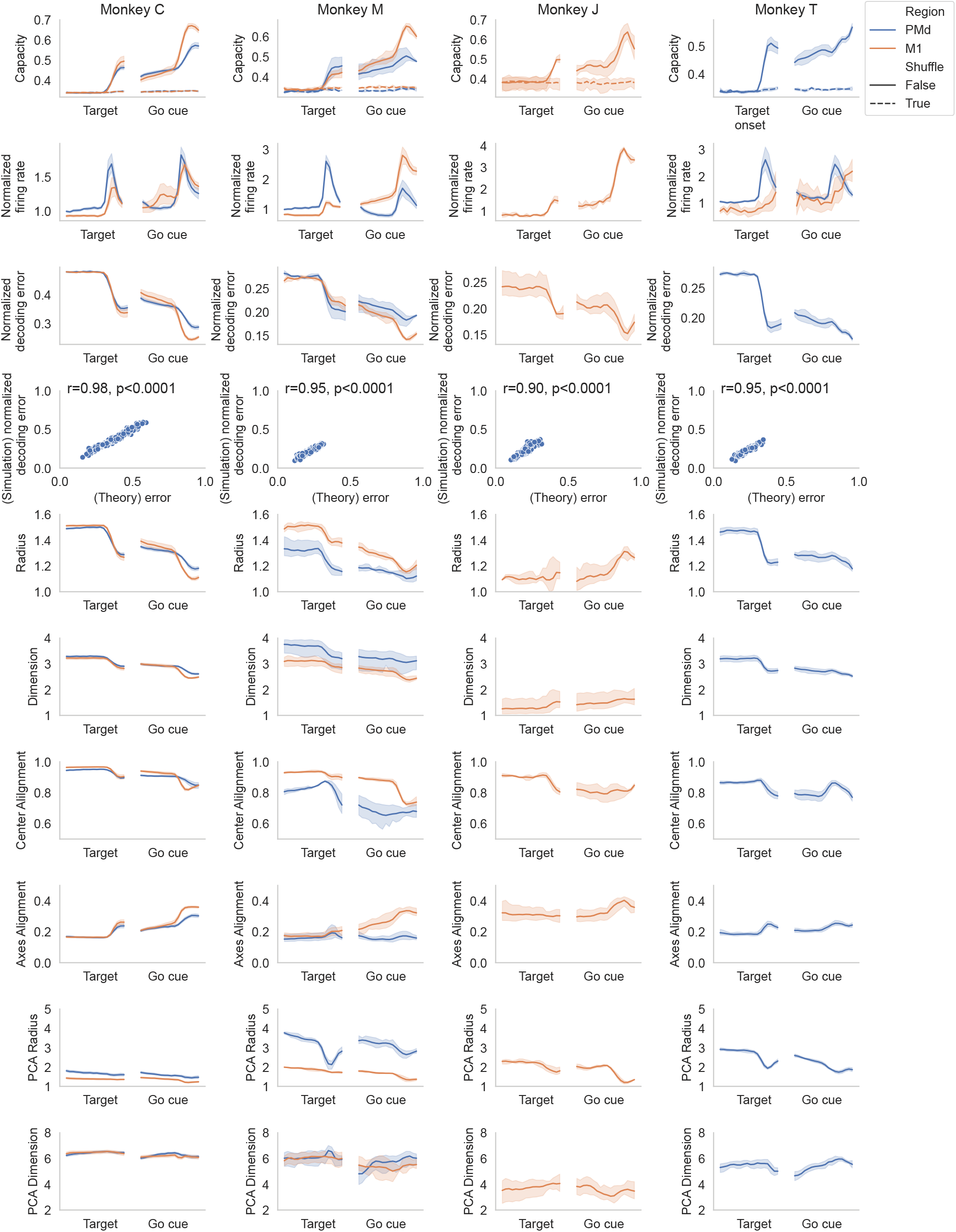
Supplementary figure for the GLUE analysis on a monkey reaching dataset [44].

**Figure S12:**
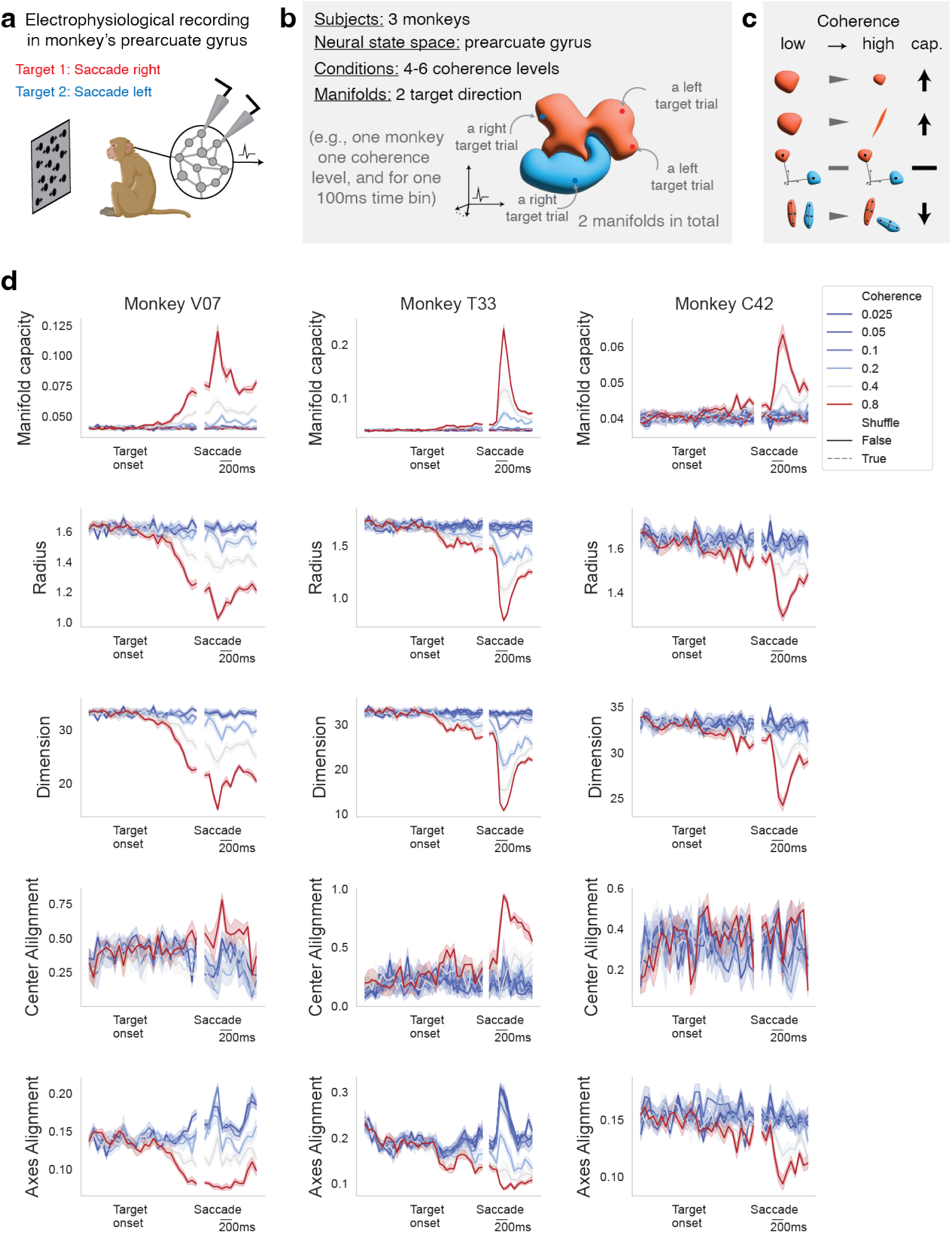
Supplementary figure for the GLUE analysis on a monkey decision-making dataset [47, 48].

#### S3.7 Mouse spatial memory dataset

In Supplementary Figure S14 we analyze the dataset from [42] on a mouse spatial memory task with calcium imaging recordings in hippocampus. See Methods for more analysis details.

Recall that this is a mouse VR linear track task where there are five environmental morph levels (Supplementary Figure S14a). There are two groups of mice: the frequent mice were exposed to all five morph levels during training while the rare mice were only exposed to the two extreme morph level. The GLUE analysis is conducted on the neural recordings from the testing stage where both types of mice were tested on all five morph levels. We first conduct a manifold alignment analysis and recover a consistent result to [42] on the representational similarity between morph levels (Supplementary Figure S14). Next, interestingly, we find that the capacity of the rare mice is higher than the capacity of the frequent mice (Supplementary Figure S14b). And the effective manifold geometry suggests that the difference in capacity is caused by the compression of effective radius and dimension. This information compression at the manifold level provides an evidence for population coding for spatial memory in mouse hippocampus. Further, we speculate these results hint at the possibility that training on extreme task conditions could lead to more compressed spatial representations, which facilitates better interpolation of the unseen environments. We pose it as an interesting future research direction to explore the interpolation and generalization in spatial memory at the neural manifold level.

#### S3.7 Artificial neural networks

In Supplementary Figure S15 we analyze the neural representations of pretrained ResNet and Vision Transformer on imagenet. See Supplementary Figure S15a-b and Methods for more analysis details.

We follow the setting and protocol in [28], which studied the manifold capacity and effective geometry of feedforward artificial neural networks. One downside of their analysis is that the authors removed the correlations between manifolds, and it was shown in [21] that this causes error in the estimation of capacity. The main purpose of this experiment here is to update their analysis with our new effective geometric measure that incorporate higher-order correlations between manifolds. In Supplementary Figure S15c, we found that the capacity formula in our theory perfectly matched with the capacity value estimated from the simulation process, while the formulas from previous work exhibit non-trivial gaps.

**Figure S13:**
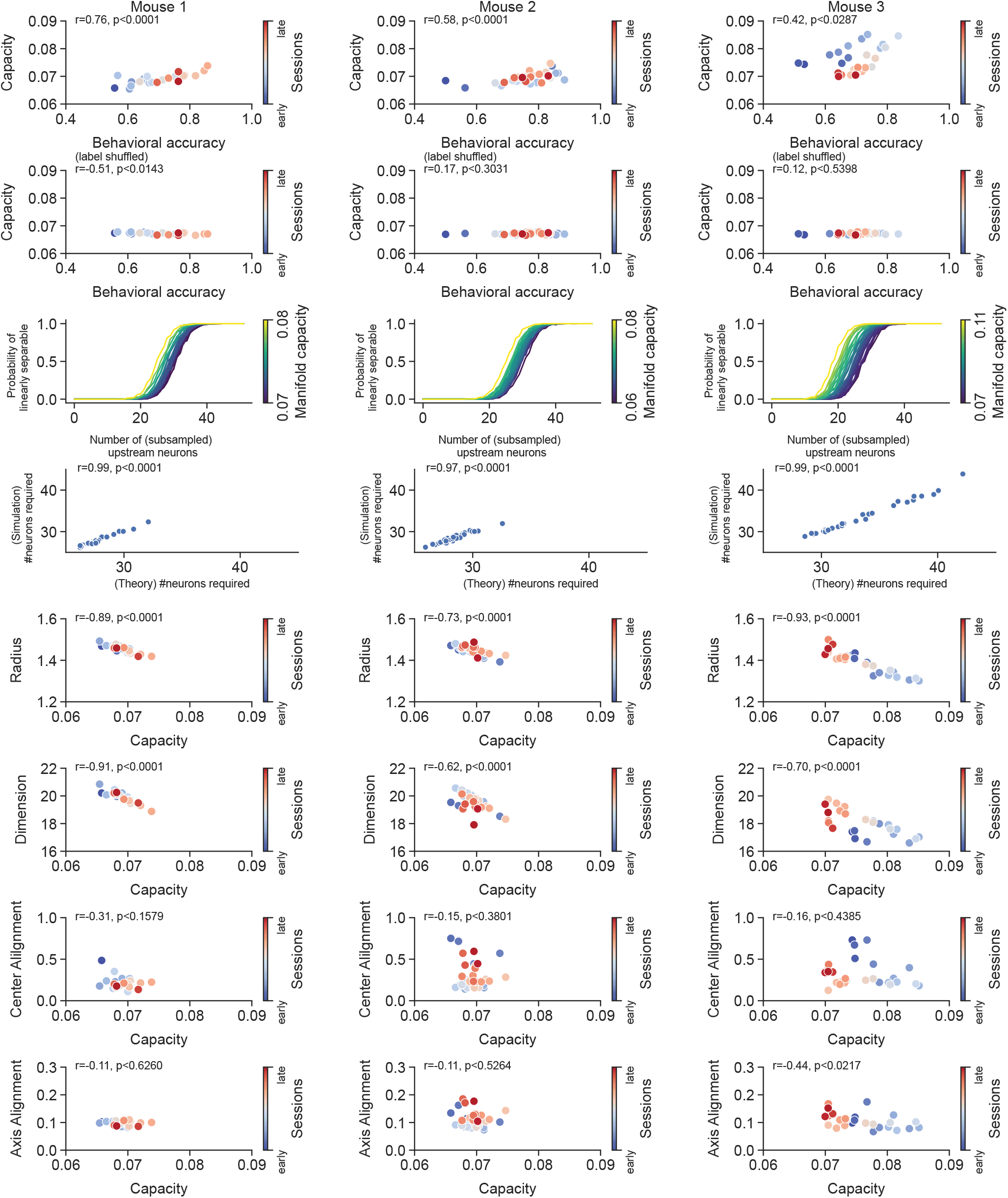
Supplementary figure for the GLUE analysis on a mouse auditory learning dataset [38].

**Figure S14:**
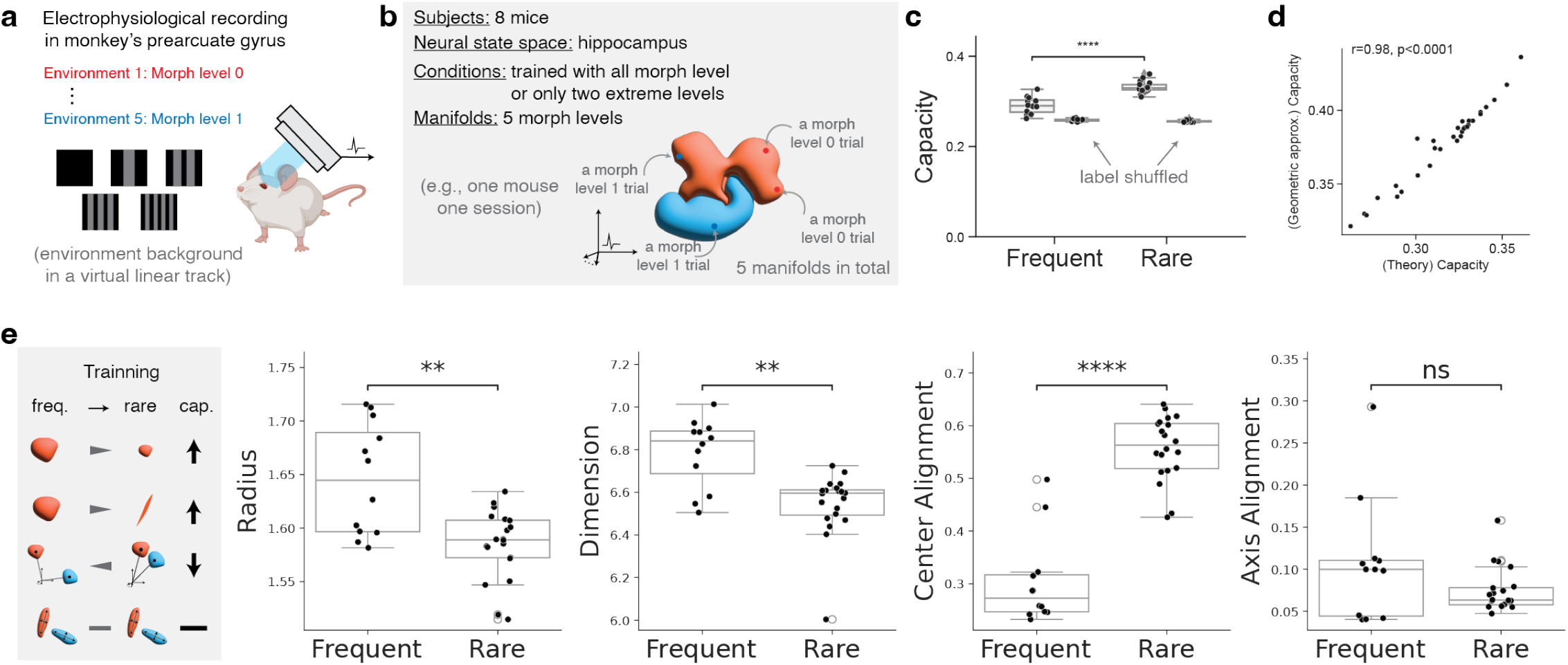
Supplementary figure for the GLUE analysis on a mouse spatial memory dataset [42].

**Figure S15:**
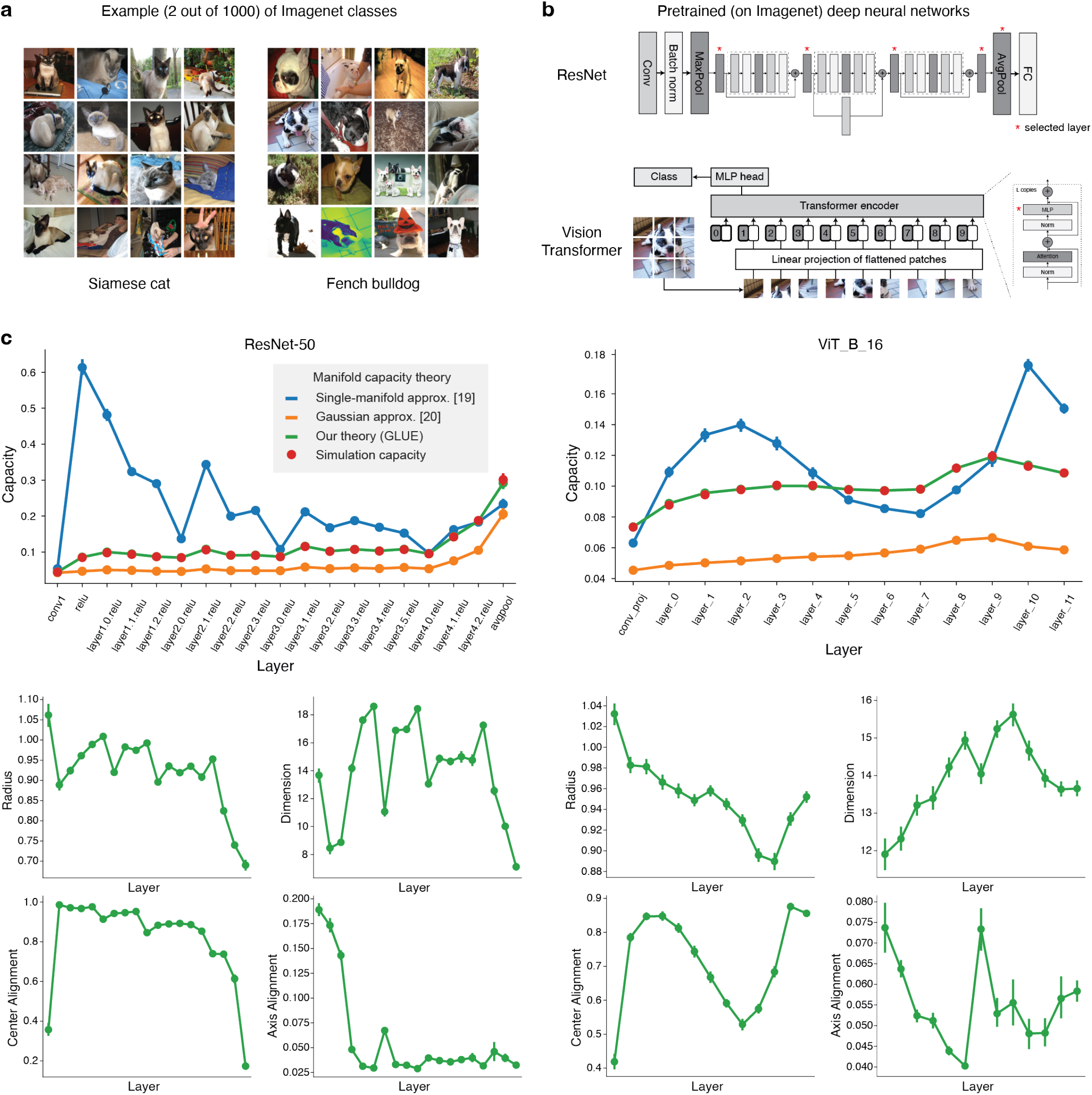
Supplementary figure for the GLUE analysis on artificial neural networks

## Footnotes

1 We term our geometric measures *effective* to highlight their analytical connection to computation, unlike conventional geometric analyses. See Supplementary Information S1 for more details. We will omit the term for brevity when the context is clear.

2 Linear classification/separation is the multidimensional equivalent to the simple thresholding operation used, for example, in single-neuron selectivity analysis (see Supplementary Figure S5).

3 Also known as *critical ratio* where critical point is a concept rooted in thermodynamics referring to the point where a phase transition happens, e.g., the liquid-vapor critical point is the temperature where water switches between liquid to gas.

4 More formally, separable with probability (over the randomness of random projection) at least 0.5.

5 To see this, notice that *p*_**y**,sim_(*N*) is monotone in *N* and takes value within 0 and 1. Namely, *p*_**y**,sim_(*N*) can be thought of as the cumulative probability function of a discrete random variable taking integer values. As a result, 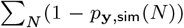 is simply the expected value of this random variable.

6 Also known as the “field” in [20].

7 Note that by the KKT condition, if the projection of **t** onto the cone of **s**(**t**) is zero, then **s**(**t**) = **0**. Thus, we have that proj_cone(**s**(**t**))_**t** = ⟨**s**(**t**), **t**⟩*/*∥**s**(**t**)∥_2_.

8 We remark that the statistical dimension of a convex cone is connected to the *intrinsic volume* of the cone as in Definition 2.2 and Proposition 5.12 of [68]

